# Localised admixture triggers parallel adaptation in a coral species trio on the Great Barrier Reef

**DOI:** 10.64898/2026.02.07.704610

**Authors:** Iva Popovic, Katharine E. Prata, Melissa S. Naugle, Ilha Byrne, Samantha M. Howitt, Zoe Meziere, Tom C. L. Bridge, Véronique J. L. Mocellin, Emily J. Howells, Line K. Bay, Nicolas Bierne, Cynthia Riginos

## Abstract

Hybridisation can shape evolutionary trajectories and fuel rapid adaptation. Yet, whether adaptive introgression repeatedly reconfigures genomes in predictable ways remains largely unknown. Here, we show that three-way admixture drives repeatable parallel differentiation in reef-building corals in the *Acropora hyacinthus* species complex. Adaptive introgression is confined to the southern Great Barrier Reef, where admixture timing coincides with a recent breakdown in species barriers despite limited historical gene flow. Genome-wide introgression is characterised by numerous shared minor-parent ancestry peaks that are enriched for stress-response genes and under positive selection for parallel adaptive admixture. Our results demonstrate that introgressive hybridisation can be geographically restricted between sympatric marine species and can promote repeated rapid adaptation that transcends species boundaries.

## Introduction

Hybridisation profoundly shapes genome evolution, bringing together new combinations of genes from distinct lineages that are subsequently winnowed by natural selection (*1*). Despite the prevalence of hybridisation across the tree of life (*2*), when and how hybridisation can fuel adaptive evolutionary change is largely unknown. Striking examples of strong positive selection on introgressed gene regions involving immunity (*3*), pigmentation (*4*) and chemical resistance (*5*) indicate that hybridisation can promote rapid adaptation and phenotypic matching to environments (*6*). Yet, whether such examples constitute rare episodes of adaptive capture of large-effect genes that leave minor modifications to species’ genomes or whether adaptive introgression can predictably restructure genomes remains a critical gap in our understanding of evolutionary change (*7, 8*).

Comparisons among hybrid zones provide a powerful framework to identify the fate of parental alleles, where parallel genome evolution in admixed populations derived from the same parental species provides compelling evidence for selection (*8*). Such comparative studies have identified pervasive negative selection against foreign ancestries, where introgressed alleles descended from the minor-parent – the lineage contributing the smaller fraction of the hybrid genome – are repeatedly purged, particularly in regions of low recombination (*9*) or high deleterious load (*10–13*). In contrast, other genomic regions move across species boundaries, producing a genomic patchwork governed by whether loci are neutral or selectively favoured, where in adaptive introgression may also generate fine-scale repeatability across distinct lineages (*4*) or replicated hybrid populations (*14*). Conventional comparative hybrid zone studies typically examine replicated hybrid populations either descended from the same parental species pair (*9, 11, 12, 14–16*) or from different species pairs but in distinct geographic locations of secondary contact (*16–18*). With this framework, repeatable ancestry sorting can be ascribed to endogenous selection arising from interactions among parental genomes and shared genomic architectures (*9, 18*), whereas differing genomic outcomes may reflect contrasting local environmental selection (*19, 20*) or some other source of contingency (*10, 11*).

Coincident admixture among three or more sympatric parental species, although rare (*21, 22*), provides an alternative conception of the comparative hybrid zone framework, where locus-specific outcomes can be contrasted across different parental genomic backgrounds (i.e., major-parent) that coexist in the same ecological setting. Under this scenario, underrepresentation of minor-parent alleles at the same loci across multiple major-parent backgrounds would provide evidence for parallel negative selection (*18*). Conversely, repeated introgression of the same minor-parent alleles into multiple major-parent genomes would constitute strong evidence for parallel positive selection. Importantly, multi-species admixture opens the possibility that genes locally adapted to the shared geographic location are repeatedly captured and reused across multiple species’ genomic backgrounds, manifesting as parallel adaptive introgression. Although not yet described, such repeated allele reuse would provide evidence that introgressive hybridisation can contribute to deterministic adaptive genetic change within multiple sympatric lineages.

Here, we document such parallel adaptive change driven by adaptive introgressive hybridisation in a coral species trio. Although hybridisation in corals is long recognised (*23, 24*), genomic studies have only recently demonstrated that ongoing gene flow is common (*25–27*) and highly heterogeneous across the genomes of closely related taxa (*28*). Hybridisation in ecologically-important *Acropora* corals has been proposed to aid adaptive responses to climate-driven selection (*29*), promote diversification (*30*) and enhance thermal tolerance (*31*). Yet, no previous study has tested whether introgression between closely related coral species can be adaptive and, consequently, predictably restructure their genomes. In this study, we examine how introgressive hybridisation reshapes coral genomes to generate parallel adaptive outcomes. Focusing on naturally occurring three-way interspecific admixture in the ‘*Acropora hyacinthus* species complex’ (*24, 32*), we test the repeatability of introgressed local ancestry across divergent hybridising species. By integrating signatures of parallel adaptive introgression with estimates of admixture timing and gene flow histories, we assess the permeability of species boundaries and provide evidence that gene flow has repeatedly transferred beneficial alleles among species through positive selection on admixed variation.

### Three-way admixture is localised to the southern Great Barrier Reef

We used whole genome data from 823 coral colonies within the *Acropora hyacinthus* species complex (‘AH’ hereafter) and 11 *Acropora millepora* colonies sampled across the Great Barrier Reef (GBR: Figure 1A, Figure 1B; (*33*)) to investigate population structure and hybridisation among AH species. AH colonies comprised four genetically distinct and sympatric species, inferred by Principal Component Analysis (PCA) and Bayesian clustering in ANGSD (*34, 35*) (Figure 1C), identified as *A. tersa* (*32*), *A. pectinata* (*36*), Taxon 3 (i.e., unnamed taxon), and *A. hyacinthus* (*35, 37*). Each taxon was genetically distinct across the GBR, with admixture among three AH species restricted to their southern range edge. For *A. pectinata*, Taxon 3 and *A. hyacinthu*s (Figure 1D), but not *A. tersa*, there was a clear genetic separation between some southern GBR colonies (Figure 1E) and central-northern populations. Strikingly, these genetically distinct southern populations were entirely admixed, with colonies deriving an average of 3.5%-14.8% of their genomes from the two other AH species (Figure 1E). Other southern colonies were free from admixture and genetically similar to parental-type populations found GBR-wide (Figure 1E). This coexistence of admixed and non-admixed southern colonies suggests partial reproductive isolation and that selection may favor admixed genotypes under certain ecological contexts. By contrast, early-generation hybrids (n=6; admixture proportions ∼40-70%) were almost exclusively found in the central and northern GBR, consistent with selection against further backcrossing maintaining reproductive isolation.

**Figure 1.**
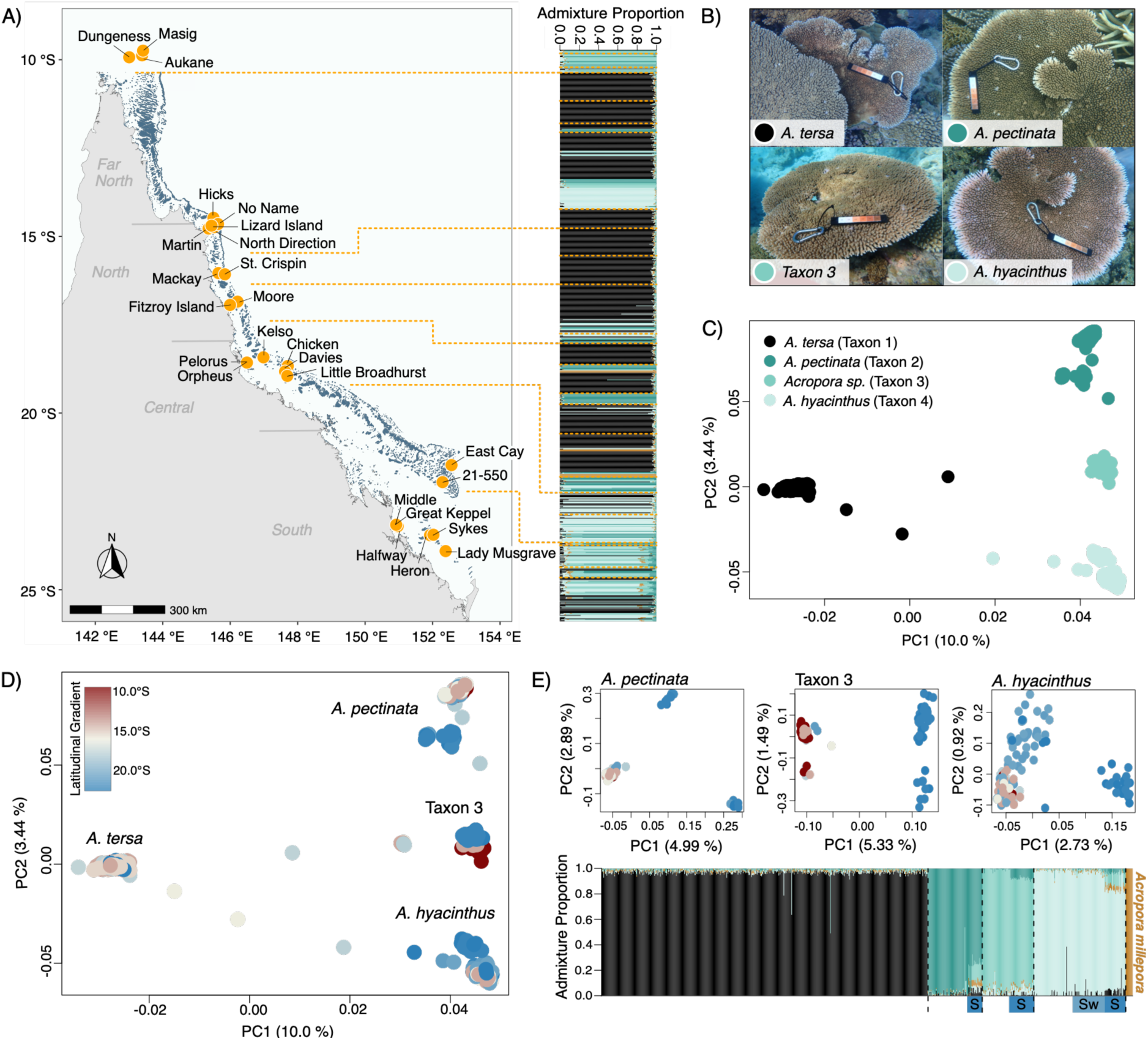
Sampling locations and population genetic structure in the *Acropora hyacinthus* species complex (AH). **(A)** Sampling across 26 reefs across four regions of the Great Barrier Reef (GBR), Australia: Far North (> -15.5°S), North (-18°S to -15.5°S), Central (-21.5°S to -18°S) and South (< - 21.5°S). **Right panel:** Bayesian clustering analysis showing four genetically distinct AH species occurring sympatrically across the GBR with dashed orange lines separating colonies from different reefs; **(B)** Colony photographs of the four AH species; **(C)** Principal component analysis (PCA) on 92,669 unlinked loci showing genetic separation among the four AH species with PC1 and PC2 explaining 10.0% and 3.44% of variation respectively; **(D)** PCA with colonies coloured by latitude; **(E)** Species-specific PCAs for *A. pectinata*, Taxon 3, and *A. hyacinthus* showing genetic separation on PC1 of southern populations from the Capricorn Bunker (S) and Swains (Sw) reef groups. Some colonies from southern reefs (blue) were genetically similar to parental-type species found GBR-wide. **Bottom panel:** NGSAdmix plot showing that differentiated southern populations are comprised entirely of admixed individuals deriving an average of 3.5%-14.8% of their genomes from the two other AH species. Six putatively recent hybrids with mixed admixture proportions (40-70%) were largely present in central-northern populations.

Consistent with three-way admixture (Figure S1), southern-admixed populations of *A. pectinata*, Taxon 3 and *A. hyacinthus* showed close genetic affinities (Figure 2A), implicating shared genetic ancestries among all three parental species. Three-way admixture was evidenced by significant migration edges connecting southern-admixed populations in TreeMix (*38*) (Figure S2). Genome-wide D-statistics (*39, 40*) also supported localised three-way introgression with significant gene flow among southern-admixed populations (Figure 2B; Table S1). In contrast, contemporary gene flow between species was largely absent in northern and central regions, excepting between *A. pectinata* and Taxon 3 (Figure 2B; Table S1). These findings provide strong evidence for region-specific reproductive isolation among broadly sympatric coral species and raise the question as to why and how species barriers are maintained in only some parts of the species’ range.

**Figure 2.**
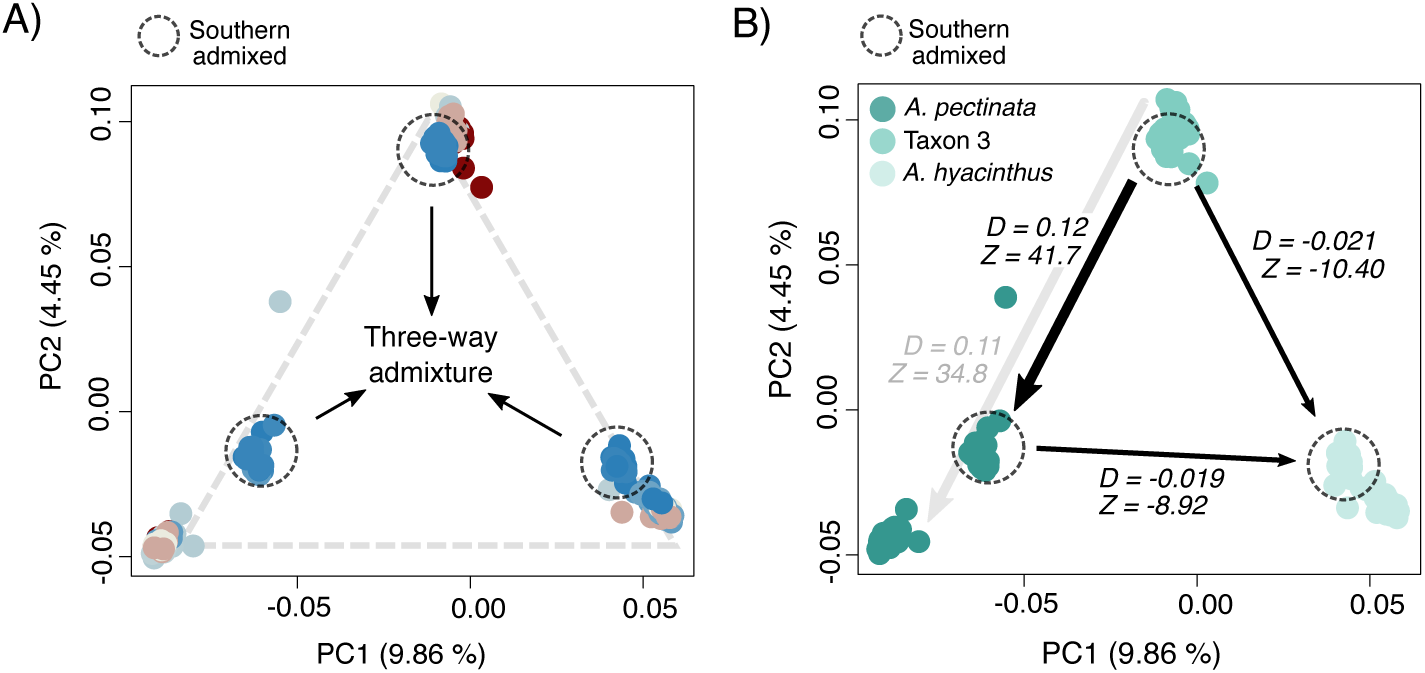
Genomic evidence for localised three-way admixture in three focal species of the *Acropora hyacinthus* species complex (AH) in the southern Great Barrier Reef. **(A)** Principal component analysis showing closer genetic affinities among *A. pectinata*, Taxon 3, and *A. hyacinthus* colonies from southern reefs (blue), consistent with expectations for three-way admixture (Figure S1). **(B)** Evidence for variable introgression in different GBR regions based on genome-wide D-statistics. In the southern GBR, all comparisons among southern-admixed populations (*A. pectinata*, Taxon 3, *A. hyacinthus*) were significant (black arrows; absolute *D*-statistic range 0.01-0.12; Z-score > abs(4)), supporting three-way introgression. In the central-northern GBR, only comparisons between *A. pectinata* and Taxon 3 were significant (grey arrow; absolute *D*-statistic = 0.10).

Given previous suggestions that introgression from *A. millepora* into one AH species in American Samoa (*31*) may have played a role in adaptive evolution, we tested whether population structure and genetic similarities observed among southern-admixed AH species could be driven by shared past *A. millepora* introgression. Genome-wide D-statistics revealed evidence for *A. millepora* introgression with *A. pectinata*, Taxon 3 and *A. hyacinthus* throughout their GBR distributions (Table S2) (*35*). To locate introgressed genomic regions putatively involved in adaptive introgression, we calculated *f_d_*, the fraction of the genome showing excess admixture in 50 kb windows (Figure S3 (*41*)). Several introgression hotspots (top 1% of *f_d_* outliers) spanning up to 400 kb were evident on several chromosomes (e.g., Figure S4), including a region corresponding to the HES1 locus previously speculated to be involved in *A. millepora* introgression (*31*). These results provide empirical evidence that AH species may harbour large chromosomal regions originating from *A. millepora* (*31*). However, because genome-wide introgression patterns were concordant among AH species and across GBR regions (Spearman’s *ρ* ≥ 0.50, *p* < 3.24 x 10^-110^; Figure S5), we inferred that *A. millepora* introgression likely largely predates lineage divergences among AH species and does not contribute to present-day genetic segregation observed in southern-admixed AH populations.

### Extensive admixture coincides with a recent breakdown of species barriers against a backdrop of limited historical gene flow

Species divergences among reef-building Scleractinian corals predate modern-day GBR reef configurations, which formed ∼7,600-10,000 years ago following rapid sea-level changes after the Last Glacial Maximum (LGM: (*42–44*)). Consequently, present-day species distributions and genetic diversity likely reflect dynamic cycles of isolation and contact during glacial-interglacial periods, resulting in broadly sympatric and partially reproductively isolated taxa with ample opportunities for genetic exchange. Episodes of enhanced gene flow can further promote rapid evolution when demographic structure is unstable (*45*), particularly under recent colonisation or ecological disturbance (*46, 47*), and may facilitate evolutionary rescue by introducing adaptive variation (*5, 19*) or masking weakly deleterious alleles in small receiving populations (*11*). To determine whether three-way admixture among AH species reflects ongoing or episodically enhanced gene flow, and whether it postdates the LGM involving all three species simultaneously, we combined local ancestry inference in southern-admixed AH populations with demographic modelling to reconstruct the evolutionary history of gene flow (*35*). By drawing on non-admixed parental-type populations from the central-northern GBR, demographic inferences allowed us to resolve deeper historical patterns of interspecific gene flow among three focal AH species and contrast patterns with those observed with *A. tersa* and *A. millepora*.

We first jointly estimated the maximum likelihood admixture timing and genome-wide local ancestry patterns by applying hidden Markov models to more than 130,000 ancestry-informative sites in Ancestry-HMM (*35, 48*). We modelled three-way admixture among southern-admixed populations (*A. pectinata*, Taxon 3 and *A. hyacinthus*) using ‘two pulse’ admixture models (*35, 49*) for each ancestral (major-parent) population and inferred posterior probabilities for ancestry states with central-northern non-admixed populations as parental-type references. Across populations and admixture pulses, analyses consistently inferred similar and recent admixture times (1,000-2,000 generations ago; Figure S6), suggesting that southern-admixed populations formed contemporaneously with post-LGM reef reconfigurations ((*35, 43*); assuming a 3-year generation time). Demographic modelling (*50*) focused on non-admixed populations provided concordant evidence for a recent, coordinated breakdown in genomic barriers to gene flow among *A. pectinata*, Taxon 3 and *A. hyacinthus* ∼2,000-4,000 generations ago (Figure 3; Table S3-A; Table S3-B). We observed patterns consistent with recent elevated gene flow affecting a much larger proportion of the genome (up to 84%; *P_E2_* = 0.16-0.40; Figure 3; Table S3-B). This contrasts to lower migration rates and strong genomic barriers affecting most of the genome in the first time period (>66%; *P_E1_* = 0.54-0.66; Table S3B) following their divergence ∼300,000 generations ago.

**Figure 3.**
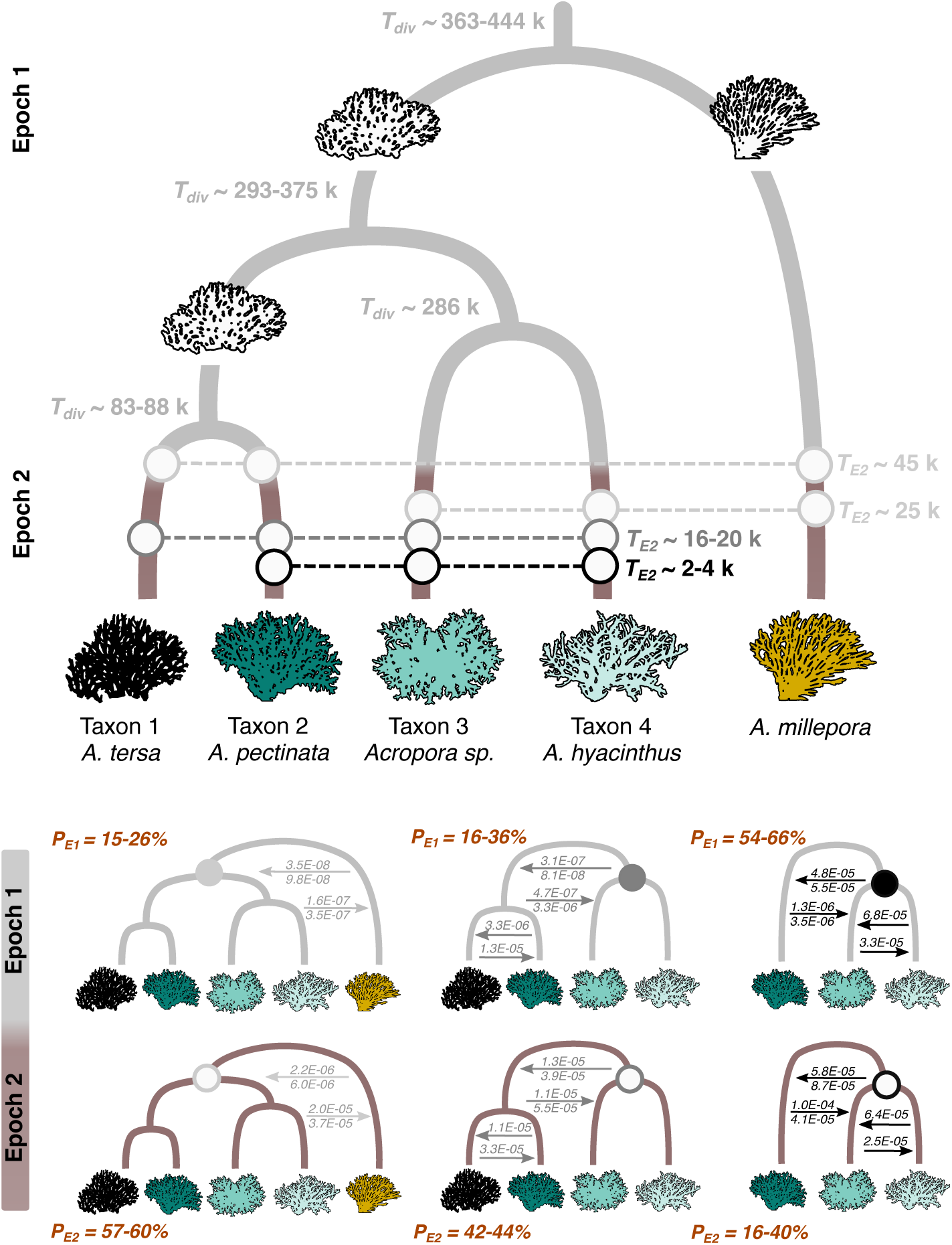
Demographic history of divergence and gene flow in the *Acropora hyacinthus* species complex (AH) based on optimised models inferred with *dadi.* **Top Panel:** Demographic tree shows divergence times (*T_div_*) in units of generations where white corals represent ancestral lineages. The timing of the second epoch (*T_E2_*) among paired comparisons is in units of generations and denoted with dashed lines and circles connecting lineages in paried model comparisons. **Lower panel:** Demographic trees showing paired model comparisons with *A. millepora* (left), *A. tersa* (middle) and comparisons among three focal AH lineages (right), where arrows indicate the migration rate as the proportion of migrants per generation (*m*). *P_E1_* and *P_E2_* denote the proportions of the genome experiencing no genetic exchange in the first and second time periods, respectively, where the width of the orange rectangles depicts a build-up of genomic barriers to gene flow with *A. millepora* (left) and *A. tersa* (middle), and a breakdown of species barriers among three focal AH lineages in the second epoch. Coral images were sourced from Phylopic (http://phylopic.org).

Elevated and recent genetic exchange among three focal AH species (Figure 3) alongside substantial contemporary admixture in the southern GBR (Figure 2) stands in sharp contrast to low historical gene flow levels with *A. tersa* and *A. millepora*. Rather than indicating a breakdown of species boundaries, demographic models involving *A. tersa* and *A. millepora* revealed increased genomic barriers at the onset of the second time period (∼20,000-45,000 generations ago), with a larger fraction of the genome experiencing no gene flow (*P_E2_* = 0.42-0.60; Figure 3; Table S3-A; Table S3-B). This shift points to the establishment of barriers to gene exchange, with subsequent introgression restricted to a small subset of loci consistent with few genomic regions showing evidence of *A. millepora* introgression based on *f_d_* outliers (Figure S3).

By leveraging comparisons among closely related, co-occurring coral species, our results demonstrate that present-day genetic diversity in GBR *Acropora* reflects complex histories of shifting species permeability and dynamic phases of interspecific gene flow. Three focal AH species remain well-isolated in the central and northern GBR with a history of low historical introgression, but where recent, enhanced gene flow with greater genomic permeability aligns with the timing of localised three-way admixture at southern reefs. These findings suggest that either historical admixed populations among modern central-northern GBR populations were ephemeral or that recent novel environmental or ecological conditions during the early-mid Holocene may have favoured admixed genotypes in the southern GBR range edges, potentially promoting adaptation through admixture arising ∼1,000-2,000 generations ago.

### Parallel genome-wide hybridisation outcomes at southern reefs

Having established that *A. pectinata*, Taxon 3 and *A. hyacinthus* remained largely isolated for at least ∼300,000 generations, followed the recent onset of contemporary admixture at southern reefs, we next test whether parallel adaptive introgression has produced similar genomic outcomes across major-parent backgrounds. Indeed, we find that southern-admixed AH populations diverged in parallel from parental-type central-northern populations (Figure S7-A). PCA-based selection scans (*51*) revealed multiple genomic regions among the top 0.01% of outliers differentiating southern-admixed and non-admixed central-northern populations (Figure S7-B). This unique scenario of three-way admixture and parallel differentiation among sympatric species allows comparisons of locus-specific outcomes across distinct major-parent genomic backgrounds that coexist in the same ecological setting.

To test whether selection has led to parallel adaptive introgression, we inferred local ancestry states across three possible ancestries (Figure S8) and quantified genome-wide ancestry correlations between southern-admixed populations. Across all pairwise comparisons, minor-parent ancestry in one major-parent background strongly predicted the same minor-parent ancestry in the other major-parent background, indicating that the same gene regions have introgressed into both non-matching parental genomes (Figure 4). These correlations were consistently positive across window sizes (20-200 kb; Spearman’s *ρ* range 0.46-0.65; *p* < 1 x 10^-16^; Figure 4; Figure S9), and persisted across ancestry-informative sites (Spearman’s *ρ* range 0.51-0.66; *p* < 1 x 10^-16^; Figure S10) and across chromosomes (Figure S11), indicating that convergent selection has shaped both broad and fine-scale genomic ancestry patterns likely under varying recombination levels (*18*). Hundreds of ancestry-informative sites (1.3%; Figure S12) contributed to shared minor-parent ancestry ‘peaks’ defined as regions where the same minor-parent ancestry has been maintained at high proportions exceeding 50% in two species. Relative to chance, admixed genomes were ∼5 times more enriched for shared minor-parent ancestry peaks (*p* < 4.9 x 10^-4^; Figure S13). Moreover, >95% of these shared minor-parent peaks corresponded to genomic regions where the same allele also reached high proportions (>50%) in the major-parent background (Figure S12), suggesting selective sweeps of the same ancestry in all three southern-admixed populations (e.g., Figure 5).

**Figure 4.**
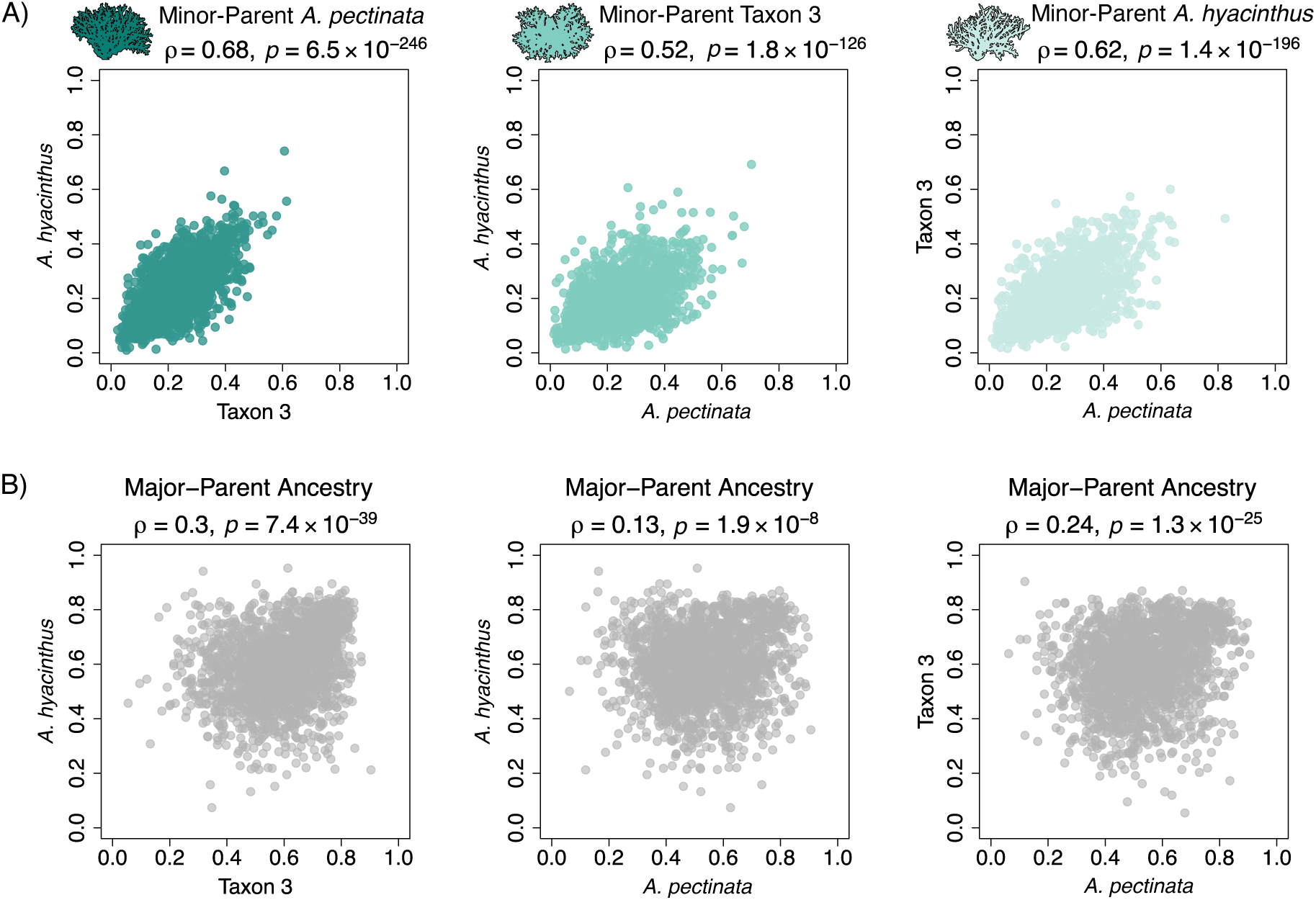
Genome-wide local ancestry correlations in three southern-admixed populations of the *Acropora hyacinthus* species complex (AH). **(A)** Significant positive correlations in minor-parent ancestry proportions averaged across 200 kb windows in divergent major-parent genomic backgrounds (Spearman’s *ρ* range 0.52-0.68; *p* < 1.8 x 10^-126^). **(B)** Major-parent ancestries averaged across 200 kb windows were weakly correlated among AH species (Spearman’s *ρ* range 0.3-0.24, *p* < 1.9 x 10^-8^).

**Figure 5.**
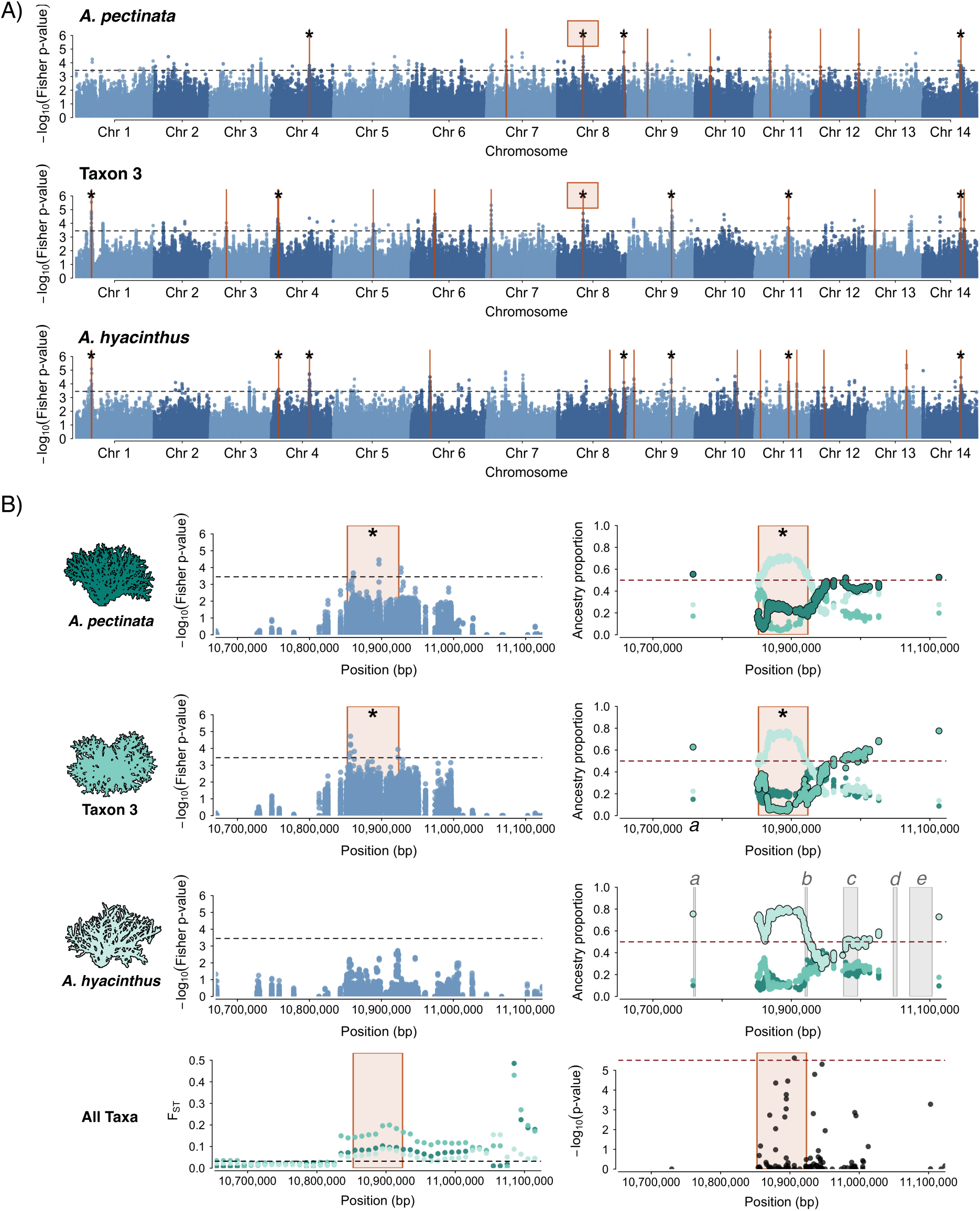
Evidence for positive selection on admixed variation in three hybridising species of the *Acropora hyacinthus* species complex (AH) in the southern Great Barrier Reef. **(A)** Genome-wide Fisher p-values, where the black horizonal lines indicate the Bonferroni significance threshold (*p* < 0.00035) and orange vertical lines indicate 29 candidate genomic regions for parallel adaptive admixture where signatures of positive selection coincide with shared minor-parent ancestry islands. Asterisks indicate candidate genomic regions with evidence for positive selection in two or more species. **(B)** Local signatures of parallel adaptive admixture on a focal Chromosome 8 region (indicated with an orange rectangle in (A) and (B)), where *A. hyacinthus* ancestry has risen to high frequencies across numerous SNPs in admixed populations. Species-specific ancestry is denoted by coral icon colors and major-parent ancestry is outlined in black. This region also coincides with parallel F_ST_ deviations between southern-admixed and non-admixed central-north populations (bottom left), including a SNP identified as a genomic differentiation outlier using a PCA-based selection scan (bottom right). This region coincides with functional annotation for an enoyl-CoA hydratase-isomerase family gene (*b*) involved in peroxisomal fatty-acid metabolism and is in close proximity to a number of genes with diverse functions (*a, c, d, e*; gene functions in Table S5).

Shared genomic architectures and neutral gene flow can generate local ancestry correlations under some circumstances (*35*) and during different phases of evolution following admixture (*8*). However, if selection against shared, large-effect genetic incompatibilities were the primary force shaping introgression, we would expect strong, concordant major-parent ancestry correlations across hybridising taxa, reflecting consistent removal of minor-parent foreign ancestry in the same genomic locations (*9, 18*). Instead, major-parent ancestries were only weakly correlated (Spearman’s *ρ* range - 0.13-0.3, *p* < 1.9 x 10^-8^; Figure 4; Figure S14; Figure S15; Figure S16), suggesting that genetic incompatibilities and linked selection only partly explain the observed patterns. Thus, in contrast to systems where minor-parent ancestry correlations between hybrid populations descended from two parental species can be interpreted as largely reflecting negative selection against foreign ancestry (*9, 12*), our three-way admixture framework reveals concordant peaks of the same minor-parent ancestry in the absence of clear genomic depletions or ‘deserts’ of major-parent alleles. Instead, observed patterns arise from selection favouring repeated introgression genome-wide within divergent major-parent genomes that otherwise resist introgression across most of their geographical range in the GBR.

Although we could not investigate how recombination shapes local ancestry patterns, the marked excess of shared minor-parent ancestry peaks between species cannot be accounted for without invoking natural selection on introgressed variation. This parallelism is consistent across all three possible minor-parent ancestries (Figure 4A), where different introgressed ancestries are repeatedly favoured across the genome, producing a ‘mix-and-match’ genomic mosaic shaped by adaptive introgression. Supporting this interpretation, shared minor-parent ancestry peaks were strongly enriched for *response-to-stress* associated Gene Ontology terms as the top functional category (*p* ≤ 0.0001; Figure S17-B), whereas private (non-shared) peaks were enriched for *reproduction-*related functions (*p* < 0.0001; Figure S17-A). These results indicate that repeatedly introgressed variation has experienced distinct selection pressures compared to lineage-specific variation.

### Shared minor-parent ancestry islands are numerous and under positive selection

Given strong evidence for genome-wide parallelism under three-way admixture, we hypothesised that positive selection may underlie repeated introgression of the same minor-parent alleles across species, if beneficial variants from either parent repeatedly sweep in distinct hybrid populations and remain advantageous after admixture (*52, 53*). Under this scenario, recurrent selection would maintain introgressed adaptive alleles, elevating allele frequencies and increasing local minor-parent ancestry proportions around putatively selected loci relative to the genome-wide mean ancestry. To test this hypothesis, we examined whether post-admixture positive selection promotes such repeatability by examining diversity patterns expected under admixture and selection to identify genomic regions involved in parallel adaptive introgression.

Two neutrality statistics, *F_adm_* and *LAD*, have substantial power to detect positive selection in admixed genomes and loci under selection during ‘adaptive admixture’ (*35, 52*) –where beneficial alleles in parental species remain advantageous and selectively favoured after introgression in hybrid populations (i.e., *scenario 1* in (*52*)). This contrasts to allelic neutrality in parental genomes followed by post-admixture positive selection (i.e., *scenario 3* in (*52*)). Evidence of adaptive admixture would indicate that predictable hybridisation outcomes arise through selection favouring the same introgressed variants across divergent genomes. *F_adm_* is the squared difference between observed and expected allele frequencies (based on source parental diversity), and *LAD* is the per-window local ancestry deviation from the genome-wide mean (*35*). By combining *F_adm_* and *LAD* statistics into empirical p-values using Fisher’s method as a composite test of post-admixture positive selection (*52*), we detected significant signals of adaptive admixture in southern-admixed AH populations and evaluated how often these signals overlapped within shared minor-parent ancestry ‘islands’ (clusters of five or more shared minor-parent ancestry peaks within 200 kb) across hybridising species.

Genome scans of Fisher p-values identified 29 genomic regions where evidence for adaptive admixture coincided with shared minor-parent ancestry islands (*p* < 0.00035 Bonferroni threshold; inferred with 200 kb *LAD* windows (*35*); Figure 5A), marking these genomic regions as candidates for parallel adaptive admixture. In contrast to iconic cases where adaptive introgression reflects isolated gene-capture events or older admixture (e.g., (*54–56*)), our results show that parallel adaptive introgression in southern GBR corals is sufficiently strong and pervasive enough that the same positively selected chromosomal regions have repeatedly crossed species boundaries, predictably restructuring genome-wide variation spanning multiple chromosomes (Figure 5A).

Further evidence that positive selection has influenced genome-wide introgression, comes from long-range linkage-disequilibrium (LD) patterns (*35*). Candidate regions for parallel adaptive admixture exhibited stronger associations across chromosomes in admixed populations (Kolmogorov-Smirnov Test, *p* < 1.10 x 10^-7^; Table S4) compared to random subsets of ancestry-informative sites or minor-parent ancestry peaks. These candidate regions also showed elevated proportions of sites in strong LD (e.g., r^2^ > 0.6; Figure S18) with clusters of SNPs associated with candidate loci on other chromosomes (e.g., Figure S19), indicating that genome-wide diversity in admixed populations is likely sustained by extensive LD among positively selected introgressed alleles. By contrast, if selection on introgressed variation were sporadic or inconsistent, LD would be restricted to independent sweeps rather than extend across chromosomes (*57, 58*). No candidate regions overlapped with *f_d_* outliers for *A. millepora* introgression, consistent with adaptive alleles derived from within the focal AH species trio rather than from *A. millepora*. Instead, *f_d_* peaks among AH species tended to occur within genomic regions lacking ancestry-informative sites, likely reflecting shared *A. millepora* introgressed ancestry.

Functional annotation and gene ontology assignments for candidate regions revealed signatures of positive selection across functionally diverse genes (Table S5), highlighting environmental stress as a possible driver of coral adaptation mediated by introgression. We detected positive selection signatures on a heat shock protein (Figure S20) as well as an enoyl-CoA hydratase-isomerase family gene on Chromosome 8 (Fisher *p* < 0.00035), where *A. hyacinthus* ancestry has risen to high frequency across numerous SNPs in admixed populations and coincides with parallel F_ST_ deviations between admixed and non-admixed populations (Figure 5B). Enoyl-CoA hydratase-isomerase enzymes play central roles in peroxisomal fatty-acid metabolism (*59*) and environmentally-induced antioxidant responses (*60*); they share functional similarities with stearoyl-CoA fatty acid desaturase (delta9-desaturase), a gene implicated in cold adaptation in *A. millepora* (*61*). Another positively selected region on Chromosome 4 lies within ∼300 kb from the *Sacsin* heat shock molecular chaperone (Figure S21), a previously identified thermal-tolerance candidate under balancing selection in *Acropora* corals (*62*). Although the *Sacsin* gene region lacked ancestry-informative sites in our dataset, it showed elevated nucleotide diversity and is in proximity to a candidate region for parallel adaptive admixture (Figure S21). Thus, adaptive introgression could provide an alternative explanation to balancing selection for creating elevated *Sacsin* diversity in *A. millepora* (*62*) and potentially within the *A. hyacinthus* species complex.

### Conclusions

Hybridisation is a pervasive evolutionary force (*2*), raising fundamental questions about how reproductive barriers persist despite gene flow and whether selection yields predictable genomic outcomes (*7*). Demonstrating adaptive functions for introgressed variation, however, remains challenging without establishing direct links to phenotypes and fitness (*7*). Here, we leveraged a multi-species admixture event among sympatric corals to test for adaptive introgression within the same ecological setting. We show that recent, geographically localised three-way admixture in the southern GBR has remodelled genomes in parallel across multiple AH species, driving rapid and repeatable adaptation over a few thousand generations. Using local ancestry inference for the first time in corals, we uncovered striking repeatability in minor-parent ancestry patterns across three hybridising species. These results indicate that predictable genomic outcomes across divergent genomes arose primarily from multi-genic positive selection favouring the same introgressed variants, whereby adaptive admixture has enabled the spread and reuse of beneficial alleles by drawing on shared standing genetic variation that transcends species boundaries. In GBR *Acropora*, selection may therefore favour incomplete barriers to gene flow, maintaining permeability in parts of the genome where access to adaptive genetic variation in related species is beneficial (*63, 64*). Our findings provide the first direct evidence that hybridisation in corals is not an evolutionary accident, but a deterministic mechanism shaping adaptation.

Our estimates of gene flow timing and its genomic extent indicate that admixed AH populations likely formed fewer than ∼2,000 generations ago following a breakdown in species barriers. Despite substantial admixture at southern reefs, AH species remain genetically distinct across the broader GBR, suggesting that reproductive isolation is maintained under most ecological contexts. Our results challenge the conventional view that geographic contact among semi-reproductively isolated species triggers admixture in sympatry. Instead, they suggest that other biological or environmental factors of geography determine when and where hybridisation occurs and influences evolutionary trajectories (*47*). Introgressive hybridisation may therefore be favoured only under specific ecological or demographic conditions (*8, 65*). We propose that range-edge dynamics in the southern GBR, such as reduced population sizes after colonisation, environmental change, or via limited access to conspecific mates, may have created episodic windows in which selection favoured some hybrids genotypes, temporarily enabling adaptive introgression across a wider interspecific gene pool, while preserving species boundaries elsewhere. Moreover, the coexistence of genetically differentiated admixed and non-admixed colonies at southern reefs suggests that admixed lineages have evolved to a stable state (*66*), implicating reproductive isolation with parental-type lineages that appears to be maintained by genome-wide, multi-genic selection (*67*) rather than selection localised to large-effect genetic incompatibilities. Together, our findings suggest that hybridisation may promote diversification in *Acropora* corals, generating differentiated mosaic lineages with potentially distinct phenotypes or enhanced adaptive potential that may facilitate responses to environmental change (*46*).

## Supplementary Methods and Materials

### Sample Collection

We collected adult colonies using SCUBA from 26 reefs across the Great Barrier Reef (Figure 1A). Sampling targeted colonies of the *A. hyacinthus* species complex (AH hereafter), recently taxonomically revised by (*32*), but morphologically similar under field conditions. All colonies were photographed at the colony and corallite scale, and GPS coordinates of colonies or sampling plots were recorded. All colonies were sampled under permits from the Great Barrier Reef Marine Park Authority (G19/43148.1, G21/45166.1, and G19/39364.1) and with Free Prior and Informed Consent of Traditional Owners of the Sea Country where the study sites were located. Samples were fixed in 100% ethanol after collection and stored at 20°C in the laboratory. A subset of the genotyped individuals was included in (*33*) (n=625; BioProject ID PRJNA982441) and new sampled colonies (n=303) were collected between 2021-2023.

### DNA Extraction, Genomic Library Preparation and Sequencing

Genomic DNA was extracted using the DNeasy QIAGEN Blood and Tissue kit and purified with SPRI magnetic beads following manufacturers specifications. Whole genome libraries were prepared using the Lotus DNA Library Prep Kit for NGS with 10 ng of input DNA and enzymatic fragmentation to achieve average insert sizes of 350 bp. Final amplification consisted of eight PCR cycles. Libraries were quantified using a Quant-iT dsDNA assay kit and up to 192 libraries were multiplexed in equimolar ratios for sequencing to achieve ∼10x coverage. Sequencing was performed by Azenta Life Sciences on the NovaSeq S4 300 cycle using 150 bp paired-end reads.

### Genomic Data Processing, Cluster Identification and Analyses of Species Affinity

We processed raw genomic data for a total of 928 coral colonies including *Acropora millepora* (n=11) and *Acropora kenti* (n=1) which were used as outgroups. FASTQC was used to examine read quality and adapter contamination and raw reads were trimmed in Trimmomatic v0.39 (*68*) using a 4 bp sliding window, a minimum phred-score quality of 20 and a minimum read length of 50 bp. Adapter sequences were removed using the Illuminaclip option in ‘palindrome mode’. Trimmed reads were mapped to the *Acropora hyacinthus* genome (GCA_020536085.1; specimen from Palau; (*69*)) using the Burrow-Wheeler Aligner (BWA, version 0.7.17; (*70*)) and the MEM algorithm with default settings. The resulting alignment SAM files were converted to indexed and sorted BAM files using SAMtools v1.13 (*71*). Read group information was added to individual BAM files and PCR duplicates were marked and removed using picard (http://broadinstitute.github.io/picard/). All samples with < 80% primary mapped reads aligning to the reference genome were removed to exclude potentially misidentified samples (n=26) from downstream analyses.

We initially visualised the genomic data using ANGSD v0.934 (*34*) to identify genetically distinct clusters. We identified polymorphic sites and estimated genotype likelihoods using the SAMtools method (*-gl* 1) to account for statistical uncertainty associated with sequencing errors and missing genotypes. The sites used for this analysis passed minimum filtering criteria of mapping quality 30, base quality 30, coverage ≥ 3 reads in at least 95% of individuals. We called major and minor alleles directly from the genotype likelihoods assuming biallelic sites and considered only polymorphic sites with a likelihood ratio test p-value < 0.000001. To evaluate population genetic structure, we extracted an individual covariance matrix with PCAngsd (*51*) applying a 0.05 minor allele frequency (MAF) threshold. We computed eigenvectors and eigenvalues in R and performed a Principal Components Analysis (PCA). We performed Bayesian hierarchical clustering admixture analyses in PCAngsd using the ‘admix’ option (NGSAdmix) (*72*) to estimate individual ancestry proportions assuming 2-6 ancestral populations (K genetic clusters) applying MAF = 0.05. PCA and clustering analyses supported the presence of four genetic clusters, which we hereafter refer to as Taxon 1 (*Acropora tersa*), Taxon 2 (*Acropora pectinata*), Taxon 3 (*Acropora sp.*) and Taxon 4 (*Acropora hyacinthus*) (refer to species assignment analyses below; Figure 1B, 1C)

Previous molecular studies (*24, 31*) suggest that the *A. hyacinthus* species complex is comprised of at least four genetically distinct lineages across the Pacific. To test the hypothesis that four genetically distinct AH species identified in (*33*) and in this study fall within nominal AH species based on recent taxonomic revisions (*32*), we sequenced eight voucher (taxonomist-verified) reference AH specimens collected from the Palm Islands on the GBR. These reference specimens are registered in the collection of the Queensland Museum Tropics, Townsville, and identified with reference to specimens identified by (*32*). We also downloaded and processed raw sequence reads from 20 AH samples from Ofu, American Samoa representing four genetic taxa HA, HC, HE and HD (n=5 per taxon; (*31*); BioProject: PRJNA657822; described in (*24*)). We then co-analysed these samples together with ten representative colonies from each distinct AH species from our GBR dataset, allowing up to 15% missing data in ANGSD. Based on the most recent taxonomy (*32*), we identified Taxon 1 as *Acropora tersa* (*32*), Taxon 2 as *Acropora pectinata* (*36*) and Taxon 4 as *Acropora hyacinthus* (*37*). Because Taxon 3 represents an undescribed species that was not included in (*32*), therefore, we refer to this species as *Acropora sp.* ‘Taxon 3’ hereafter.

### Clone Identification

Potential clones were identified following the approach of (*73*). This method identifies likely clones by applying a threshold on genetic similarities between colonies estimated as identity-by-state (IBS). Technical sequence replicates allowed us to determine the maximum pairwise genetic differences for multi-locus genotypes within each species. There were 10, 1 and 2 technical replicate pairs within *A. tersa*, Taxon 3 and *A. hyacinthus*, respectively. We applied the lower limit of relatedness (1-IBS) as a similarity threshold to identify putative clones. We used ANGSD to estimate IBS between colonies, where IBS is the proportion of randomly sampled reads covering a polymorphic site that are identical between compared individuals. For this analysis, we retained sites with MAF > 0.05 and a minimum of three aligned reads in ≥ 95% of samples within each taxon separately. Because there were no technical replicates in *A. pectinata*, we removed clones applying the similarity threshold from *A. tersa* (1-IBS = 0.179). We identified a total of 44 colonies as putative clones showing levels of pairwise genetic distances (1-IBS) below taxon-specific thresholds calculated as the maximum pairwise genetic difference among technical replicates. All clone pairs were sampled from the same reef. We retained a single representative for each clone pair and technical replicate with higher coverage for downstream analyses. In total, we removed 23 clones and 13 replicate individuals, keeping a single representative (higher coverage individual) for each pair. We additionally removed two samples exceeding > 20% missing loci, resulting in 834 individuals for downstream analysis.

### Population Genetic Structure and Global Ancestry Inference

To examine population genetic structure within and between species we first analysed taxon-specific datasets using the same filtering scheme as above, additionally removing sites exceeding a total read depth of 2 * taxon-specific mean (i.e., total depth per site across all colonies within each genetic group). Each species-specific dataset was then used as input for ngsparalog. ngsparalog jointly models sequencing depth, allele coverage ratios and heterozygosity excess relative to Hardy Weinberg proportions in order to identify and remove deviant sites due to mismapping errors, paralogy and copy number variations in low-coverage sequencing data (*74*). We subset the number of individuals to 80 per species to limit computational time and applied the same filtering as in ANGSD. From each taxon, we compiled a list of sites to exclude with a likelihood of being misaligned below *p* = 0.001 following (*75*). To reduce physical linkage between loci within each genetic cluster, we used ngsLD v1.2.0 (*76*) to quantify physical linkage and created separate datasets for linked and unlinked loci. We specified a maximum distance between SNPs as 50 kb (MAF = 0) and randomly sampled 80% of comparisons to identify linked sites with a squared correlation coefficient between genotypes r^2^ ≥ 0.5. For species-specific analyses of population structure, we estimated linkage decay and compiled lists of quality-filtered and linkage-pruned sites. For downstream analyses of population structure (all species combined) we merged quality-filtered sites and removed sites with MAF < 0.005 (equivalent to a minor allele count of four alleles across all 834 individuals) to exclude extremely rare, likely private alleles. We confirmed between and within-species population structure by calculating genotype likelihoods and performing PCA using PCAngsd applying a minimum MAF threshold of 0.05 (as above). Global ancestry proportions were estimated for each colony using NGSAdmix implemented in PCAngsd (‘admix’ option), using *A. millepora* as an outgroup and assuming 2-8 ancestral (K) groups.

### Genome-wide Analyses of Introgression

We first used TreeMix admixture graphs (*38*) to confirm the genetic relationships among AH species and *A. millepora* and to estimate migration between species by testing whether adding migration events improves the likelihood of the phylogeny relative to the tree with no migration. Following the genotype likelihood approach of (*77*), we first generated allele frequency files for each species using a common set of unlinked SNPS (as above) called across all individuals and forcing the major allele frequency to be that of the ancestor, where the *A. hyacinthus* genome was used as *-anc*. We then used the script *input_for_treemix.py* (https://github.com/joanocha/ngsIntrogression) to merge allele frequency files by chromosome, position and major allele to create input files for TreeMix, where minor allele counts were estimated by multiplying the allele (EM) frequency estimate for each site by the number of individuals (2Nind) and major allele counts were estimated by subtracting the minor alleles count from 2Nind. We then ran TreeMix assigning individuals to either genetically distinct non-admixed populations (central-northern GBR) or differentiated southern GBR populations and using *A. millepora* as an outgroup. We used options *-k500*, *global*, with standard error estimations and sample size correction allowing 0-9 migration events. We then used stepwise comparison Akaike information criterion (AIC) between sequential migration models to determine whether adding additional migration edges improved the likelihood of the population tree (≥AIC > 2) averaging across 10 replicate runs. We calculated AIC values as (−2(ln(likelihood)) + 2K), where K is the number of free parameters in the model and each migration edge adds three free parameters (edge weight, source, recipient).

To examine patterns of introgression among non-sister taxa, we computed genome-wide D-statistics (*39, 40*). This approach identifies excess sharing of derived alleles based on bi-allelic patterns ABBA and BABA. Given three groups – P1, P2, P3, and an outgroup, P4 – with the relationship (((P1, P2), P3), P4), ABBA patterns are sites where P2 and P3 share the derived allele B and P1 carries the ancestral allele A as defined by the outgroup. BABA sites are those where P1 and P3 share the derived allele and P2 carries the ancestral allele. Under incomplete lineage sorting, the number of ABBA and BABA sites should be equal, whereas a significant excess of ABBA sites results in a significantly positive D-statistics indicating gene flow between P2 and P3, and a negative D-statistic indicates gene flow between P1 and P3.

We computed genome-wide D-statistics in ANGSD (*-doabbababa2 1*) with a blocksize of 5 Mbp and only analysing quality filtered and unlinked sites where the outgroup is non-polymorphic (*-enhance 0*), followed by the Rscript *estAvgError.R* to obtain Z-scores and p-values for significance. Analyses assumed the relationship ((3,4),2),1), *A. kenti*) among focal AH species as inferred from the maximum likelihood TreeMix phylogeny (Figure S2). To better understand how introgression varies across the GBR species range, we performed the analysis separately using colonies sampled within reefs in the northern, central, and southern GBR regions (Figure 1A). We first tested for introgression between AH species by computing D-statistics using bam files aligned to the *A. hyacinthus* reference genome and assigning *A. kenti* as the outgroup (P4). We then tested for introgression between AH species and *A. millepora* using *A. kenti* as an outgroup. Significant differences in ABBA and BABA patterns were identified using a Z-test on the genome-wide mean D-statistic (where non-significant Z-score < abs(4)) and the bias-corrected D-statistic derived from jackknife resampling on “nBlocks” genomic blocks (i.e., V(JK-D)).

For comparisons showing significant genome-wide introgression between AH species and *A. millepora*, we examined putatively introgressed outlier genomic regions by quantifying the extent and direction of introgression in genomic windows. Because extreme values of D-statistics can arise in regions of low diversity (*41*), we adapted the ABBA-BABA test to calculate *f_d_*, the fraction of the genome showing excess admixture. *f_d_* compares the observed value of an introgression test to a scenario assuming maximum introgression and complete homogenisation of allele frequencies within a genomic window. Following the approach of (*77*), we repeated the ABBA-BABA test in 50 kb windows and ran *- doabbababa2* using all quality filtered sites to obtain numerators for combinations P1,P2,P3,P4 and P1,P2,P2,P4 and P1,P3,P3,P4. We then extracted results for these taxon combinations and merged outputs using an adapted version of the python script *Fraction_admix.py* (https://github.com/joanocha/ngsIntrogression) to obtain the fraction of the introgressed genome *fd* = numerator_(P1,P2,P3,P4)_/numerator_(P1,PD,PD,P4)_, where PD = P2 or P3, depending on which taxon has the higher derived allele frequency (to maximise the denominator). To identify *f_d_* outliers, genomic windows above the 99^th^ percentile of the empirical *f_d_* distribution (as in (*77*)) were considered as introgression outliers from P3 (donor) into P2 (recipient) (=positive D-statistic).

To determine the region of the *A. hyacinthus* alignment corresponding to the HES1 locus as in (*31*), we used Ragtag (*78*) to align the *A. hyacinthus* (query) and *A. millepora* (target) genomes. We extracted alignments with a minimum mapping quality >20 and alignment length >5 k base pairs to determine the coordinates in the query genome overlapping the target HES1 region in *A. millepora* (i.e., Chromosome 7: 20.35-20.57 Mb).

### Genomic Analysis for Demographic Inference

To investigate the timing of divergence and gene flow among AH species and *A. millepora,* we reconstructed demographic histories using the Diffusion Approximation of Demographic Inference (*dadi*) v.2.3.3 (*50*). To generate input files for *dadi* analyses, we used taxon-specific datasets, quality-filtered sites for maximum depth, and used ngsparalog to exclude potentially paralogous sites (as above) but excluding MAF filters. We selected 20 individuals with global major ancestry q-values >0.98 and ensured similar coverage among samples. Individuals were selected from sympatric populations from the central and northern GBR, excluding colonies from southern (admixed) and far northern Torres Strait populations. Using realSFS, we generated folded 1D Site Frequency Spectra (SFS) for each GBR AH species (n=20) and *A. millepora* (n=11), retaining only sites on 14 assembled chromosomes and with a minimum mapping quality 30, base quality 30 (i.e.,*-minMapQ 30 -minQ 30 -remove_bads 1 - uniqueOnly 1*), and coverage ≥ 3 reads in all individuals (i.e., no missing data). 2D-folded SFS were generated for each pairwise combination (four AH species and *A. millepora*) using 100 bootstrap replicates. Each SFS was projected to 80% of haplotypes in each population for smoothing and handling of missing bins.

### Modelling Demographic History Using dadi

Unmodeled population size changes may bias model choice towards migration scenarios and affect parameter estimation (*79*). We therefore first used *dadi* to test for population size changes within each species in the 1D-SFS by comparing four models: (i) constant population size (i.e., standard neutral model), (ii) instantaneous population size change, (iii) a two-epoch model with two instantaneous population size changes (i.e., bottleneck) and (iv) a two-epoch model with exponential population size change. For each taxon, models including population growth were favoured (Table S6), and thus two population size changes across two time periods were incorporated into downstream 2D models.

To assess the demographic history of each species pair, we used the Genetic Algorithm of Demographic Model Analysis (GADMA) software v2.0.3 (*80, 81*) for global optimisation comparing among replicates while using the *dadi* engine for local optimisation within each demographic model. The global optimiser compares models with and without certain parameters in the optimal model; we set population size changes as sudden and migration as asymmetric. The initial structure of the model was 1,1 with final structure 1,2, which represents the number of epochs before and after divergence. Selecting for two epochs after divergence allows for the assessment of typical two population divergence scenarios (isolation with migration, ancient migration, and secondary contact). As SNPs were linked, we used the CLAIC across bootstraps (*82*) and log-likelihood model comparison. Global optimisations were run sixteen times with eight replicates and four processes within each replicate for each population pair, essentially allowing 128 optimisation comparisons per pair. During this optimisation, the more complex models were preferred. We then customised the GADMA optimised models by adding variable gene flow rates across the genome, the rationale being that (i) GADMA only allows global optimisation of specific models that do not include heterogeneous gene flow (i.e., variable rates of gene flow across the genome) and (ii) we know that heterogeneous gene flow is an important feature of genetic mixing among semi-reproductively isolated taxa that may exhibit genomic barrier loci (*83*). We created two different heterogeneous gene flow models: one model (named ‘*2het*’) with two variable migration rates that contribute across different proportions of the genome, *P* and *1 - P*, and another model (named ‘*1het*’) that has one migration rate (i.e., single heterogeneous gene flow) at proportion *1 - P* and no migration at P. Both models included different genomic proportions across two epochs (*P_E1_* and *P_E2_*) and were run with the same optimisation scheme with 128 comparisons per pair per model. Because the migration rates for one of the proportions in the *2het* models always optimised to the lower bound of the migration parameter, these models essentially optimised to be concordant with *1het* models. We therefore present results from the single heterogeneous gene flow models, ‘*1het*’.

We performed confidence interval calculations on models containing the highest log-likelihoods and the top five models were compared for parameter consistency. Using bootstraps provided by realSFS, confidence intervals were calculated using the Godambe algorithm (GIM.uncert; (*82*)) and different EPS settings were used for each parameter at roughly 1/100th of the value of the optimised parameter. If the lower confidence intervals hit zero for any parameter, then nested models (models excluding these zero-hitting parameters) were optimised (using the same optimisation scheme) and log-likelihood ratios tests were applied to assess whether to include the tested parameters in the final model.

### Locus-Specific Local Ancestry Inference in Admixed Populations in the Southern GBR

To investigate the timing of admixture among hybridising AH species in the southern GBR and assess the repeatability of hybridisation outcomes (refer to next section), we used Ancestry-HMM (*48*) to jointly infer the timing of multiple admixture events and to map locus-specific ancestry patterns across the genome. We estimated local ancestry for each sample using read counts at each locus from southern-admixed populations and central-northern parental-type (non-admixed) populations. Taxon-specific reference panels were constructed using parental-type individuals from the far north, northern and central GBR regions, but excluding individuals that deviate towards admixed or interspecific groups based on PCA or those with intermediate global ancestries to avoid biases resulting from uncertain assignment to source populations. We first generated a Beagle file for each reference parental-type population for *A. pectinata* (n=60), Taxon 3 (n=39) and *A. hyacinthus* (n=58) using quality-filtered sites without MAF filters (as above). Following recommendations from (*48*) and (*49*), sites in linkage disequilibrium (LD) (r^2^ > 0.4) were removed from each reference population (but not in admixed populations to retain admixture LD). We discarded a single random SNP from each linked pair of sites within 50 kb of each other from the master sites file using ngsLD. While this filtering step led to the loss of ∼50 k ancestry-informative sites (F_ST_ ≥ 0.1), pruning of ancestral LD is necessary for unbiased admixture time estimates (*48, 49*).

Next, we generated a BCF from genotype likelihoods with *-doBcf* including all three species-specific reference populations and a single southern-admixed population resulting in three separate datasets. The BCF file was converted to a VCF using bcftools and genotype likelihoods were converted to genotypes using beagle (v4.1; (*84*)). The resulting VCF files were used to extract reference allele counts for ancestry informative sites with a minimum of 10 samples with data in each reference population and a minimum allele frequency difference of 0.1 among reference populations using the script *vcf2ahmm.py* (https://github.com/jesvedberg/Ancestry_HMM-S). Next, a sites file was created from the ancestry-informative reference sites passing *vcf2ahmm.py* filters and used to estimate site-specific read counts for each admixed individual using ANGSD (*-dumpCounts 3*). We then converted A/C/G/T counts into major/minor allele counts for each reference population using a modified scripts from (*85*) (https://github.com/ecalfee/bees/tree/master/local_ancestry) and reordered each input file with the parental-type (ancestral) population first followed by admixed populations.

We ran Ancestry_HMM using our NGSAdmix results as a prior for population ancestry proportions (0.8, 0.1, 0.1) and did not set the population size option. We modelled simple three-way admixture among southern GBR admixed populations (*A. pectinata*, Taxon 3 and *A. hyacinthus*). We ran three separate analyses, starting with each respective ancestral (major-parent) population then specifying two distinct admixture events with introgressing taxa or migration ‘pulses’ (*49*), where each subsequent pulse replaced 10% of total ancestry in the ancestral population as an initial estimate that is optimised when fitting the admixture model. Two pulse model were as follows: *Model 1*: *A. pectinata* (ancestral), Taxon 3 (pulse 1), *A. hyacinthus* (pulse 2); *Model 2*: Taxon 3 (ancestral), *A. pectinata* (pulse 1), *A. hyacinthus* (pulse 2); *Model 3*: *A. hyacinthus* (ancestral), *A. pectinata* (pulse 1), Taxon 3 (pulse 2). We specified an initial admixture time of 1,000,000 generations to account for distant admixture and used priors for admixture pulses ranging from 10 k and 100 k generations ago; because these priors had no effect on migration time estimates and model likelihoods, we proceeded with a final run using priors for the first admixture pulse at 10 k generations ago and a second pulse 1000 generations ago running 100 bootstrap replicates and a block size of 5000 SNPs. Using the output, we calculated ancestry proportions at each SNP by marginalising over the posterior probability for homozygous and heterozygous ancestries inferred by Ancestry_HMM (i.e. *A* = *p*(*AA*) + 1/2(*p*(*BA*) + *p*(*CA*)) (as in (*85*)). Based on the rates of linkage decay in parental populations at ∼5-10 kb (Figure S22), we calculated ancestry components in non-overlapping 20 kb, 50 kb, 100 kb and 200 kb windows using the R package ‘windowscanr’ for downstream analyses.

### Genome-wide Local Ancestry Correlations

To test whether hybridisation has resulted in repeatable genomic outcomes among the major-parent backgrounds and specifically whether adaptive introgression has predictably reconfigured major-parent genomes, we compared patterns of local ancestry between different southern-admixed AH species. We were specifically interested in identifying regions where selection may maintain high proportions of minor-parent ancestry across two or more populations. We tested for correlations between the same minor-parent ancestry between southern-admixed populations and major-parent ancestry across the genome (excluding unplaced scaffolds) and for each chromosome individually. We compared per-locus ancestry and mean ancestry calculated in non-overlapping 20 kb, 50 kb, 100 kb and 200 kb windows to account for recombination rate variation and used a Spearman’s correlation test implemented in R. We then explored whether the number of shared minor-parent ancestry peaks – defined as genomic regions where the same minor-parent ancestry is maintained at high proportions > 50% among two admixed populations – exceeded what would be expected by chance by running 10,000 permutations of the data in R. To account for linkage among sites and preserve local ancestry structure across the genome, we repeated permutation tests on mean ancestry calculated in genomic windows of 20 kb, 50 kb, 100 kb and 200 kb.

### Annotation and Enrichment of Gene Ontology Terms

Next, we examined functional enrichment among genes involved in introgression. We downloaded predicted protein sequences for the *A. hyacinthus* genome assembly (https://github.com/eloralopez/CoralGermline) and performed functional annotation on the predicted gene models using the EggNog-mapper webserver with default settings (http://eggnog-mapper.embl.de). To identify enriched Gene Ontology (GO) terms among private (non-shared) and shared minor-parent ancestry peaks, we used the topGO R package (*86*) with the *weight01* algorithm, considering only GO terms assigned to at least ten genes (node size = 10) and the Fisher’s exact test to assess significantly enriched Biological Process categories (*p* ≤ 0.05).

### Testing for Positive Selection on Admixed Variation using FADM, LAD and Fisher’s Method

To test the hypothesis that positive selection underlies repeated introgression of the same minor-parent alleles across multiple major-parent backgrounds, we examined patterns of diversity expected under admixture and selection to identify genomic regions involved in parallel adaptive introgression across southern-admixed AH species’ boundaries.

We calculated two neutrality statistics, *F_adm_* and *LAD,* that have substantial power to detect positive selection in admixed genomes (*52*). By identifying outliers in local ancestry and allele frequency variation, these statistics are well-suited to detect loci under selection during ‘adaptive admixture’–where beneficial alleles are introgressed from parental species and remain advantageous and selectively favoured in hybrid populations, rather than selection acting solely on standing variation in hybrid populations post-admixture (*52*). Because post-admixture selection on standing variation in admixed populations (that is not beneficial in parental source populations) is likely to act on multiple haplotypes, these statistics are not confounded by signatures of post-admixture selection where the beneficial allele is neutral in the source population or no longer adaptive in the admixed population (*52*). Additionally, these statistics also allow us to focus on adaptive admixture events leading to moderate admixture changes in the last few 1000 generations.

First, we computed *F_adm_,* which measures the deviation in the observed allele frequency from the expected allele frequency under admixture and neutrality for each allele at a polymorphic site, normalised by the expected heterozygosity in the admixed population

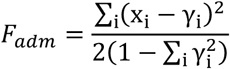

where *γ_i_* is the sum of the average allele frequency in each source parental population weighted by the genome-wide admixture proportion. *F_adm_* detects selection in the admixed population and is independent of local ancestry inference, because it relies only on allele frequencies and is robust to error in the estimate of admixture proportions and possible associated errors. To compute *F_adm_,* we calculated normalised average genome-wide admixture proportions for three parental ancestry components, for each population. Allele frequencies were calculated in ANGSD assigning the major allele as the reference *(-doMajorMinor 4 -doMaf 1*), where each file was generated using a common set of SNPs for ancestry inference (as above). We excluded sites where *x_i_* in the admixed population is higher or lower than the frequency in the source populations as recommended by (*52*).

Next, we computed the local ancestry inference-based statistic *LAD*, which measures the per-window local ancestry deviation from the genome-wide average ancestry estimated using local ancestry inference:

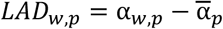

Where *α_w,p_* is the admixture proportion from a given source population *p* for a given window *w* and *α̅*_*p*_ is the genome-wide admixture proportion estimated from local ancestry inference averaged across loci. The rationale is that when beneficial alleles are transferred from a source population to the admixed population, local ancestral proportions surrounding these loci are expected to increase relative to neutral loci and the genomic mean. As above, we calculated *LAD* in non-overlapping 50 kb, 100 kb and 200 kb windows.

Next, we combined *F_adm_* and *LAD* statistics by ranking SNPs using Fisher’s Method defined as:

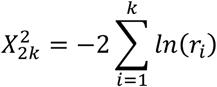

Where *k* = 2 is the number of statistics and *r_i_* is the rank of a given SNP for statistic *i* divided by the total number of analysed SNPs to obtain the empirical p-value. Thus, each SNP was assigned three *LAD* values (one for each parental ancestry), and all SNPs within the same window were assigned *LAD* values. We applied a conservative Bonferroni correction on Fisher’s p-value threshold of 0.05, divided by the number *LAD* windows as the number of tests.

### Genome Scans of Differentiation and Diversity

To identify genomic regions involved in parallel genetic differentiation between southern-admixed AH populations and non-admixed parental-type central-northern populations, we performed PCA-based selection scans using the *-selection* flag in PCAngsd (*51*) and excluding *A. tersa*. This method uses posterior genotype expectations to identify SNPs exceeding neutral expectations along a focal axis of genetic variation. This test statistic follows a χ^2^ distribution with 1 degree of freedom and we calculated per-SNP probabilities (MAF < 0.05) using the *pchisq* function (lower.tail = FALSE) for the third principal component in R. Loci falling above the 99.9^th^ percentile (top 0.01%) of the empirical p-value distribution were considered as outliers of genomic differentiation considering the third major axis of differentiation (PC3).

To examine genetic diversity and differentiation in candidate genomic regions for parallel adaptive admixture (identified with tests of positive selection as described above), we generated 1D-SFS and unfolded 2D-SFS using realSFS for each population pair and calculated pairwise F_ST_ in non-overlapping windows of 50 kb (10 kb sliding window) in ANGSD. We used folded 1D-SFS and *sat2theta* with the *thetastat do_stat* option to calculate Watterson’s theta and estimated allele frequencies assigning the major allele as the reference *(-doMajorMinor 4 -doMaf 1*).

### Linkage Network Analysis of Candidate Genomic Regions for Parallel Adaptive Admixture

Constructing linkage disequilibrium (LD) networks can provide valuable insights into the architecture of local adaptation (*57*). Introgressed loci under selection are expected to display stronger associations across chromosomes relative to other genomic regions if selected genomic backgrounds are concurrently favoured following admixture. By contrast, if selection on introgressed variation were sporadic or inconsistent, linkage disequilibrium would be restricted to independent sweeps rather than extend across chromosomes (*57, 58*). To investigate these patterns among candidate genomic regions involved in parallel adaptive admixture, we calculated LD among loci within admixed populations using ngsLD v1.2.0 for three datasets for each species: (Dataset 1) comparisons among all minor-parent ancestry peaks (sites exceeding 50% minor-parent ancestry proportions) within each major-parent background (private and shared among species); (Dataset 2) comparisons among sites within 29 candidate genomic regions for parallel adaptive admixture (normalised to 200 kb base pairs wide); and (Dataset 3) comparisons among a randomly selected subset of ancestry informative sites equivalent to the number of minor-parent ancestry peaks (Dataset 1: *A. pectinata*, n = 13,760; Taxon 3, n = 8,913; *A. hyacinthus*, n = 7,287). For each dataset, we estimated associations between pairs of loci across all chromosomes, computed probability density function curves in R and compared LD distributions using two-sample Kolmogorov-Smirnov tests. We also visualised the location of sites in Dataset 1 with strong associations r^2^ ≥ 0.6 with at least one site within candidate genomic regions in histograms of 200 kb bins.

## Results

### Population Genetic Structure, Species Affinities and Global Ancestry Inference

We analysed 823 AH and 11 *A. millepora* colonies that were sequenced with an average of 26,634,808 reads per sample (range = 16,829,743 to 63,841,053) following quality filtering. Samples had an average of 93.12% primary reads mapped to the reference genome and mean genome-wide coverage among individuals of 11.5x and 7.05x across 14 assembled chromosomes. PCA on 92,669 genome-wide unlinked SNPs (Figure 1C) showed four AH genetic clusters comprised of previously analysed (*33*) and newly sampled colonies. As expected, all four genetic clusters occurred in sympatry across the GBR and appeared to represent semi-reproductively isolated AH species, which we refer to be *A. tersa* (Taxon 1), *A. pectinata* (Taxon 2), Taxon 3 (*Acropora sp.*) and *A. hyacinthus* (Taxon 4) (see species assignment below; Figure 1B, 1C). The primary principal components PC1 and PC2 explained 10% and 3.44% of total genomic variation respectively (Figure 1C).

We contextualised the identity of the four AH species in the GBR by co-analysing ten representative colonies per taxon alongside eight specimens from the Queensland Museum collection identified with reference to type materials and specimens examined by (*32*), as well as 20 AH samples from Ofu, American Samoa (HA, HC, HE and HD; (*31*)). Based on the most recent AH taxonomy (*32*), we confirmed Taxon 1 as *Acropora tersa* (referred to as *A. hyacinthus* “neat” in (*33*) prior to the formal description of *A. tersa* by (*32*)), Taxon 2 as *Acropora pectinata* and Taxon 4 as *Acropora hyacinthus*. Taxon 3 represents a currently undescribed species that was not included in (*32*). Analyses with representative specimens belonging to AH species from Ofu, American Samoa (HA, HC, HE and HD; (*31*)) showed close affinities on PC1 and PC2 between Taxon 1 and HC, Taxon 2 and HA, Taxon 4 and HD. Taxon HE did not show clear affinities to AH species in the GBR (Figure S23).

Genomic divisions among AH colonies in the GBR were supported by Bayesian clustering analyses in NGSAdmix (Figure 1D). Six colonies showed high mixed admixture proportions (40-70%) consistent with intermediate placement between major genetic clusters in PCA, suggesting they are putative recent hybrids (Figure 1D). These colonies were sampled primarily from central and northern GBR regions, with colonies from Pelorus Island (n = 2; central), Moore Reef (n = 2, north), Chicken Reef (n = 1, central) and Little Broadhurst Reef (n = 1, central). One exception was a colony with mixed *A. hyacinthus* and *A. tersa* ancestry from Lady Musgrave Reef (south). Minor global ancestry assignment (< 5%) to *A. millepora* was also evident among all AH colonies (Figure 1D).

We found complex patterns of population structuring by latitude within three AH species: *A. pectinata*, Taxon 3 and *A. hyacinthus* (Figure 1D), whereas *A. tersa* showed no clear evidence of latitudinal segregation. PCA of *A. pectinata*, Taxon 3 and *A. hyacinthus* showed clear separation on PC1 between some southern colonies (Capricorn Bunker and Swains reef groups; Figure 1E) and central and northern populations, although some southern colonies clustered with northern and central populations. NGSAdmix analysis showed that these distinct southern populations were comprised entirely of admixed individuals deriving an average of 3.5%-14.8% of their genomes from two other AH species. (Figure 1E), while other southern colonies were non-admixed and genetically similar to parental-type taxa found GBR-wide (Figure 1E). PCA on a combined dataset of *A. pectinata*, Taxon 3 and *A. hyacinthus*, showed closer affinities among southern-admixed populations, consistent with three-way hybridisation (Figure 2A; Figure S1). These distinct southern-admixed populations also diverged in parallel on PC3 from non-admixed parental-type populations found GBR-wide (Figure S7-A). Using PCA-based selection scans in PCAngsd, we identified genome-wide outliers exceeding the top 0.01% of the empirical -log10(p-value) distribution that significantly contributed to differentiation between southern-admixed and non-admixed parental-type populations (Figure S7-B).

### Genome-wide Analyses of Introgression

The maximum likelihood population tree generated with TreeMix explained 96.1% of variation in the allele frequency covariance matrix (Figure S2). The addition of three migration edges returned the highest likelihood tree (based on AIC comparisons) and explained > 98% of the variation.

We calculated genome-wide D-statistics using *A. kenti* as an outgroup to test for gene flow among AH species and with *A. millepora* in the northern, central and southern GBR regions (Figure 1A). We found evidence for variable introgression among AH species in different parts of the GBR range (Table S1). In the central and northern GBR, only comparisons between *A. pectinata* and Taxon 3 were significant (absolute *D*-statistic = 0.10, Z Score > 30; Figure 2B). Whereas in the southern GBR, all comparisons between admixed *A. pectinata*, Taxon 3 and *A. hyacinthus* were significant (absolute *D*-statistic range 0.01-0.12; Figure 2B), supporting localised three-way introgression in this region. We also found weaker evidence for introgression between *A. tersa* and non-admixed *A. hyacinthus* at southern reefs (i.e., Taxon 4-1; Table S1).

Next, we calculated D-statistics among AH species and *A. millepora* to test whether population structure and genetic similarities observed among AH species from southern reefs could be driven by shared past *A. millepora* introgression. Using *A. kenti* as an outgroup, we found a significant excess of allele sharing between *A. millepora* and *A. pectinata*, Taxon 3 and *A. hyacinthus* (abs(*D*) range 0.01-0.04; Figure 2B) in the northern, central and southern GBR regions (Table S2), but not with *A. tersa*, indicating past gene flow between *A. millepora* and AH species, which was consistent with minor global ancestry assignment (< 5%) of AH colonies to *A. millepora* (Figure 1D). To locate introgressed genomic regions putatively involved in adaptive introgression, we calculated *f_d_* the fraction of the genome showing excess admixture, in sliding 50 kb windows. For all comparisons within GBR regions, genome-wide *f_d_* values were centred around zero (Figure S3) and few genomic windows showed extreme values in the top 1% of the empirical *f_d_* distribution that might implicate adaptive introgression (Figure S3; Figure S5). The top 1% of *f_d_* outliers revealed several introgression hotspots with excess ancestry on all chromosomes, including large chromosomal segments up to 400 kb in length (e.g., Figure S4), including a Chromosome 4 region corresponding to the HES1 locus speculated to be involved in *A. millepora* introgression and bleaching tolerance in AH species in American Samoa (*31*). *f_d_* outlier windows were generally concordant among all three AH species and across GBR regions (Spearman’s *ρ* ≥ 0.50, *p* < 3.24 x 10^-110^; Figure S5), supporting ancient introgression with *A. millepora* that does not contribute to present day genetic segregation in southern-admixed AH populations.

### Genomic Local Ancestry Inference and Admixture Timing

We estimated locus-specific local ancestry patterns across genomes of southern-admixed AH species using final datasets of 131,725, 132,004 and 131,662 ancestry informative sites for *A. pectinata*, Taxon 3 and *A. hyacinthus*, respectively. The average minor-parent ancestry estimates inferred from Ancestry_HMM were slightly higher values for individual colonies (range: 21.4%-26.4%) than global estimates from NGSAdmix (range: 3.5%-14.8%), suggesting that introgression rates are relatively high. Several genome-wide patterns of local ancestry emerged (Figure S24). Major-parent ancestry distributions were centred around 50%, with many sites showing balanced ancestry proportions for both major and minor-parent ancestries, likely reflecting heterosis, or genomic regions where introgressed alleles segregate neutrally or where there is little power to differentiate among parental ancestries. Minor-parent distributions were left skewed and centred around 20% with few regions showing minor-parent ancestries exceeding 50% (Figure S24).

Admixture times inferred by Ancestry_HMM were not sensitive to initial admixture time priors. Analyses consistently returned similar admixture times between 1,000-2,000 generations for all populations (Figure S6). The most recent admixture pulse was for *A. pectinata*, with introduced Taxon 3 ancestry (first pulse) 1,044 generations ago [95% CI: 1,016-1,072], while admixture with *A. hyacinthus* (second pulse) occurred 1,130 generations ago [95% CI: 1,092-1,168]. Slightly older estimates of 1,591 [95% CI: 1,544-1,639] and 1,924 [95% CI: 1,812-2,035] generations ago were inferred for Taxon 3 and *A. hyacinthus* respectively; however, the models could not distinguish between the timing of the first and second admixture pulses (Figure S7). We reasoned that Ancestry_HMM models could not discern between admixture events occurring within similar time periods, if a relatively small proportion of the total ancestry in the admixed population was introduced by the second migration pulse (*49*) or if periodic or ongoing admixture violated the key assumption of discrete hybridisation ‘pulses’ (*87*). Additionally, limited numbers of ancestry informative sites due to modestly differentiated parental populations (e.g., mean F_ST_ < 0.1), relatively recent admixture events (< 5,000 generations ago) and a lack of a recombination map may have further limited our admixture timing inferences (*49*). Overall, estimates suggest that three-way admixture among AH species from the southern GBR happened relatively recently in the past (i.e., during the Holocene) following the Last Glacial Maximum (LGM) starting between 3,000-5,000 years ago depending on the generation time of 3-5 years. This figure is congruent with geological evidence from reef cores that coral growth was likely minimal in the southern GBR during much of the last ice age, with tabular *Acropora* appearing on the southern GBR within the last 7,500 years (*43, 44*).

### Reconstructing Demographic Histories of Divergence and Gene Flow Using dadi

We performed demographic modelling to reconstruct the evolutionary histories of gene flow among three focal AH species and contrasted these patterns with those observed with *A. tersa* and *A. millepora*. This allowed us to determine whether the observed admixture among AH species reflects ongoing or episodic gene flow, or whether migration has only occurred post-LGM affecting species simultaneously (Figure 3). We drew on non-admixed parental-type populations from the central-northern GBR to resolve deeper historical patterns of interspecific gene flow among three focal AH species and contrast patterns with those observed with *A. tersa* and *A. millepora*.

Initial GADMA optimised models supported ongoing gene flow between *A. tersa* comparisons (with *A. pectinata,* Taxon 3 and *A. hyacinthus*) and secondary contact for all other comparisons among three focal AH species (*A. pectinata*, Taxon 3, and *A. hyacinthus*) and *A. millepora*. However, heterogeneous gene flow models (Figure S25; Figure S26; Figure S27; Figure S28) presented slightly different results compared to initial homogeneous models. Divergence times among paired comparisons for *A. pectinata*, Taxon 3, and *A. hyacinthus* were consistently around 300 k generations ago (range: 286-325 k generations; Figure 3). In the first time-period, these three species experienced moderate gene flow (*Nm*: 91-130) but maintained strong genome-wide barriers affecting the majority of the genome (*P_E1_* = 0.54-0.66; Table S3). In the second time-period, however, we observed patterns consistent with a more recent breakdown of species barriers, whereby gene flow at generally higher migration rates affected up to 84% of the genome (*P_E2_* = 0.16-0.40) ∼2,000-4,000 generations ago, consistent with admixture timing inferred from local ancestry inference in southern-admixed populations (Figure 3; Table S3).

For paired comparisons between *A. tersa* and Taxon 3 or *A. hyacinthus*, we found divergence times ∼350 k generations ago with minimal ongoing gene flow (*Nm* < 1), except for higher migration from *A. tersa* to *A. hyacinthus* (*Nm* = 8.47) across > 70% of the genome (*P_E1_* = 0.16-0.36) in the first time period (*T_E1_*) (Figure 3). By contrast, more of the genome experienced no gene flow in the second time-period (*P_E2_* = 0.29-0.46), indicating increased genomic barriers to migration (Figure 3). For comparisons between *A. tersa* and *A. pectinata*, divergence times were consistently more recent, approximately ∼80 k generations ago, relative to divergence times with Taxon 3 and *A. hyacinthus* (Table S3). For this species pair, the more complex gene flow model (*het1*) had multiple parameters hitting the lower boundary (*N1*, *Nm1_12*, *Nm1_21*, *P_E1_* and *T_E2_*); log-likelihood ratio tests between the complex model and two nested models excluding these parameters (*hetim:* continuous gene flow; *hetsc:* secondary contact) found no significant differences, thus favouring the nested models (Table S7). Among nested models, the secondary contact model (*hetsc*) was favoured with higher log-likelihood and confidence intervals were reasonable (none hitting lower parameter bounds) compared to the continuous gene flow model (*hetim*) (Table S7). In the secondary contact model, the onset of contact was ∼20 k generations ago with moderate gene flow (*Nm* = 6.4-19.4) across ∼70% of the genome (*P_E2_* = 0.29).

Divergence times between AH species and *A. millepora* and were consistently older, ∼400 k generations ago (range 363-444 k generations). There was no or minimal gene flow in the first time period across a large proportion of the genome (*P_E1_* = 0.15-0.26) and low to moderate (*Nm* = 6.7-61.7) ongoing gene flow in the second period. A larger proportion of the genome (*P_E2_* = 0.57-0.6) experienced no genetic exchange at the onset of the second pulse of genetic exchange ∼45,000 generations ago with *A. tersa* and *A. pectinata* and ∼25,000 generations ago with Taxon 3 and *A. hyacinthus*.

### Parallel genome-wide outcomes of hybridisation

To investigate the repeatability of genome-wide local ancestry patterns in multiple hybridisation events, and to test whether selection has led to parallel adaptive introgression, we inferred genome-wide local ancestry states and quantified correlations in local ancestry between admixed populations. We identified significant positive correlations in minor-parent ancestries based on ancestry informative sites (Spearman’s *ρ* range 0.51-0.66; *p* < 1 x 10^-16^; (Figure S9), such that minor-parent ancestry in one major-parent background strongly predicted the same minor-parent ancestry at the same genomic location in the other major-parent background. Significant correlations in mean minor-parent ancestry were also consistent at various window sizes from 20-200 kb (Spearman’s *ρ* range 0.46-0.68; *p* < 1 x 10^-16^ ; Figure 4; Figure S10) and across chromosomes (Figure S11) suggesting that repeatability in broad and fine-scale ancestry patterns likely persists under varying recombination levels and across chromosomes. Hundreds of ancestry-informative sites (1.3%; Figure S12) contributed to shared minor-parent ancestry peaks in two or more major-parent genomes. Out of 122,977 common ancestry informative sites, an average of 5.6% sites were private (non-shared) minor-parent ancestry peaks (Figure S12). We identified 1,229 (1%), 959 (0.8%) and 2,496 (2%) sites with shared minor-parent ancestry peaks in two taxa for *A. pectinata*, Taxon 3, and *A. hyacinthus* ancestries, respectively. Relative to expectations by chance based on 10,000 permutations, admixed genomes were >5 times more enriched for overlapping introgressed loci considering both ancestry sites (*p* < 4.9 x 10^-4^; Figure S13) and mean ancestry calculated across various window sizes (20 kb to 200 kb) exceeded sharing excepted by chance (*p* < 1 x 10^-16^ ; p < 0-0.0004). Of these shared minor-parent peaks, >95% of these shared minor-parent peaks corresponded to genomic regions where the same allele reached high proportions (>50%) in the major-parent background (Figure S12). By contrast, major-parent ancestries were weakly correlated among species (Spearman’s *ρ* range -0.17-0.3, *p* < 7.9 x 10^-56^), considering both ancestry informative sites and average ancestry proportions within 20-200 kb windows (Figure 4; Figure S14; Figure S15; Figure S16).

Most hybridisation events involve the interplay between negative and positive selection acting during different phases of evolution following admixture (*8*). Thus, several models of selection could explain the concordant patterns we observe, including purging of minor-parent ancestries due to linked selection surrounding shared barrier loci or deleterious introgressed alleles, as well as ecological selection repeatedly favouring the same minor-parent alleles in different species in the same ecological setting (*8, 88*). Correlations in minor-parent ancestry are also expected to emerge because neutral introgressed alleles are more likely to persist in genomic regions where high recombination rates quickly decouple them from deleterious variants (*9, 14, 16, 89, 90*). Although we were unable to investigate the relationship between local ancestry patterns and recombination rates, we do not expect different minor-parent ancestries to reach high frequencies in different parental backgrounds without a shared history of positive selection on the same alleles in hybrid genomes (*12*). Nevertheless, strong genetic drift following admixture may also substantially altering local ancestries in early-generation hybrids and result in parallel outcomes by chance in different major-parent genomic backgrounds (*85*). Masking of weakly deleterious alleles in small, receiving populations through heterosis could also favour the introgression of the same minor-parent alleles from the species with a higher effective population size (and fewer deleterious mutations), especially in low recombination regions (*10–12*). However, our best demographic models indicated similar effective population size among AH species so this scenario could be less plausible (approximate *Ne* range ∼500,000-1,000,000 individuals; Table S3). Under both scenarios, however, we would not expect the same minor-parent ancestry to repeatedly rise to high frequencies in the same genomic regions whereby parallelism is consistent across all three minor-parent ancestries (Figure 4), with different introgressed ancestries repeatedly favoured in different genomic regions, producing a ‘mix-and-match’ genomic mosaic that is consistent with adaptive introgression of shared targets of selection in divergent AH species following hybridisation. Knowledge of local recombination rates and mapping the locations of genomic incompatibilities is lacking for coral species but would provide important guides for assessing the impact of these processes on genetic variation.

### Functional Enrichment of Gene Ontology Terms at Private and Shared Minor-parent Ancestry Peaks

To examine functional enrichment among genes involved in introgression, we obtained GO terms for 7,470 out of 27,110 predicted protein sequences in the *A. hyacinthus* assembly. Candidate genes overlapping with private (non-shared) minor-parent ancestry peaks were enriched for GO terms involved in *reproduction* as the top Level-2 Biological Process (*p* ≤ 0.0001; Figure S17-A). In contrast, genes within shared minor-parent ancestry peaks were significantly enriched for GO terms associated with *response-to-stress,* as the top functional category (*p* ≤ 0.0001; Figure S17-B).

### Positive Selection on Introgressed Variation Underlies Parallel Adaptive Admixture

By combining *F_adm_* and *LAD* statistics into empirical p-values using Fisher’s method as a composite test of post-admixture positive selection (*52*), we detected significant signals of adaptive admixture in southern-admixed AH populations and evaluated how often these signals overlapped within shared minor-parent ancestry islands (clusters of five or more shared minor-parent ancestry peaks separated by 200 kb) across hybridising species. Applying a conservative Bonferroni threshold for significance (*p* = 0.00035), we first identified several candidate loci for adaptive admixture, including genome-wide evidence for positive selection in multiple southern-admixed populations (Figure 5A).

To identify positively selected genomic regions within shared minor-parent ancestry peaks in multiple taxa, we first defined intervals of ‘minor-parent ancestry islands’ with clusters of shared loci within 200 kb regions for each ancestry (retaining only intervals with five or more shared sites); this resulted in 83 candidate genomic regions with 38, 14 and 31 regions in *A. pectinata*, Taxon 3, and *A. hyacinthus*, respectively. We then defined genomic candidates for parallel adaptive admixture by examining which intervals contained at least one site with significant Fisher p-values in one or more species (with *LAD* calculated in 200 kb windows; *p* < 0.00035 Bonferroni threshold). This resulted in 38 candidate regions (45.8%) for parallel adaptive admixture, comprising 5/38 (13.2%) regions with shared *A. pectinata* minor-parent ancestry positively selected in Taxon 3 and 9/38 regions (23.7%) positively selected in *A. hyacinthus*. There were 3/14 (21.4%) candidate genomic regions with shared Taxon 3 ancestry positively selected in *A. pectinata* and 4/14 regions (28.6%) positively selected in *A. hyacinthus*. There were 7/31 regions (22.6%) for shared minor-parent *A. hyacinthus* ancestry under positive selection in *A. pectinata*, 8/31 regions (25.8%) positively selected in Taxon 3, and also 2/31 regions (6.45%) with significant Fisher p-values in *A. hyacinthus*. Out of these 38 regions, 29 mapped to unique (non-overlapping) chromosomal regions, out of which 8/29 showed evidence for selection in two or more species (Figure 5A; denoted by an asterisk).

The distribution of LD among sites within 29 candidate genomic regions for parallel adaptive admixture were significantly different compared to random subsets of ancestry-informative sites or minor-parent ancestry peaks (Kolmogorov-Smirnov Test, *p* < 1.54 x 10^-17^ for all species; Table S4). These candidate regions also showed elevated proportions of strongly linked sites (e.g., r^2^ > 0.6; Figure S18) and clusters of SNPs in high LD with other candidate loci on other chromosomes, including regions where the same ancestry is under positive selection (Figure S19). Only nine out of 29 candidate genomic regions for parallel adaptive admixture overlapped with SNPs contributing to differentiation between admixed and non-admixed southern populations inferred from PCA-based scans of selection (Figure S7), suggesting that both positive selection on introgressed variation and novel selective sweeps on standing genetic variation represent two pathways for adaptation in corals. No candidate regions overlapped with *f_d_* outliers for *A. millepora* introgression, suggesting that adaptive introgression in these positively regions in admixed populations is not associated with *A. millepora* ancestry. Instead, *f_d_* peaks appeared to coincide with regions lacking ancestry informative sites, consistent with shared *A. millepora* ancestry in AH species within these genomic regions.

Functional annotation and gene ontology assignments of gene regions involved in parallel adaptive admixture revealed signatures of positive selection across functionally diverse genes (Table S5). We detected signatures of positive selection on a heat shock protein on Chromosome 8 with signatures of positive selection on Taxon 3 ancestry (selected in *A. pectinata* and 4; Fisher *p* < 0.00035) (Figure S20). We also detected a prominent signal of positive selection on Chromosome 8 (Fisher *p* < 0.00035) spanning an enoyl-CoA hydratase-isomerase family gene, where *A. hyacinthus* ancestry has risen to high frequency across numerous SNPs in all admixed species and coincides with parallel F_ST_ deviations between admixed and non-admixed populations (Figure 5B). Another positively selected region on Chromosome 4 lies within ∼300 kb of the *Sacsin* heat shock molecular chaperone (Figure S21), a previously identified thermal-tolerance candidate in *Acropora* corals (*62*).

## Acknowledgements

We acknowledge the Traditional Owners of the Great Barrier Reef for giving their free, prior and informed consent to conduct this research on the Sea Countries. We pay our respects to elders, past, present and future. We thank the Australian Institute of Marine Science and the ecoRRAP team for supporting and organising field expeditions and CSIRO for their assistance with molecular laboratory work and genomic sequencing. We thank Gus Crosbie for field assistance to collect reference specimens and Sage Rassmussen for taxonomic assistance. This work was supported by the Reef Restoration and Adaptation Program, funded by the partnership between the Australian Government’s Reef Trust and the Great Barrier Reef Foundation.

## Data Availability Statement

Data files and supporting code will be deposited on the University of Queensland Library eSpace and on Github (https://github.com/ivapops/Acropora_hybridisation). Sequence data will be deposited on the NBCI Short Read Sequence Archive and associated metadata will be available on GEOME.

---

**Figure S1.**
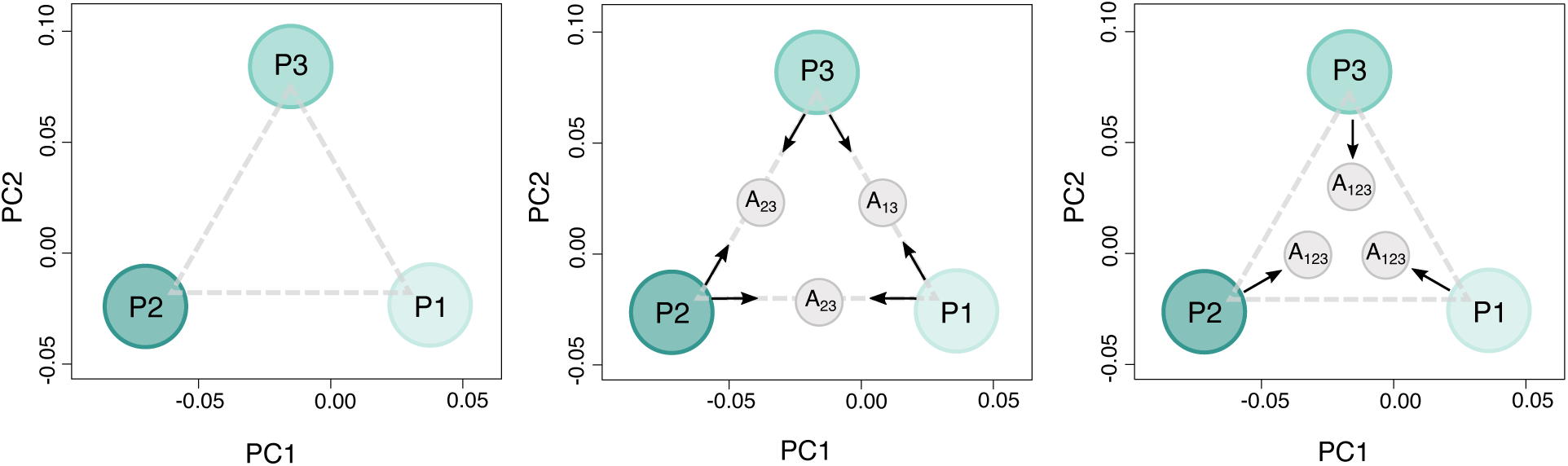
Schematic illustrating genomic patterns in two primary PC axes that are consistent with three-way admixture. **Left panel:** Hypothetical dataset with three hybridising parental populations (P1, P2, P3). **Middle panel:** Admixed populations between any two parental populations fall along the line of the triangle between hybridising taxa. **Right panel:** Under three-way admixture, admixed populations share genetic ancestry with all three parental populations and are expected to be pulled towards each other inside the triangle.

**Figure S2.**
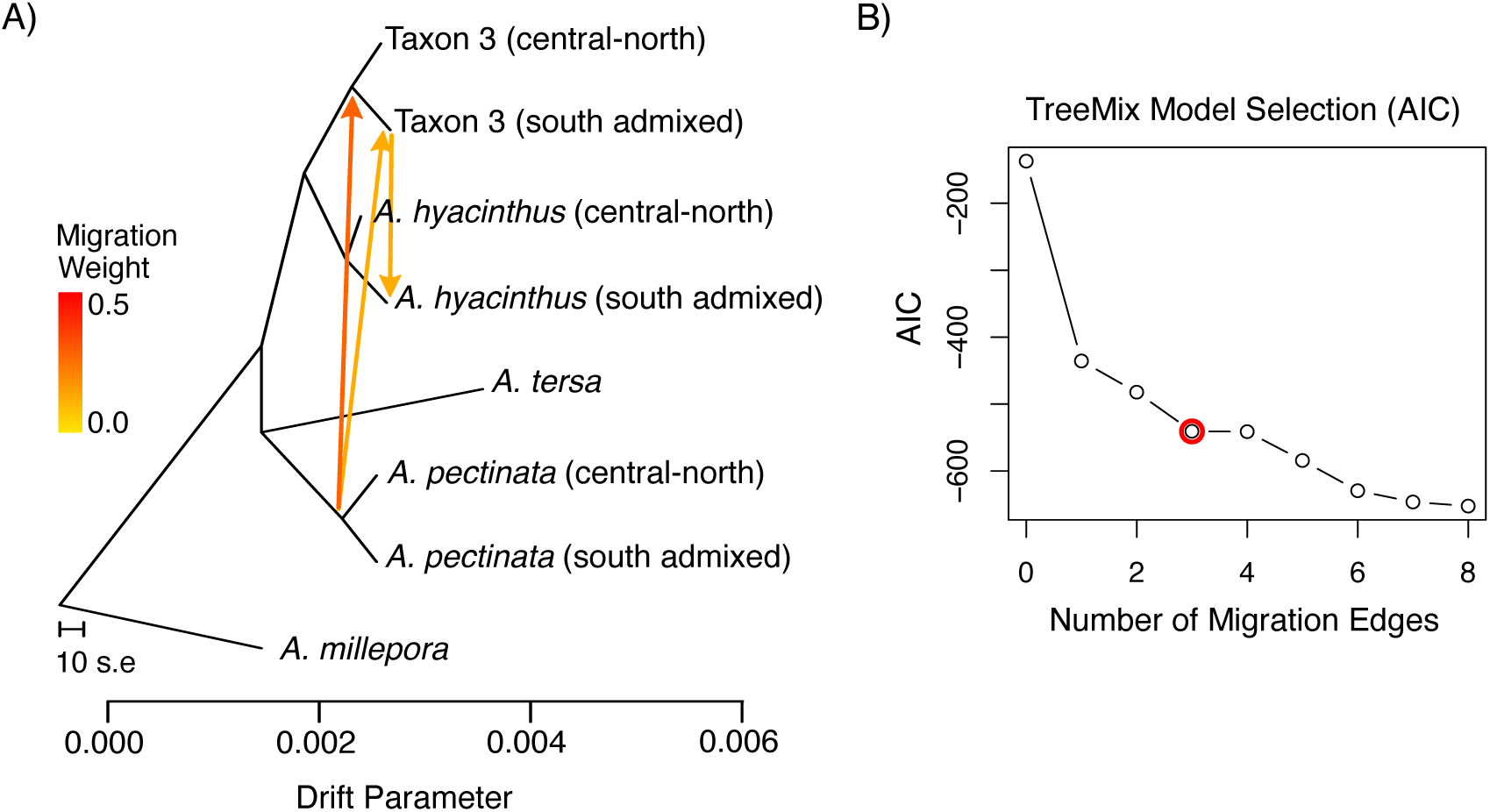
TreeMix admixture graph assigning colonies to genetically distinct southern admixed AH populations and non-admixed parental-type AH populations (northern and central Great Barrier Reef), with *A. millepora* as an outgroup. **(A)** Maximum likelihood population tree with three migration edges, which returned the highest likelihood tree (based on AIC comparisons) and explained > 98% of the variation; **(B)** Stepwise comparison Akaike information criterion (AIC) between sequential migration models supported three migration edges, as the difference between nested models with four migration edges was less than two (ΔAIC > 2) averaging across 10 replicate runs.

**Figure S3.**
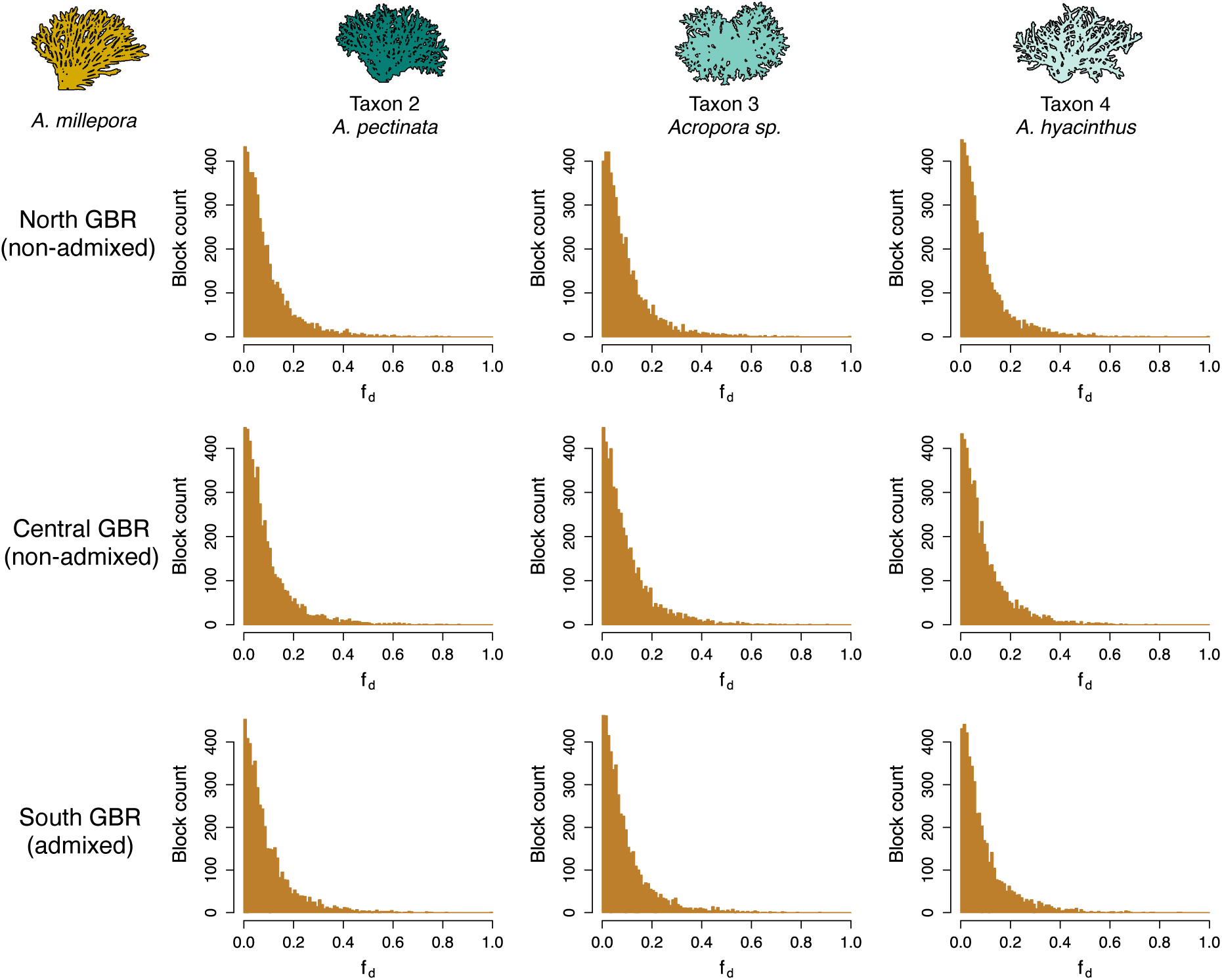
Genome-wide distributions of *f_d_* values showing the fraction of introgression (calculated in 50 kb sliding windows) between *A. millepora* and three focal AH species in the northern, central and southern Great Barrier Reef. For all regional comparisons, negative *f_d_* values and values exceeding 1 were excluded. Genome-wide *f_d_* values were centred around zero with few genomic regions showing extreme values within the top 1% that might implicate adaptive introgression.

**Figure S4.**
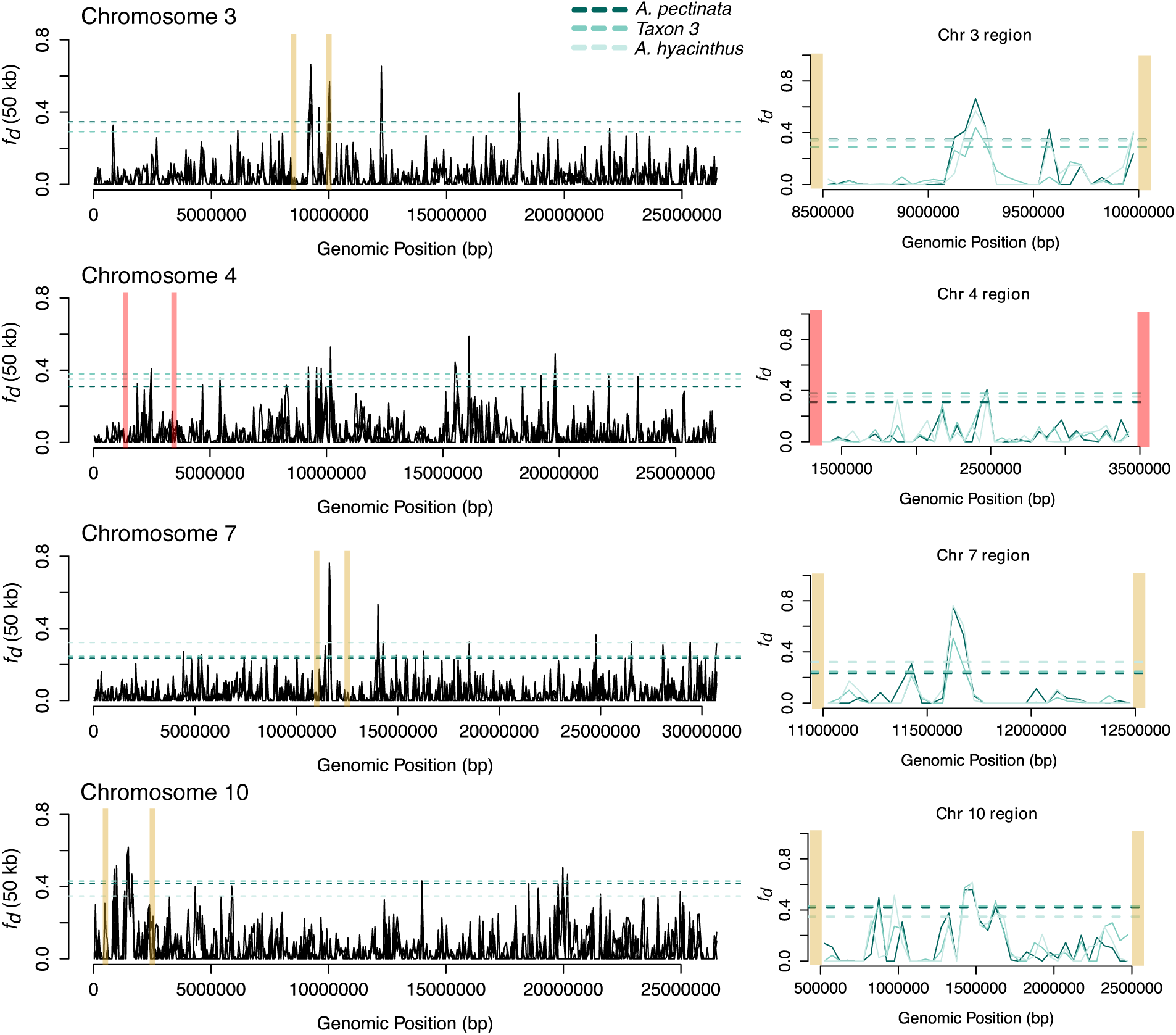
Genome scans for selected chromosomes showing the fraction of introgression *f_d_* (calculated in 50 kb sliding windows) between *A. millepora* and three focal AH species in the southern Great Barrier Reef (northern and central data not shown). Highlighted genomic regions correspond to insets on the right side, showing an example of *f_d_* peaks within the top 1% of the empirical distribution. *f_d_* outlier thresholds are indicated by the dashed lines coloured by species according to the legend. The chromosomal region corresponding to the *A. millepora* HES1 locus (Rose et al. 2021) is show between red lines.

**Figure S5.**
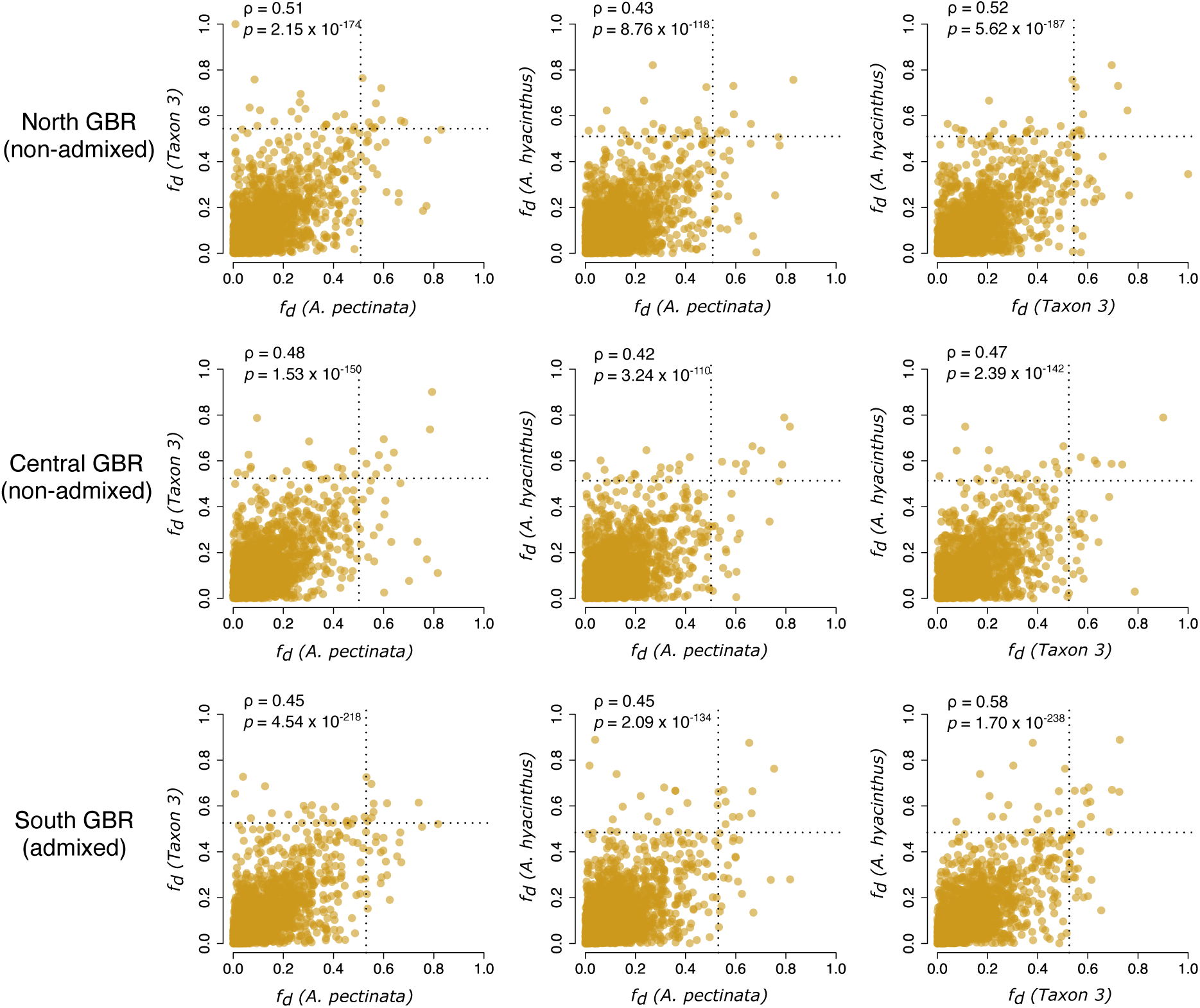
Paired *f_d_* correlations showing the fraction of introgression (calculated in 50 kb sliding windows) between *A. millepora* and AH species in the northern, central and southern Great Barrier Reef. Black dotted lines indicate the top 0.01% of the *f_d_* distribution, excluding negative *f_d_* values and values exceeding 1.

**Figure S6.**
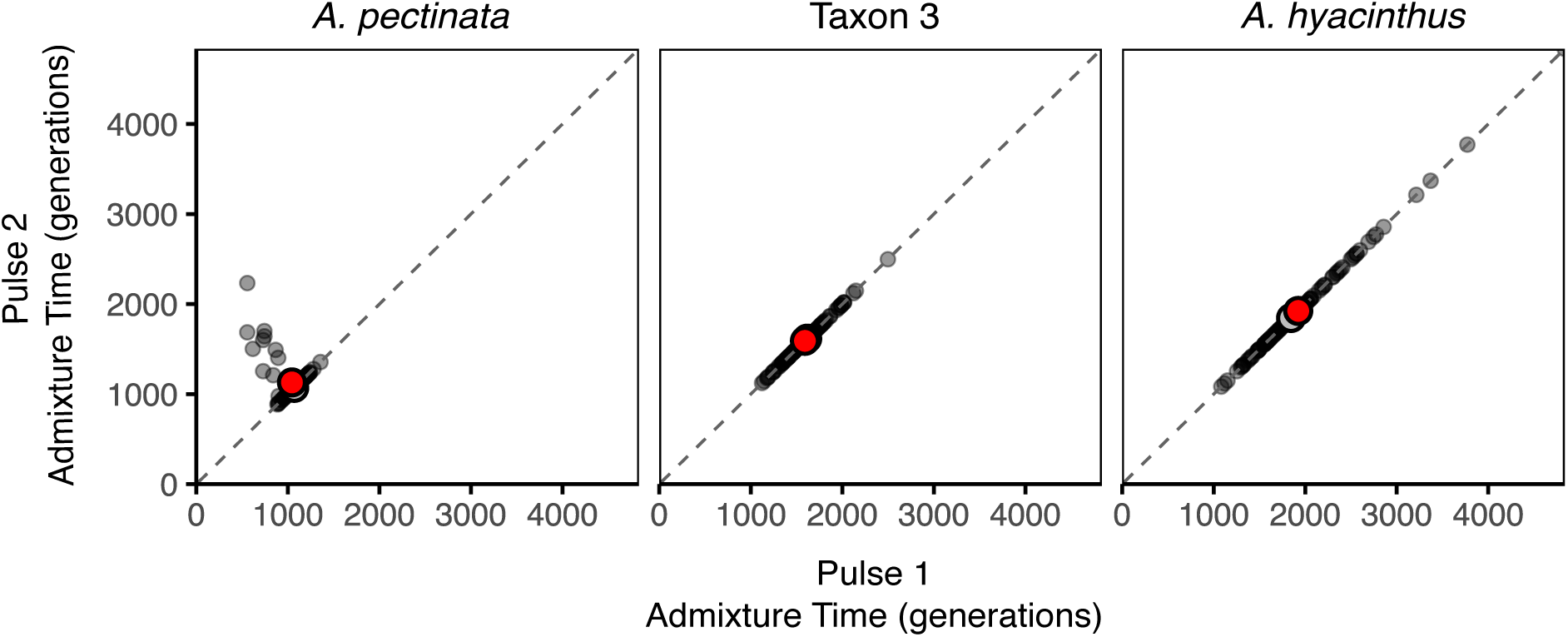
Admixture time estimates inferred from local ancestry inference in Ancestry_HMM using 100 bootstrap replicates (block size 5000 SNPs). The mean and optimum admixture times as inferred by Ancestry_HMM are indicated by large grey and red circles, respectively. Analyses consistently returned similar optimum admixture times between ∼1000-2000 generations for all southern admixed populations.

**Figure S7.**
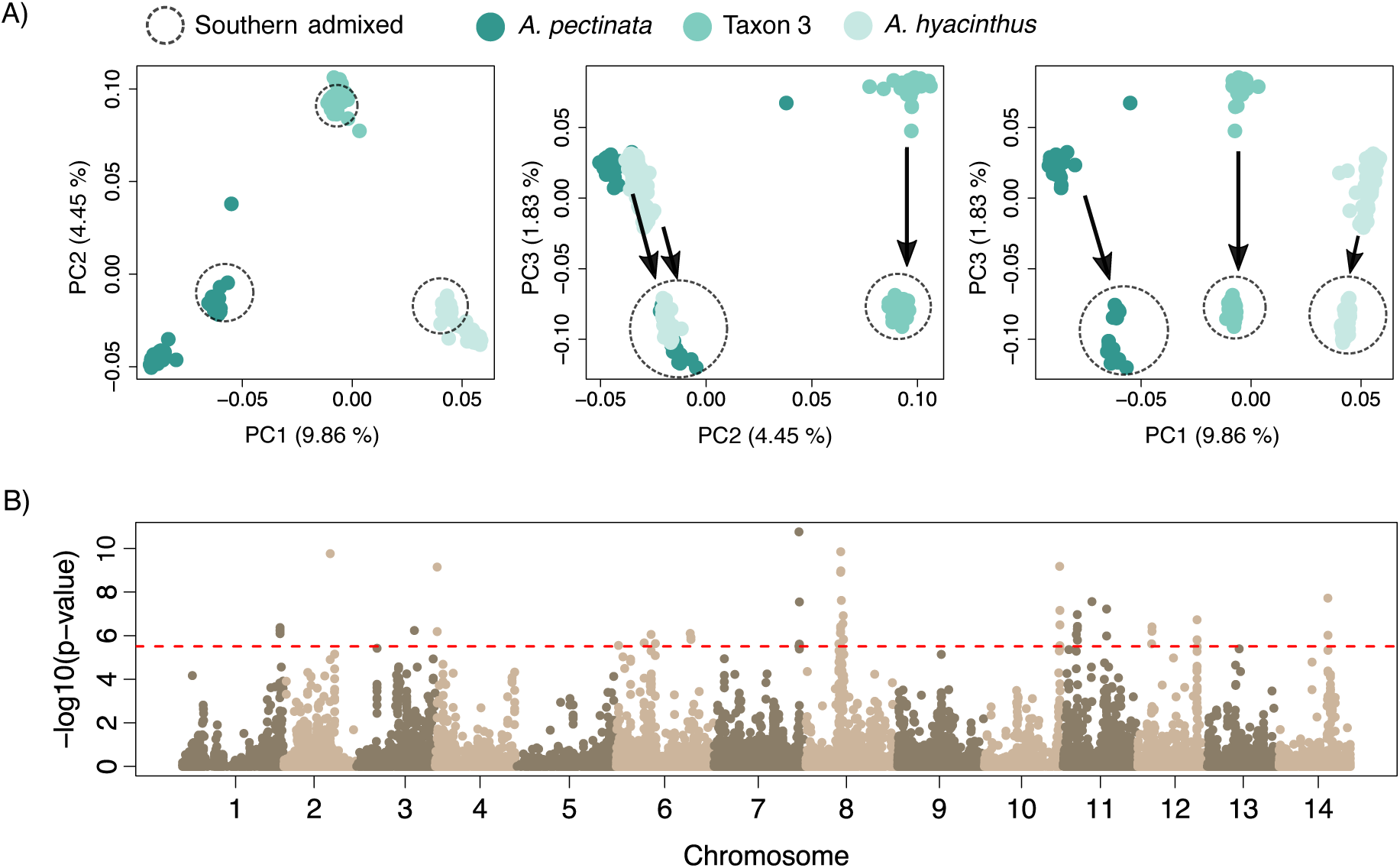
Analysis of parallel genetic differentiation among three focal AH species: *A. pectinata*, Taxon 3 and *A. hyacinthus*; **(A)** PCA showing southern admixed AH populations diverging in parallel from non-admixed parental-type populations on PC3; **(B)** Genome-wide differentiation scan using the *-selection* flag in PCAngsd. Loci falling within the top 0.01% of the empirical -log10(p-value) distribution (dashed red line) were considered as outliers contributing significantly to genomic differentiation between southern admixed and non-admixed taxa on PC3.

**Figure S8.**
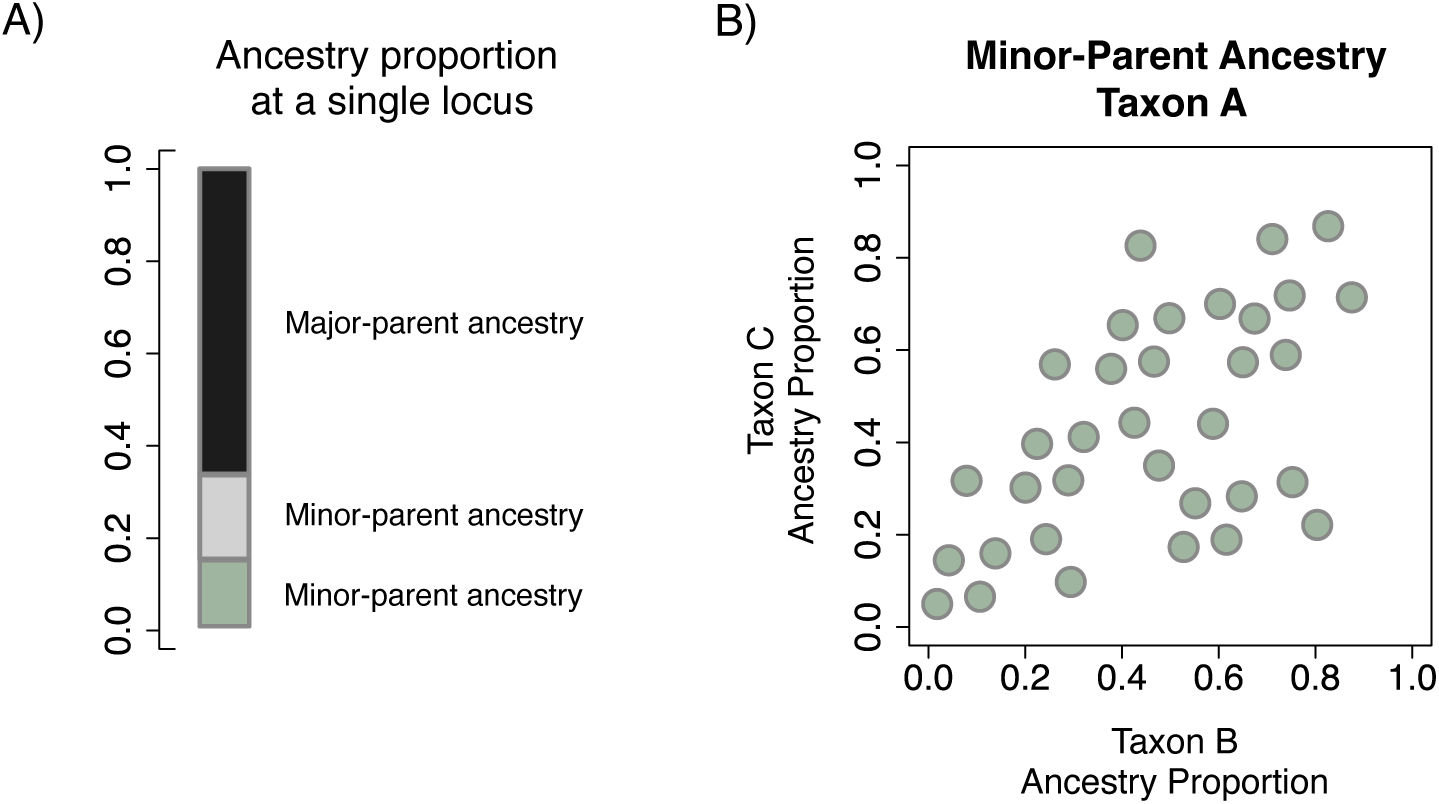
Schematic of local ancestry in the context of three-way admixture. **(A)** Three possible ancestries at a single locus in a hybrid genome. The major-parent ancestry comprises the larger proportion of the hybrid genome and is typically derived from the receiving species (i.e., major-parent genomic background). In a three-way admixture scenario, there are two possible minor-parent ancestries, which comprise the smaller proportion of the hybrid genome at a single locus and are typically derived from the introgressing species. **(B)** Hypothetical example of genome-wide positive correlations in the same minor-parent ancestry (derived from Taxon A; green in (A)) in the major-parent genomic background of Taxon B and Taxon C. Loci exceeding 50% minor-parent ancestry proportions in both Taxon B and Taxon C are considered shared minor-parent ancestry peaks.

**Figure S9.**
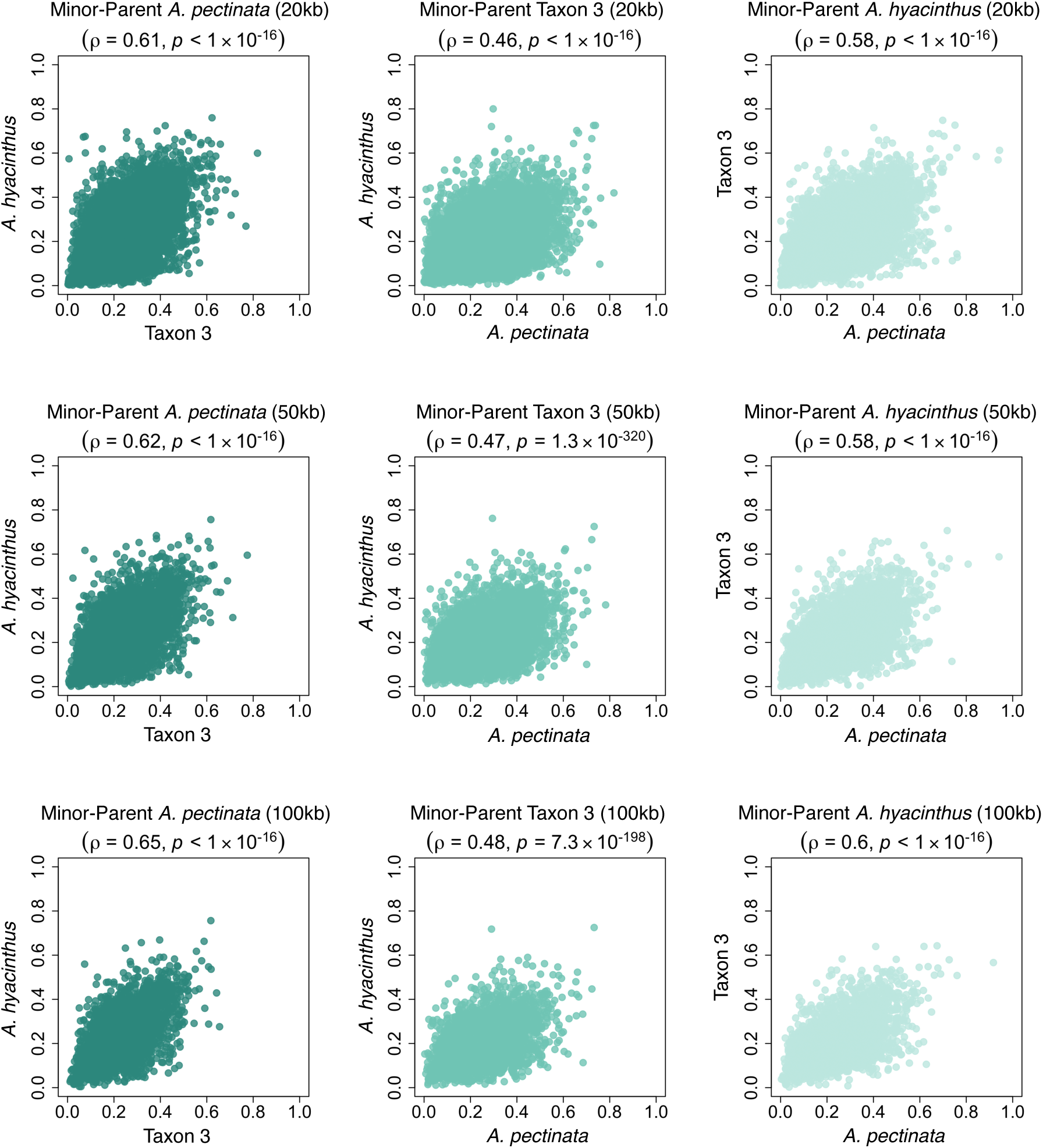
Genome-wide minor-parent ancestry correlations in three admixed AH species. The same minor-parent ancestry is positively correlated among hybridising taxon pairs considering average ancestry proportions within 20-100 kb windows (Spearman’s *ρ* range 0.46-0.65; *p* < 1 x 10^-16^; 200 kb data shown in Figure 4).

**Figure S10.**
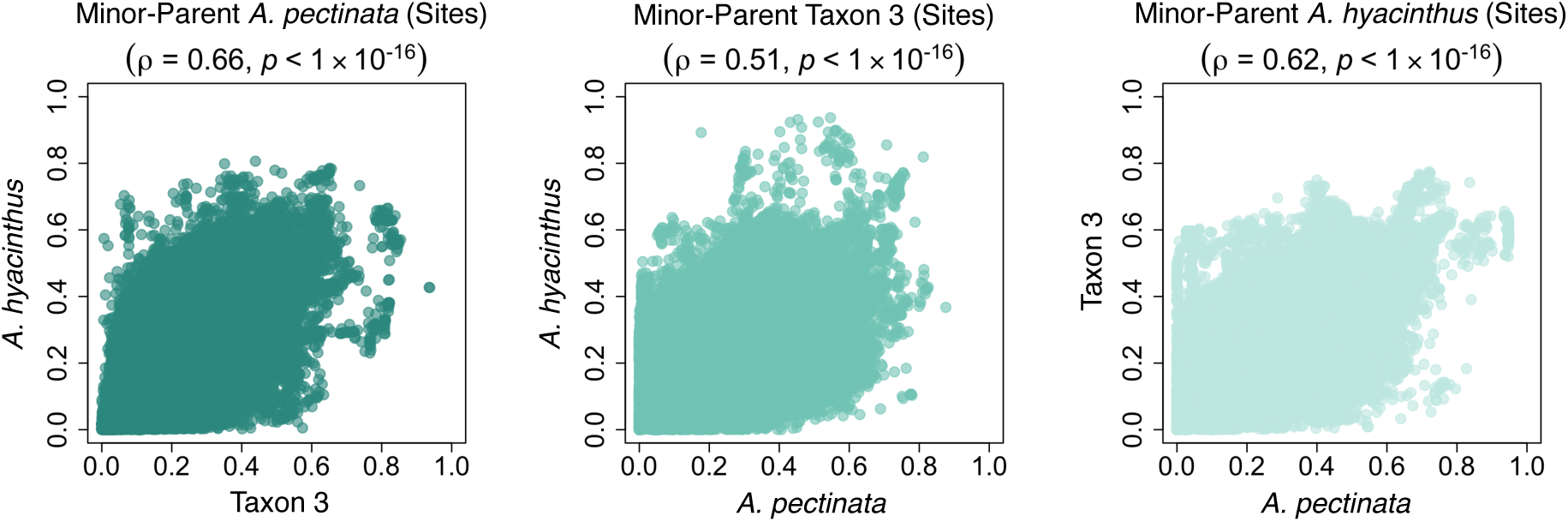
Genome-wide minor-parent ancestry correlations in three admixed AH species. The same minor-parent ancestry is positively correlated among hybridising taxon pairs (Spearman’s *ρ* range 0.51-0.66; *p* < 1 x 10^-16^) considering ancestry informative sites.

**Figure S11.**
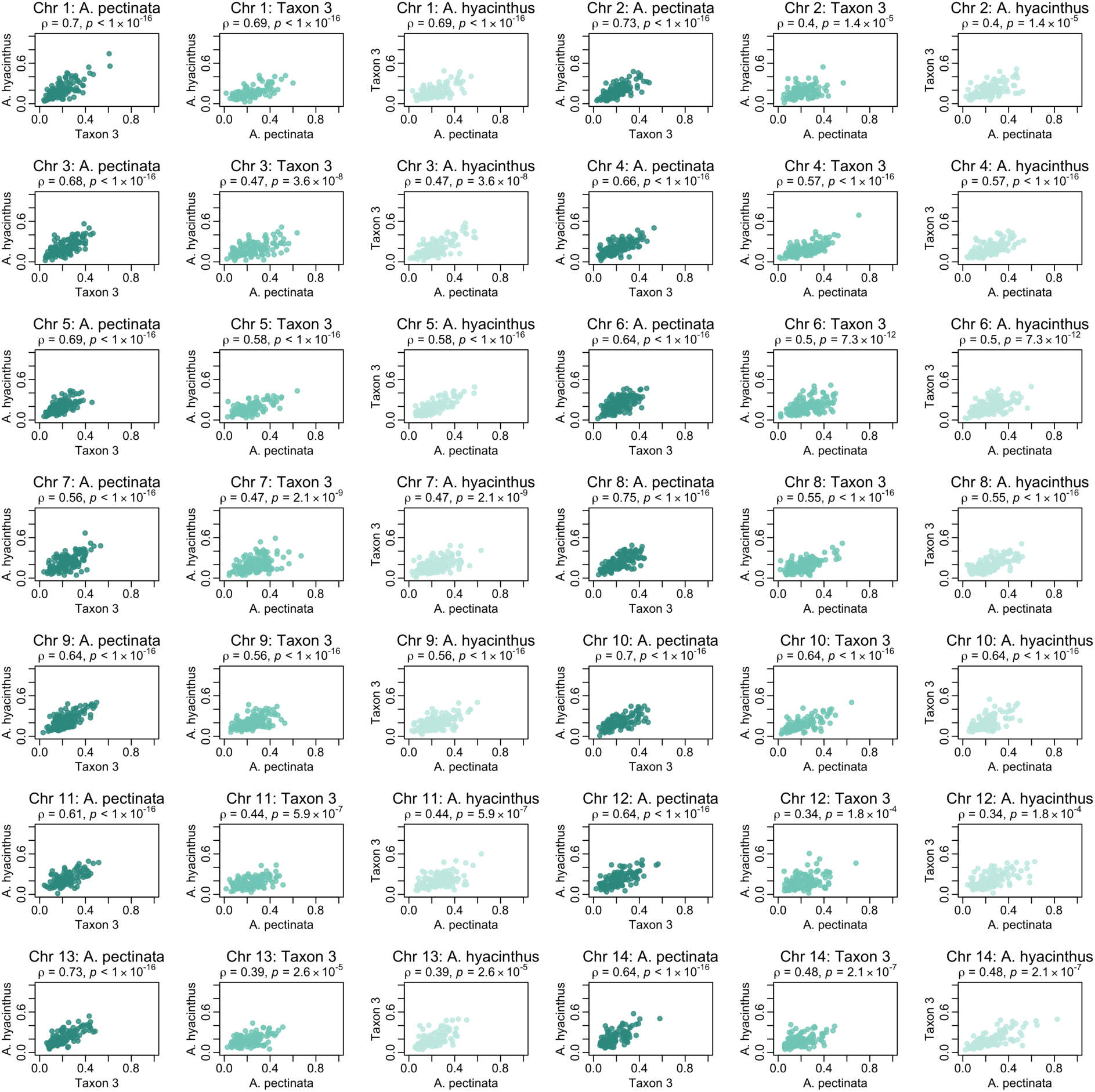
Minor-parent ancestry correlations in three admixed AH species per chromosome. The same minor-parent ancestry is positively correlated among hybridising species pairs on each chromosome (Spearman’s *ρ* range 0.34-0.75; *p* < 2.6 x 10^-05^) considering average ancestry proportions within 200 kb windows. Minor-parent ancestry correlations on all chromosomes were consistently positive considering ancestry informative sites and average ancestry within windows between 20-100 kb (not shown).

**Figure S12.**
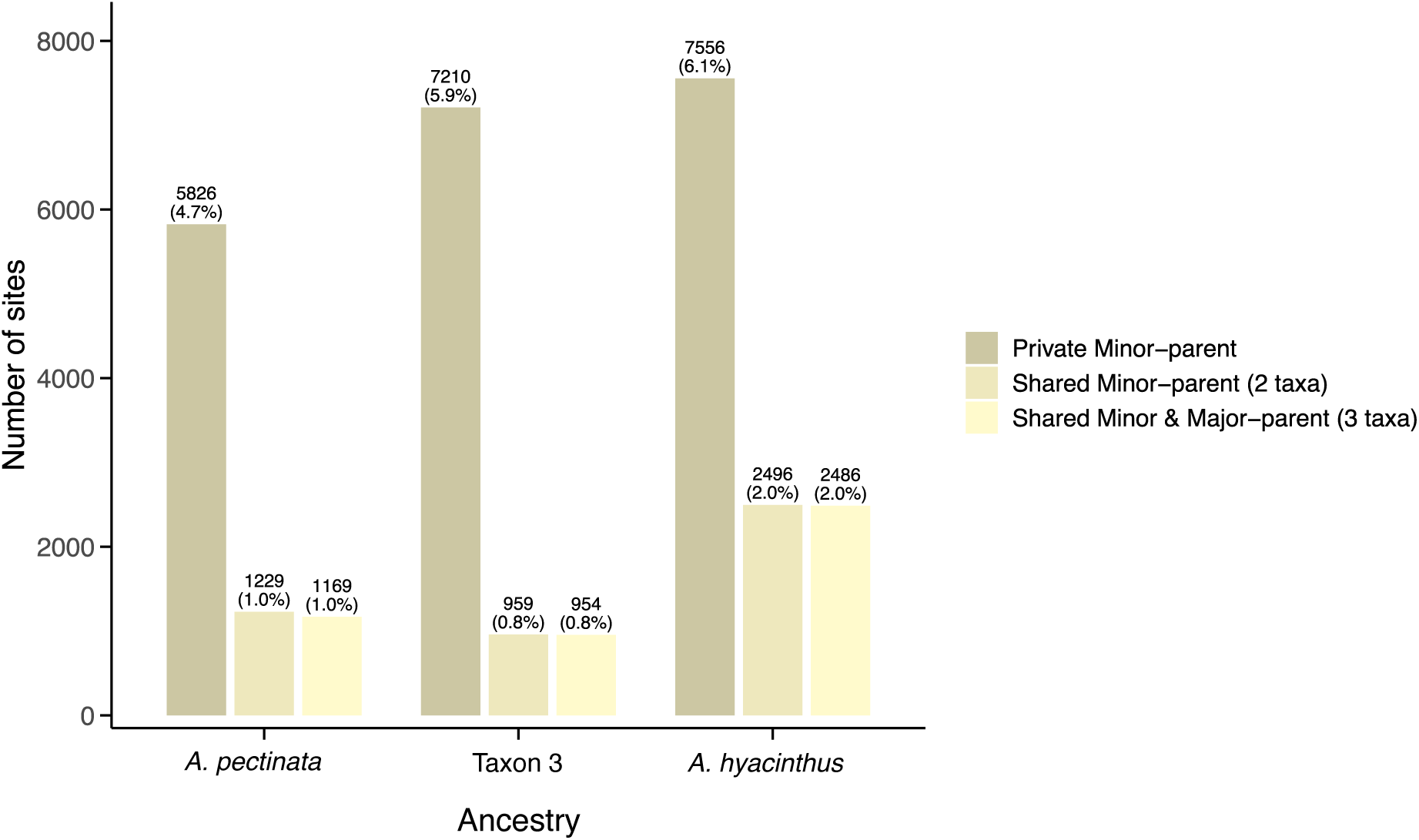
Counts of sites with minor-parent ancestry peaks (ancestry proportions exceeding 50%) for species-specific ancestry across different major-parent backgrounds. The proportion of ancestry informative sites is shown in brackets. Out of 122,977 common ancestry informative sites, an average of 5.6% of sites were private (non-shared) minor-parent ancestry peaks, 1.3% of sites showed shared minor-parent ancestry peaks in two taxa. Of these, the majority of loci (>95%) also showed major-parent ancestry peaks at the same loci.

**Figure S13.**
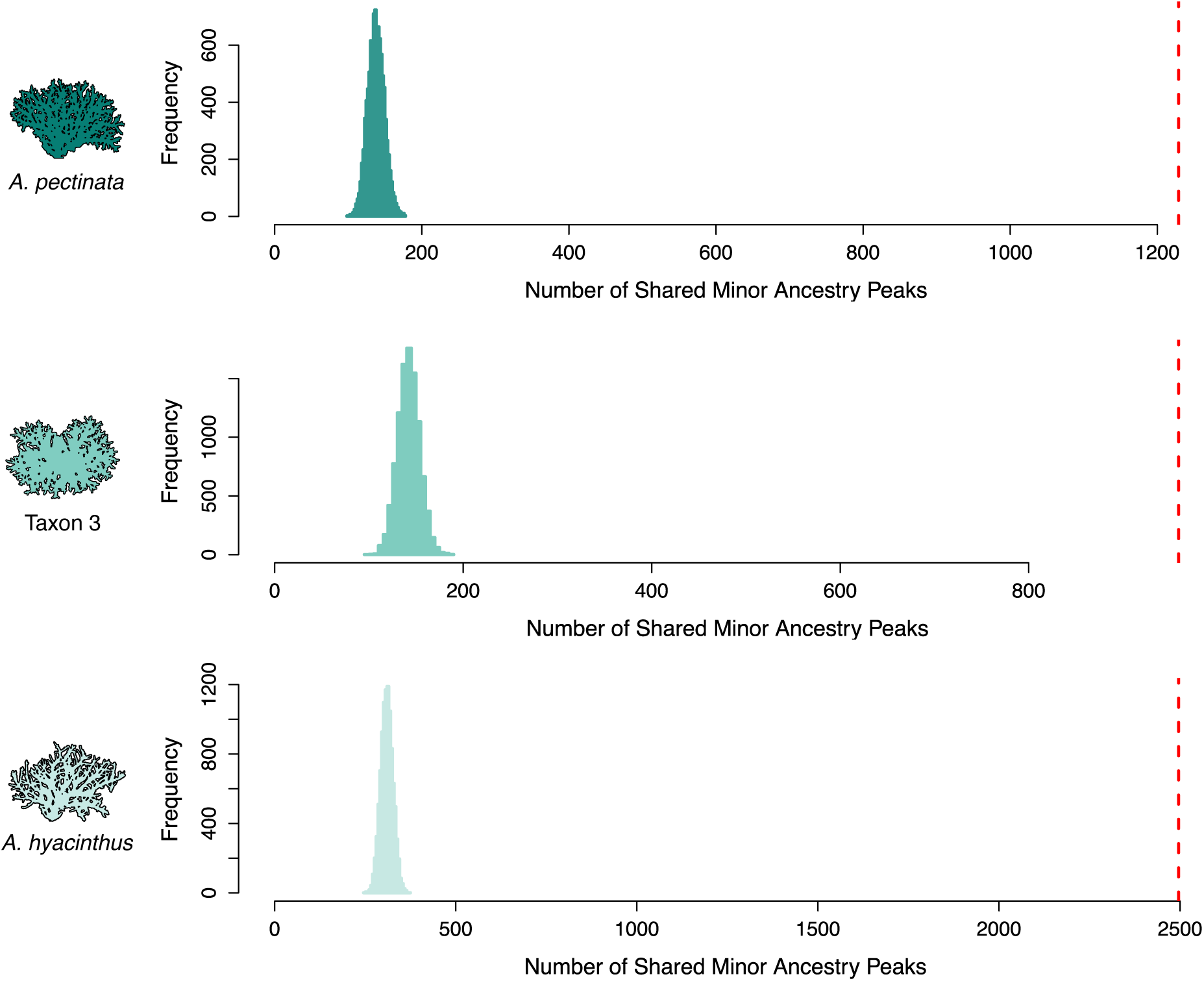
Distributions of shared minor-parent ancestry peaks (sites > 50% ancestry proportions) expected by chance based on 10,000 permutations of the data. For each AH species, the number of observed shared minor-parent ancestry peaks (dashed red line) exceeded sharing excepted by chance considering ancestry informative sites (*p* < 9.9 x 10^-5^) and mean ancestry calculated in genomic windows of 20 kb (*p* < 9.9 x 10^-5^), 50 kb (*p* < 9.9 x 10^-5^), 100 kb (*p* < 9.9 x 10^-5^) and 200 kb (*p* < 4.9 x 10^-4^) (plots not shown).

**Figure S14.**
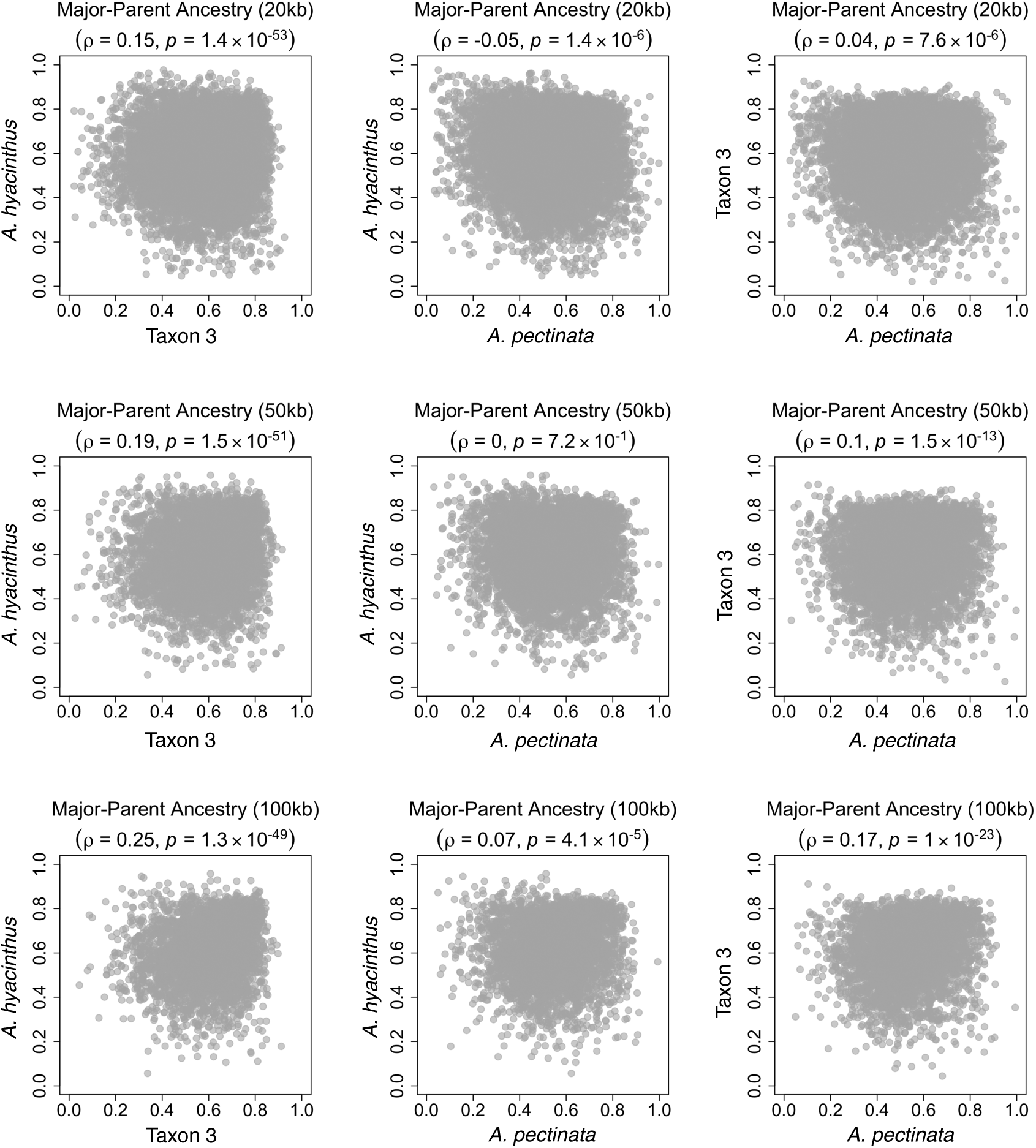
Genome-wide major-parent ancestry correlations in three admixed AH species. Major-parent ancestry correlations were weak among hybridising species pairs (Spearman’s *ρ* range: -0.05-0.25; *p* < 7.2 x 10^-1^) considering average ancestry proportions within 20-100 kb windows (200 kb shown in Figure 4).

**Figure S15.**
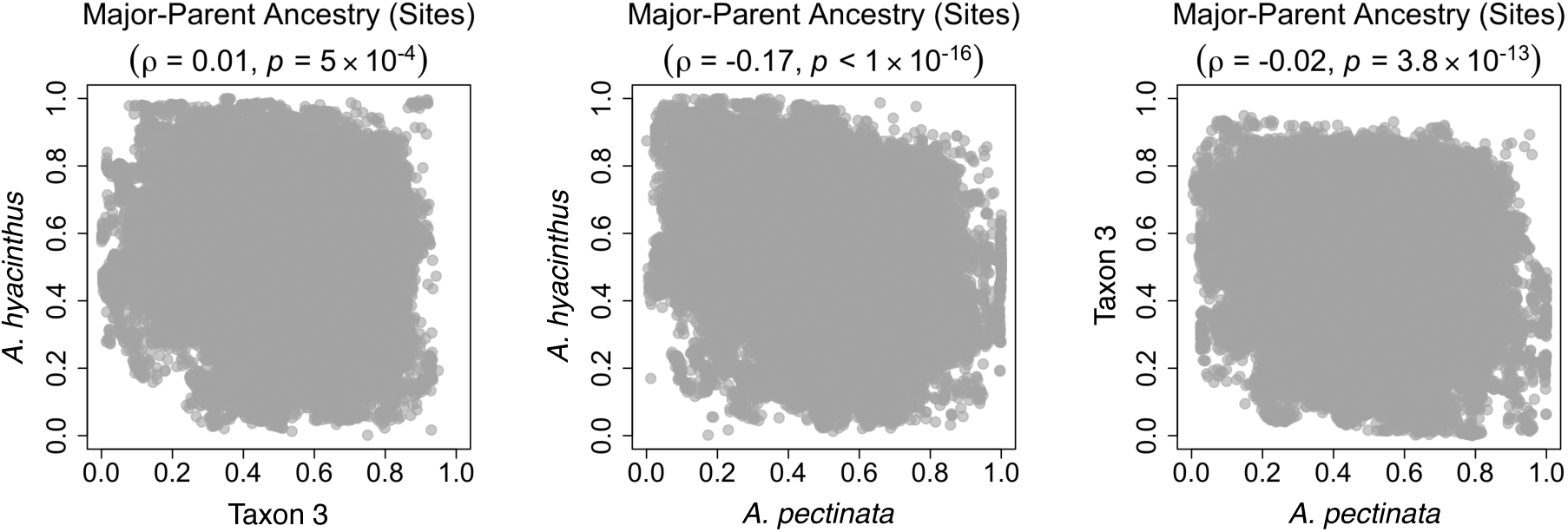
Genome-wide major-parent ancestry correlations among three admixed AH species. Major-parent ancestry correlations were weak among hybridising species pairs (Spearman’s *ρ* range: -0.17-0.01; *p* < 1 x 10^-16^) considering ancestry informative sites.

**Figure S16.**
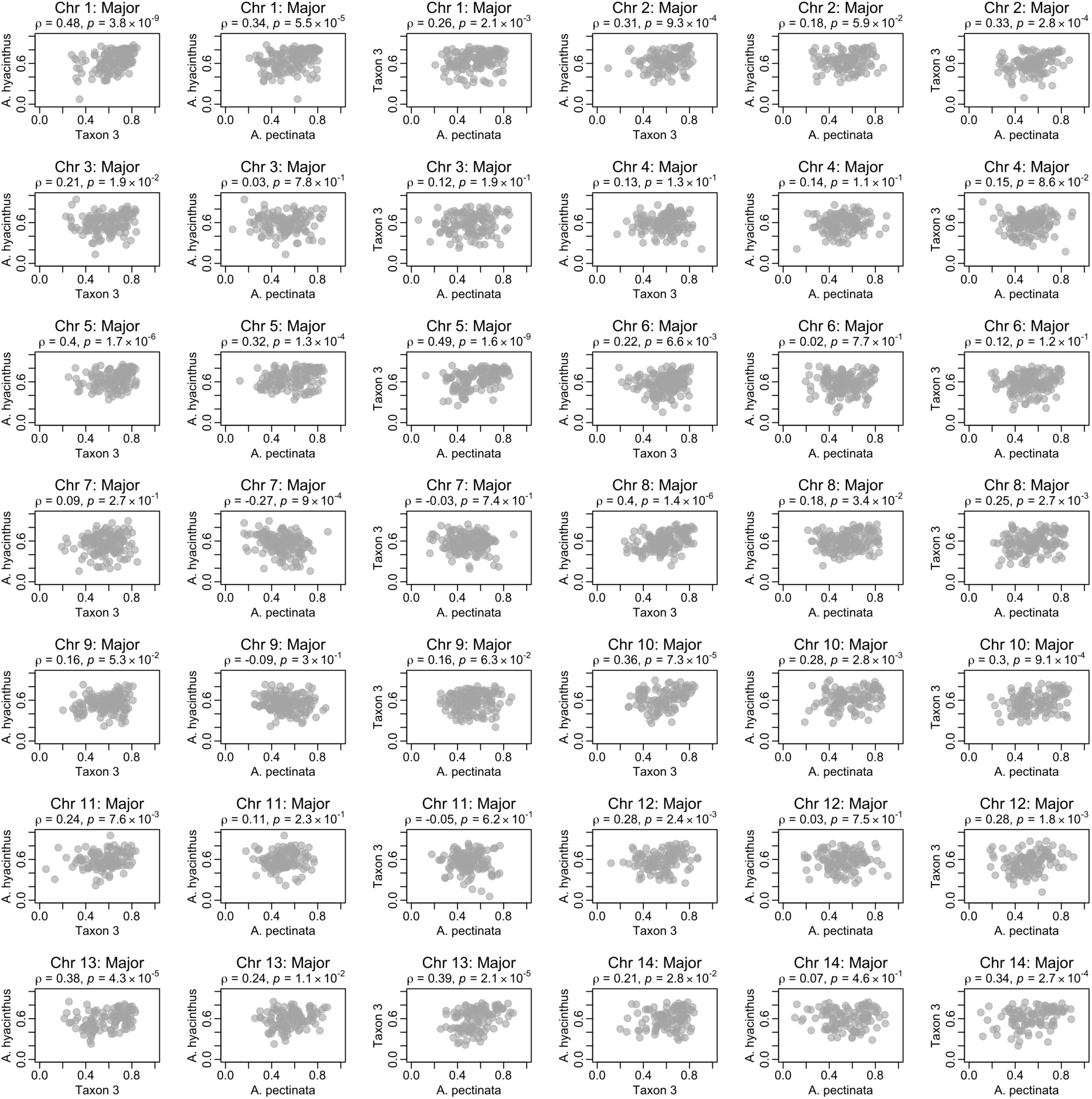
Major-parent ancestry correlations among three admixed AH species per chromosome. The major-parent ancestry correlations among hybridising species pairs were generally weak across chromosomes, considering average ancestry proportions within 200 kb windows (average ancestry within windows between 20-100 kb not shown).

**Figure S17.**
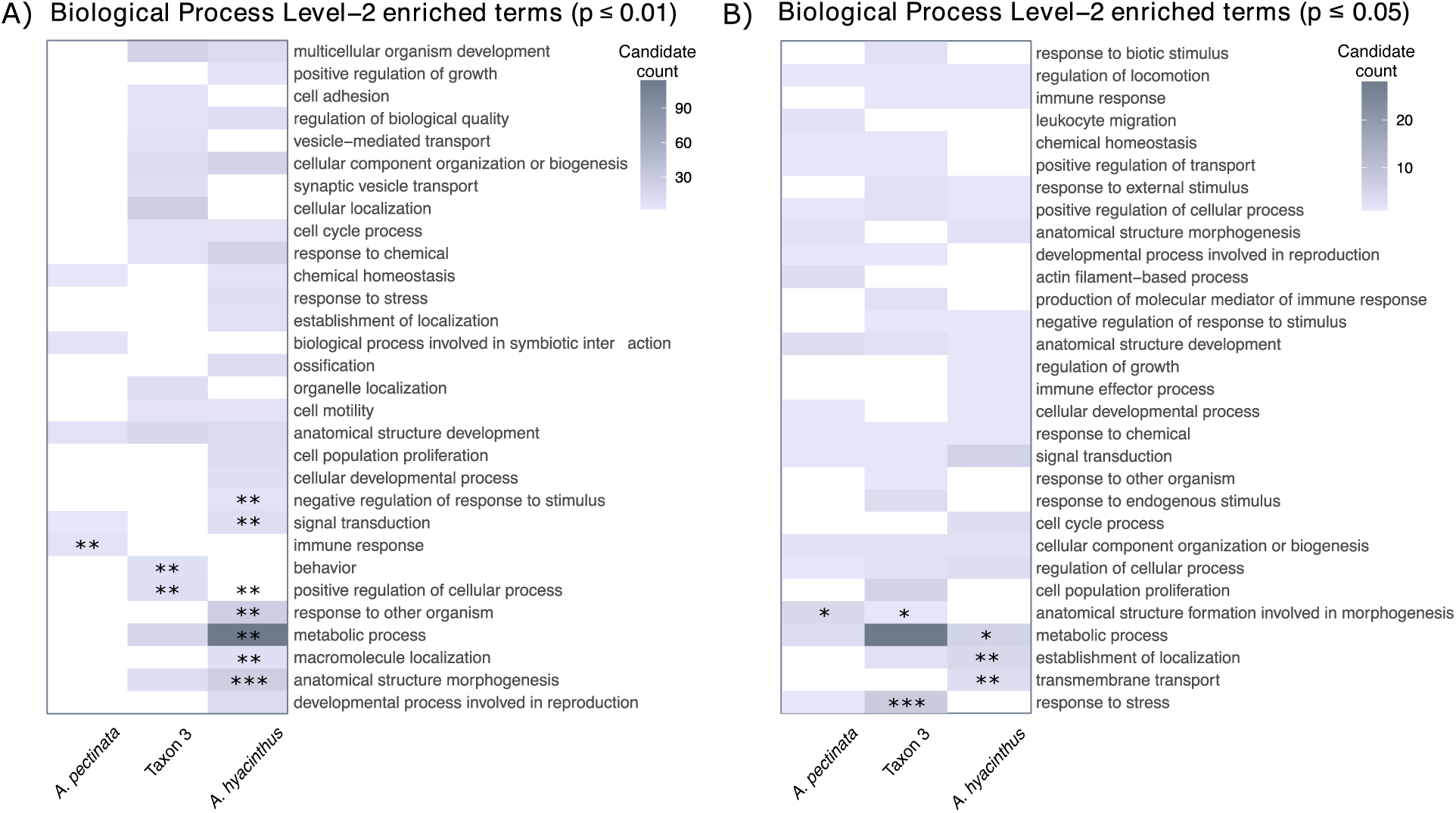
Top 30 significantly enriched Level-2 category GO-terms. **(A)** Significantly enriched GO-terms (*p* ≤ 0.01) in private (non-shared) minor-parent ancestry peaks (> 50% ancestry proportion in a single taxon), with significance of GO-term enrichment indicated with *p* ≤ 0.0001*** and *p* ≤ 0.001**; **(B)** Significantly enriched GO-terms (*p* ≤ 0.05) in shared minor-parent ancestry peaks (minor-parent ancestry > 50% in two taxa), with significance of GO-term enrichment indicated with *p* ≤ 0.0001***, *p* ≤ 0.001**, and *p* ≤ 0.01*.

**Figure S18.**
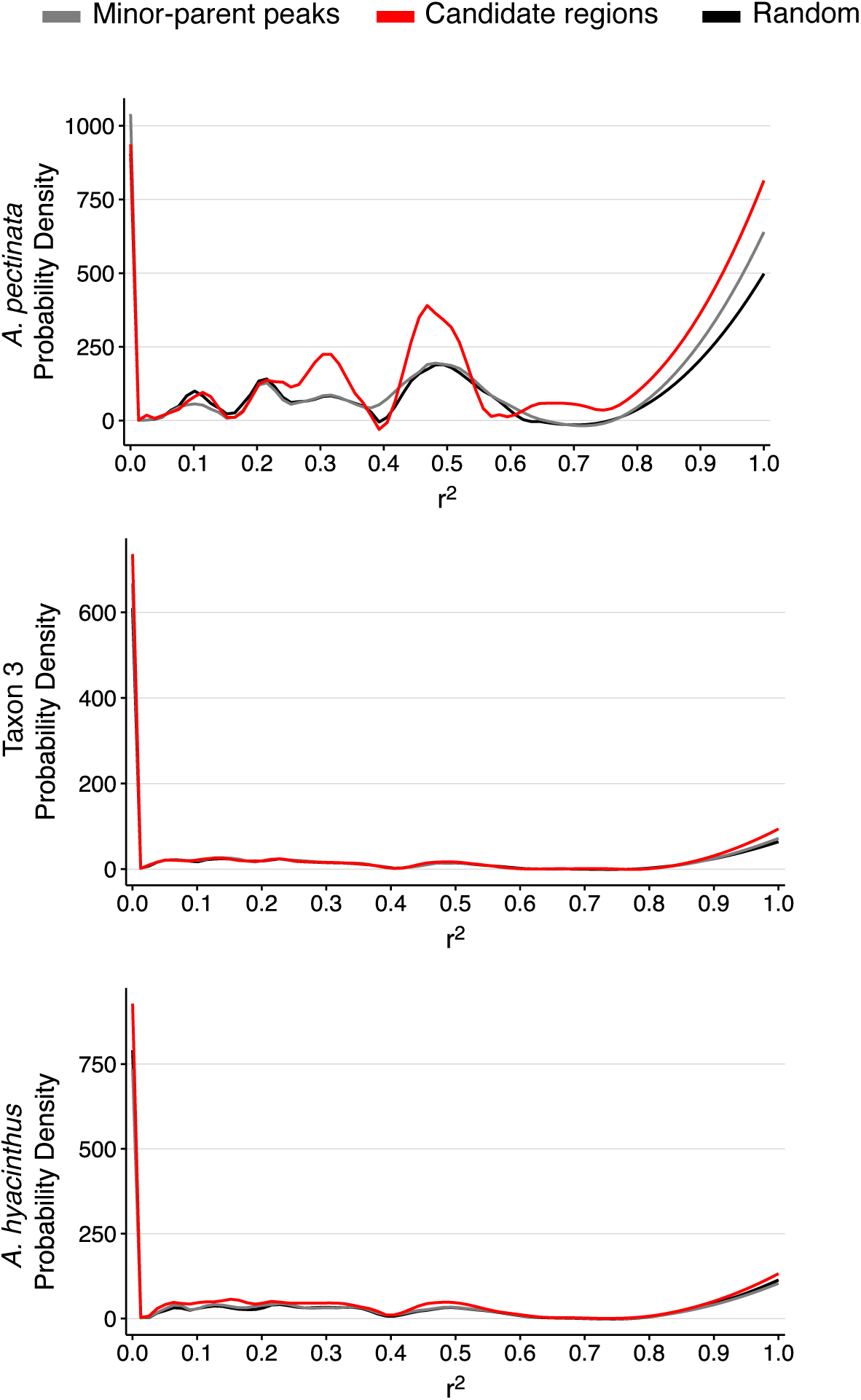
Probability density function curves for linkage disequilibrium (LD; r^2^) for each admixed AH population. Plots show comparisons among three datasets: (Dataset 1) *Minor-Parent Peaks*: sites with private or shared minor-parent ancestry peaks (sites exceeding 50% minor-parent ancestry proportions) for each AH species; (Dataset 2) *Candidate regions:* sites within 29 candidate genomic regions for parallel adaptive admixture (normalised to 200 kb base pairs wide); and (Dataset 3) *Random*: a randomly selected subset of ancestry-informative sites equivalent to the number of minor-parent ancestry peaks (i.e., Dataset 1). Lines represent locally estimated scatterplot (LOESS) smoothed probability density curves with a smoothing span of 0.20 to visualise LD distributions. LD distributions were significantly different among all datasets (Kolmogorov-Smirnov test, *p* < 1.10 x 10^-07^; Table S4), with greater proportions of sites with high linkage values among sites within 29 candidate genomic regions (e.g., r^2^ > 0.6; Figure S19).

**Figure S19.**
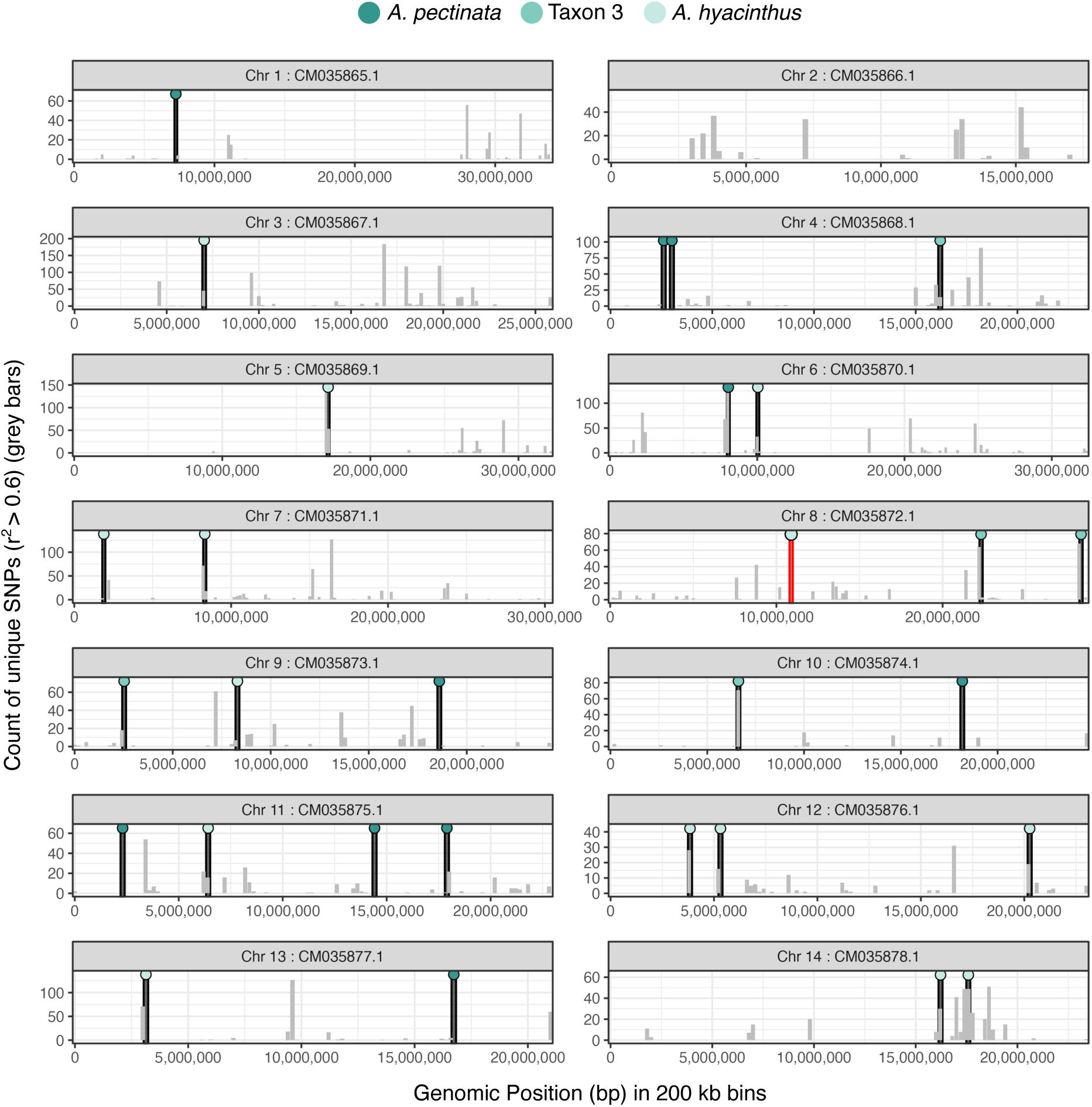
Example linkage network analysis for one candidate region for parallel admixture on Chromosome 8 in the genomic background of *A. pectinata*. At this candidate region (red bar), we detected a prominent signal of positive selection (Fisher *p* < 0.00035), where *A. hyacinthus* ancestry has risen to high frequencies across numerous SNPs in all admixed AH species (Figure 5). Linkage network analysis of sites with minor-parent ancestry proportions exceeding 50% in the genome of *A. pectinata*, where the plots show histograms of counts of unique SNPs in 200 kb bins (grey bars) that are in high LD (r^2^ > 0.6) with at least one site within another candidate region (black bar) for each chromosome. SNPs within 50 kb of the focal candidate region (red bar) are omitted for clarity. Black bars indicate the locations of other candidate regions for parallel admixture normalised to 200 kb intervals, with the selected ancestry denoted by the coloured circle.

**Figure S20.**
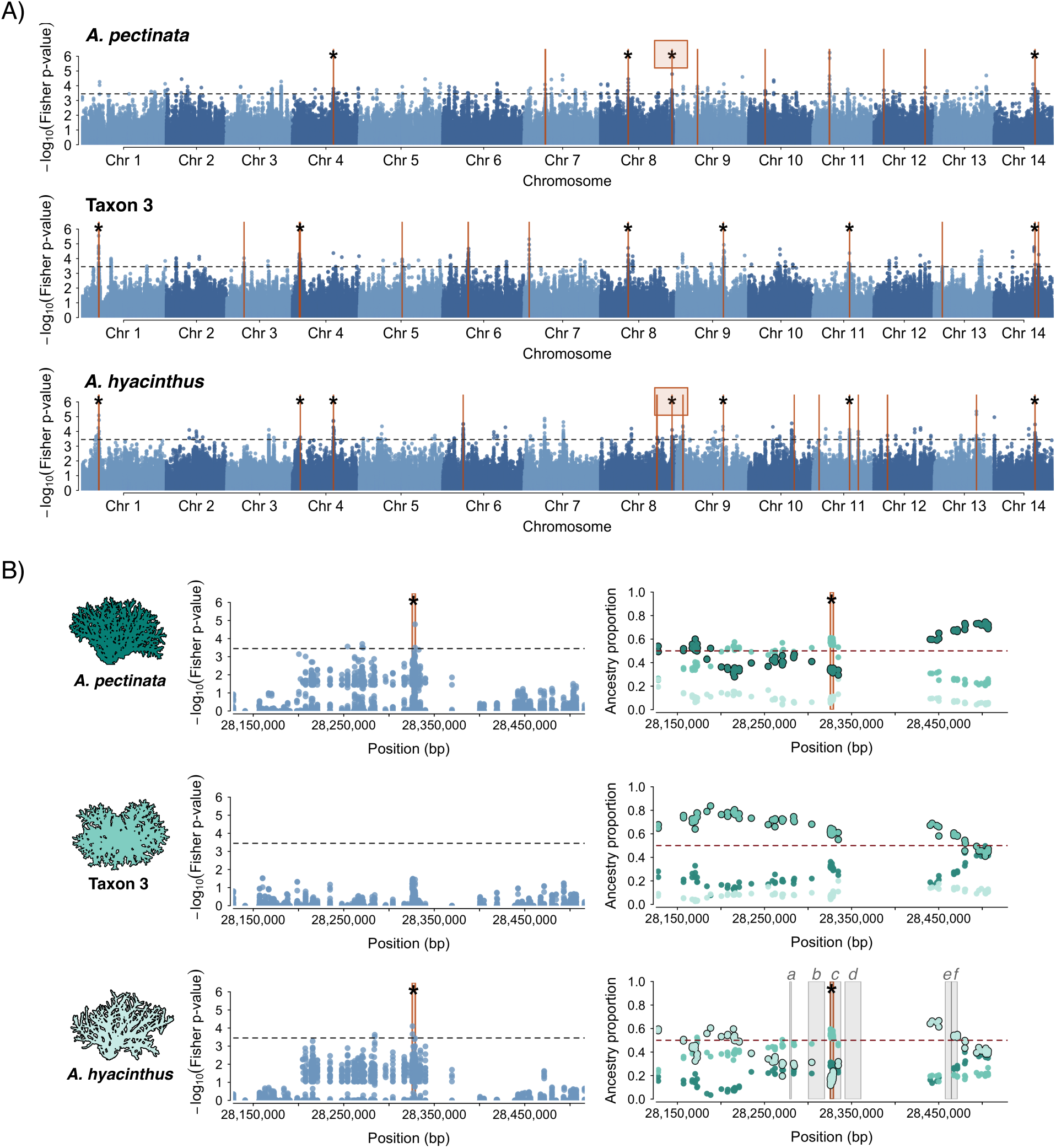
Evidence for positive selection on admixed variation in three hybridising AH species in the southern Great Barrier Reef. **(A)** Genome-wide Fisher p-values, where the black horizonal lines indicate the Bonferroni threshold and orange vertical bars indicate 29 candidate genomic regions for parallel adaptive admixture where signatures of positive selection coincide with shared minor-parent ancestry peaks exceeding 50% ancestry proportions in two AH species. Asterisks indicate candidate genomic regions with evidence for positive selection in two or more species. **(B)** Local signatures of parallel adaptive admixture on a focal Chromosome 8 region (indicated with an orange rectangle in (A) and (B)), where Taxon 3 ancestry has risen to high frequencies across numerous SNPs in all three admixed populations. Species-specific ancestry is indicated by colours shown in the coral icons and major-parent ancestry is outlined in black. This region is adjacent to a heat shock protein (*a*) and is in close proximity to a number of genes with diverse functions (*b, c, d, e, f*; gene functions in Table S5).

**Figure S21.**
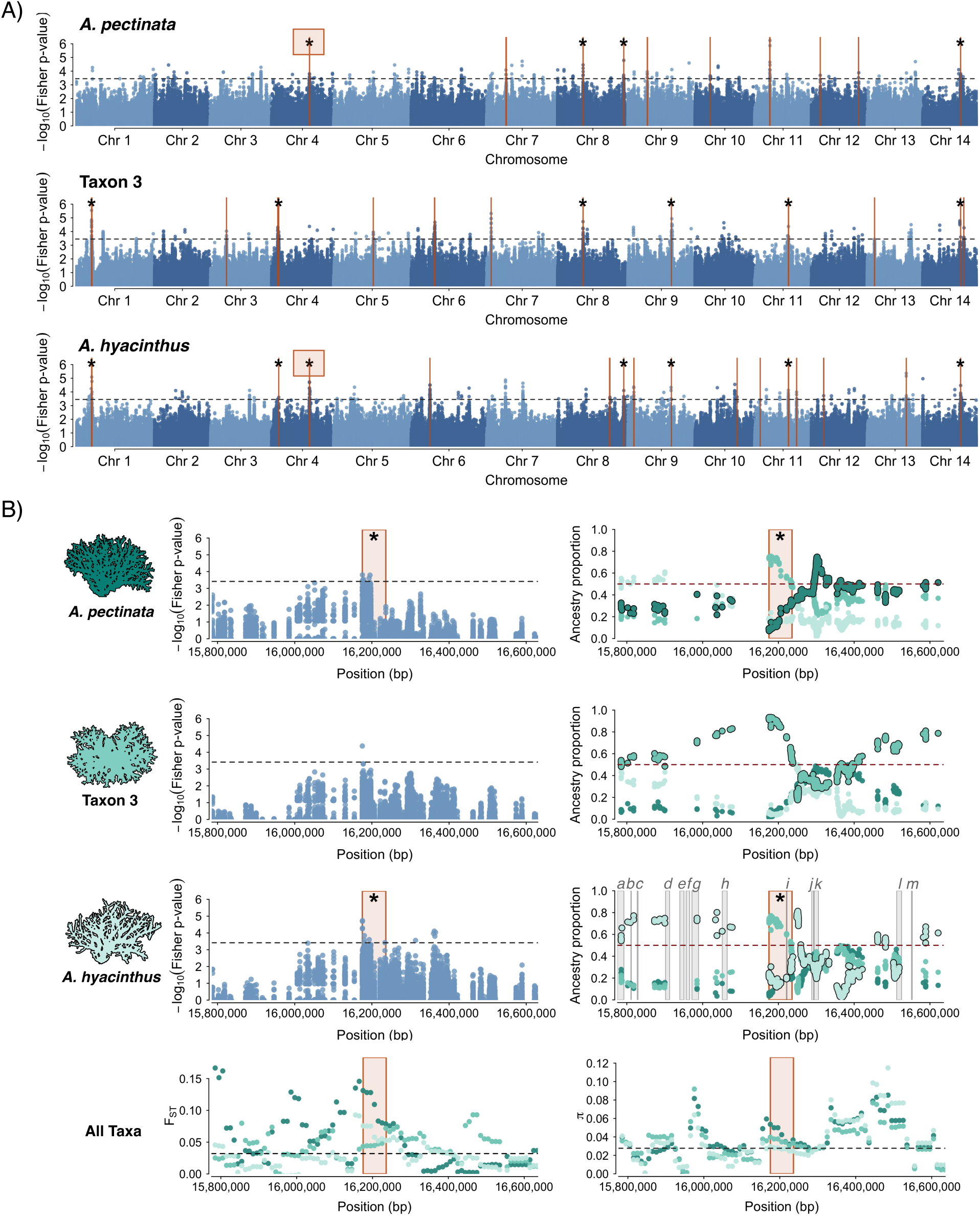
Evidence for positive selection on admixed variation in three hybridising AH species in the southern Great Barrier Reef. **(A)** Genome-wide Fisher p-values, where the black horizonal lines indicate the Bonferroni threshold and orange vertical bars indicate 29 candidate genomic regions for parallel adaptive admixture where signatures of positive selection coincide with shared minor-parent ancestry peaks exceeding 50% ancestry proportions in two AH species. Asterisks indicate candidate genomic regions with evidence for positive selection in two or more species. **(B)** Local signatures of parallel adaptive admixture on a focal Chromosome 4 region (indicated with an orange rectangle in (A) and (B)), where Taxon 3 ancestry has risen to high frequencies across numerous SNPs in all three admixed populations. Species-specific ancestry is indicated by colours shown in the coral icons and major-parent ancestry is outlined in black. This region lies within ∼300 kb of a gene orthologous to the Sacsin heat shock molecular chaperone (*l*) and is in close proximity to a number of genes with diverse functions (*a-m;* gene functions in Table S5).

**Figure S22.**
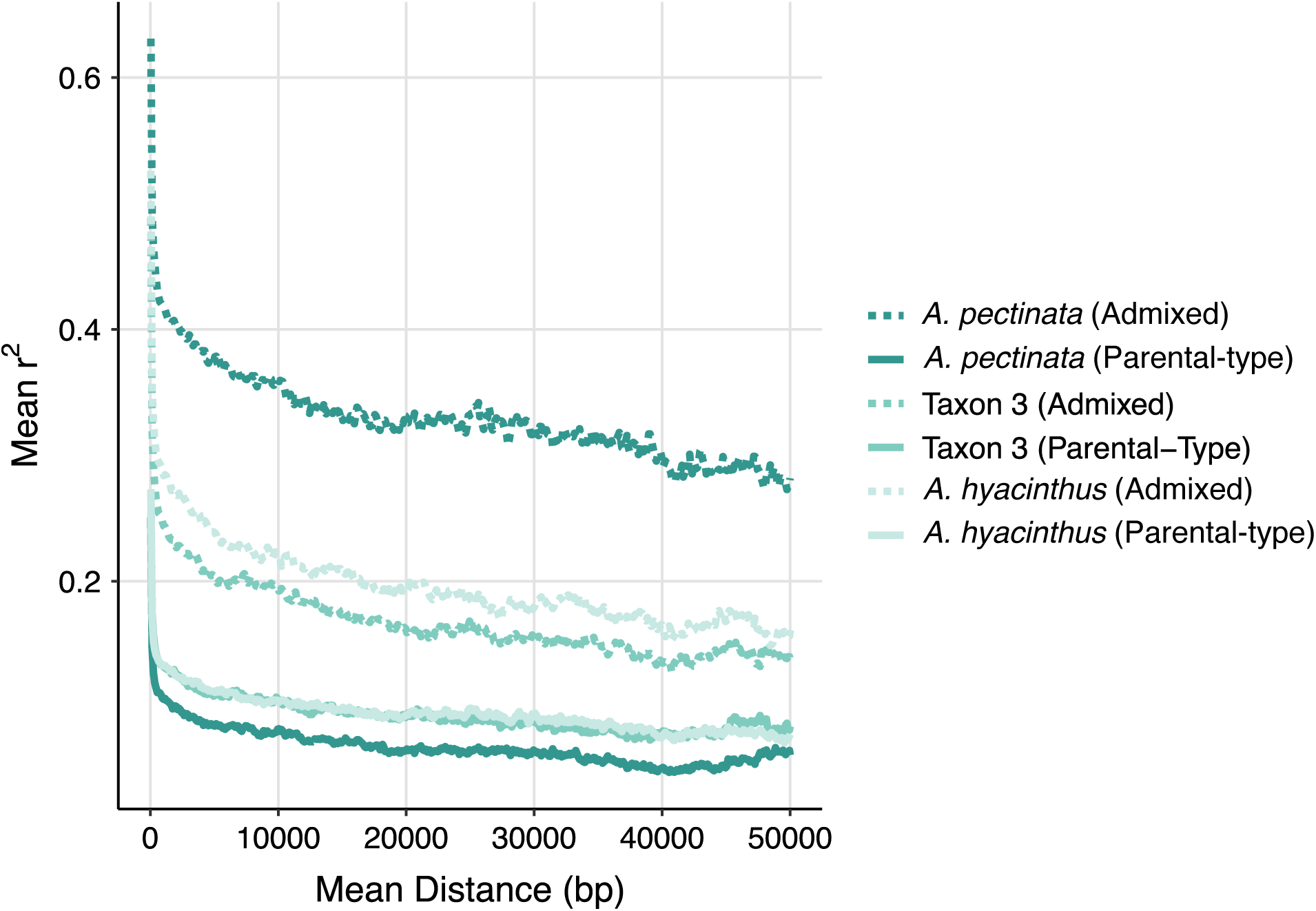
Linkage disequilibrium decay between loci in southern admixed AH populations and non-admixed parental-type populations found in the central and northern Great Barrier Reef.

**Figure S23.**
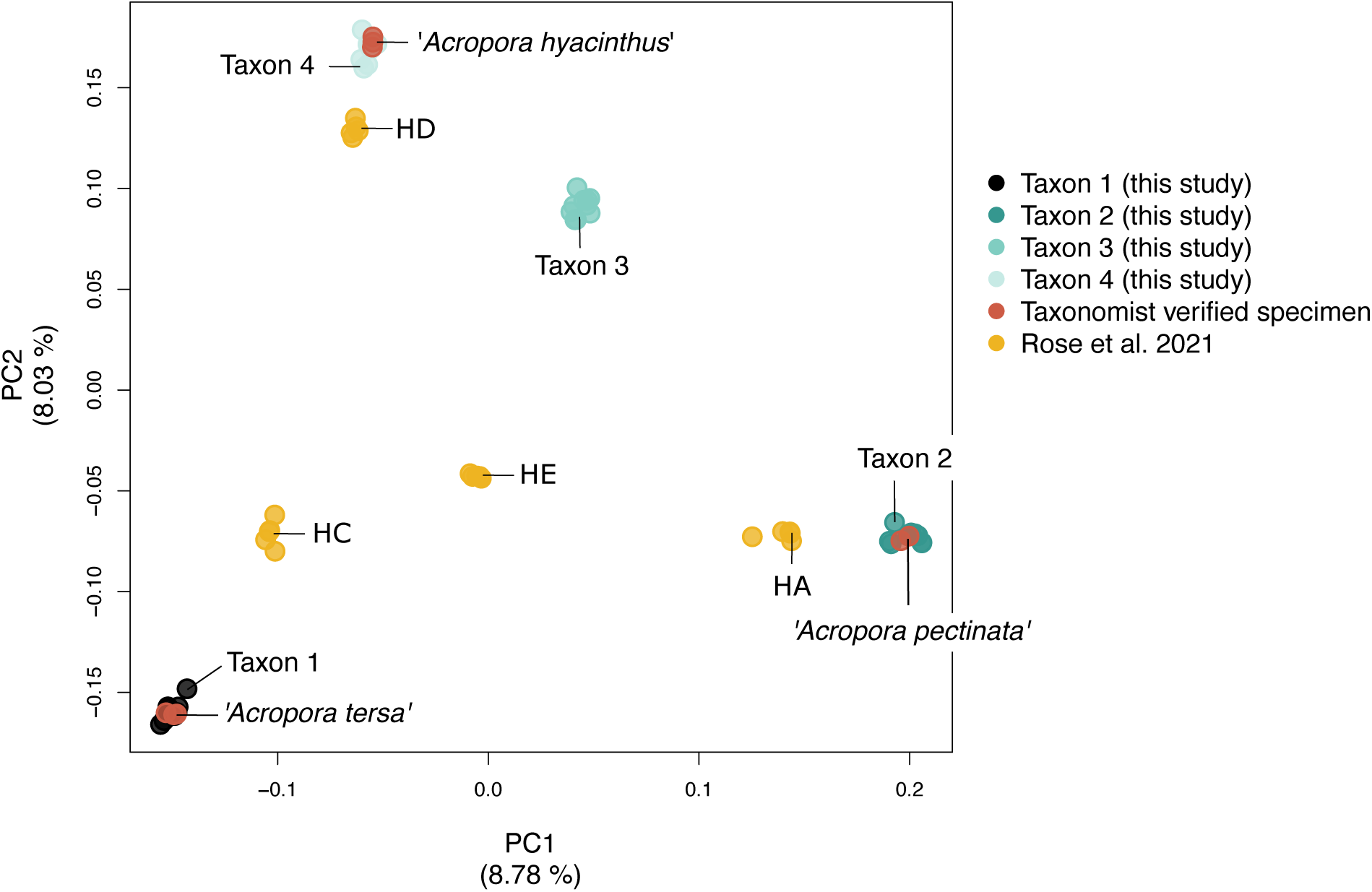
PCA of representative taxa in the *Acropora hyacinthus* species complex (AH) from this study to contextualised the identity of the four AH species in the GBR. We co-analysed ten representative colonies per taxon alongside eight specimens from the Queensland Museum collection identified with reference to type materials and specimens examined by (*1*), as well as 20 AH samples from Ofu, American Samoa (HA, HC, HE and HD; (*2*). Based on the most recent taxonomy (*1*), we confirmed Taxon 1 as *Acropora tersa* (referred to as *A. hyacinthus* “neat” in (*3*) prior to the formal description of *A. tersa* by (*1*), Taxon 2 as *Acropora pectinata* and Taxon 4 as *Acropora hyacinthus*. Taxon 3 represents a currently undescribed species that was not included in (*1*). Analyses with representative specimens belonging to AH species from Ofu, American Samoa (HA, HC, HE and HD; (*2*)) showed close affinities on PC1 and PC2 between Taxon 1 and HC, Taxon 2 and HA, Taxon 4 and HD. Taxon HE did not show clear affinities to AH species in the GBR (Figure S23).

**Figure S24.**
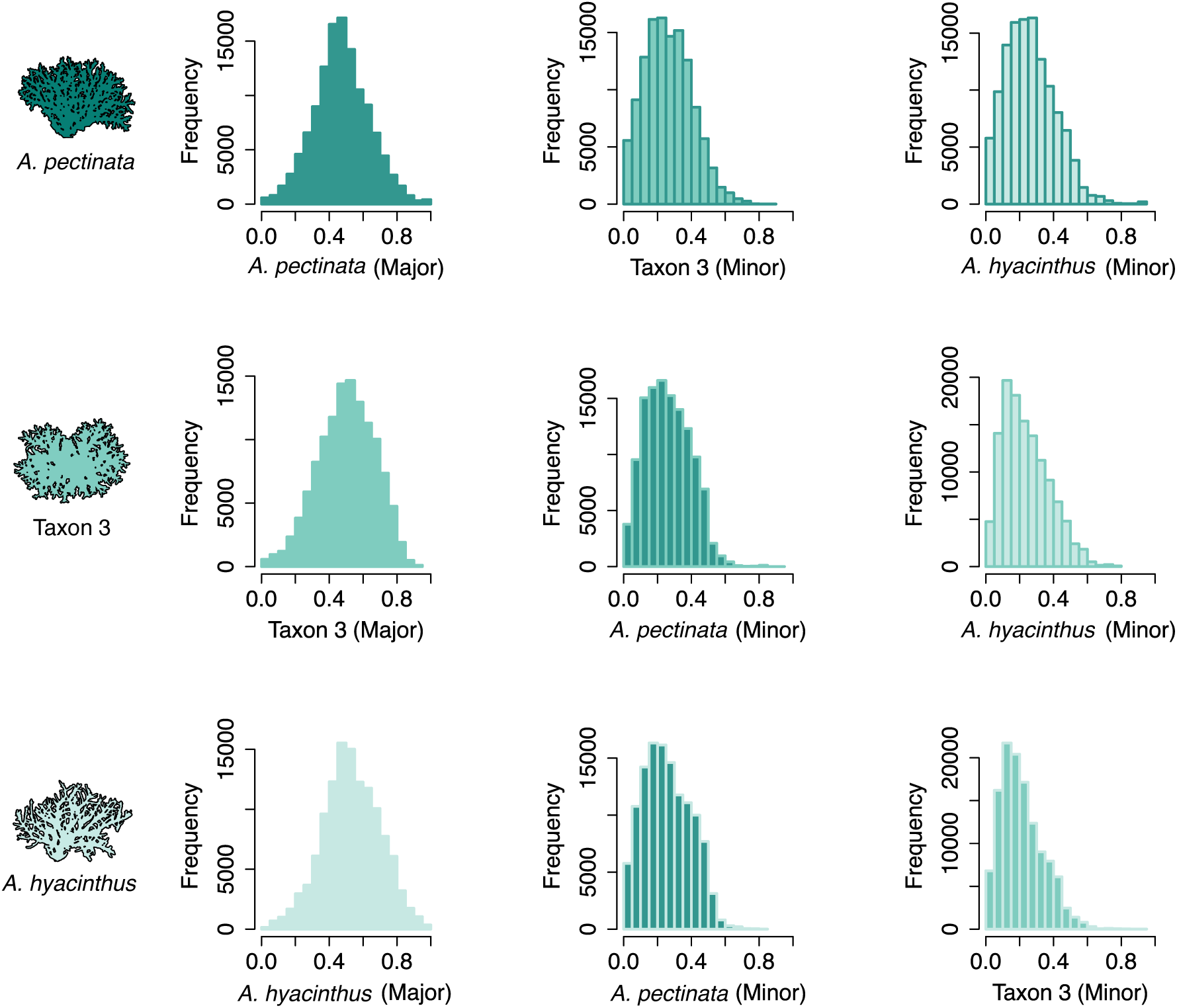
Summary of genome-wide local ancestry frequency distributions for three admixed AH species based on local ancestry inference with Ancestry_HMM, where the histogram border colour indicates the major-parent genomic background. Major-parent ancestry distributions (left column) were centred around 50%, with many sites showing balanced ancestry proportions for both major and minor-parent ancestries. In contrast, minor-parent distributions (middle and right columns) were centred around 20% with few regions showing minor-parent ancestries exceeding 50%.

**Figure S25.**
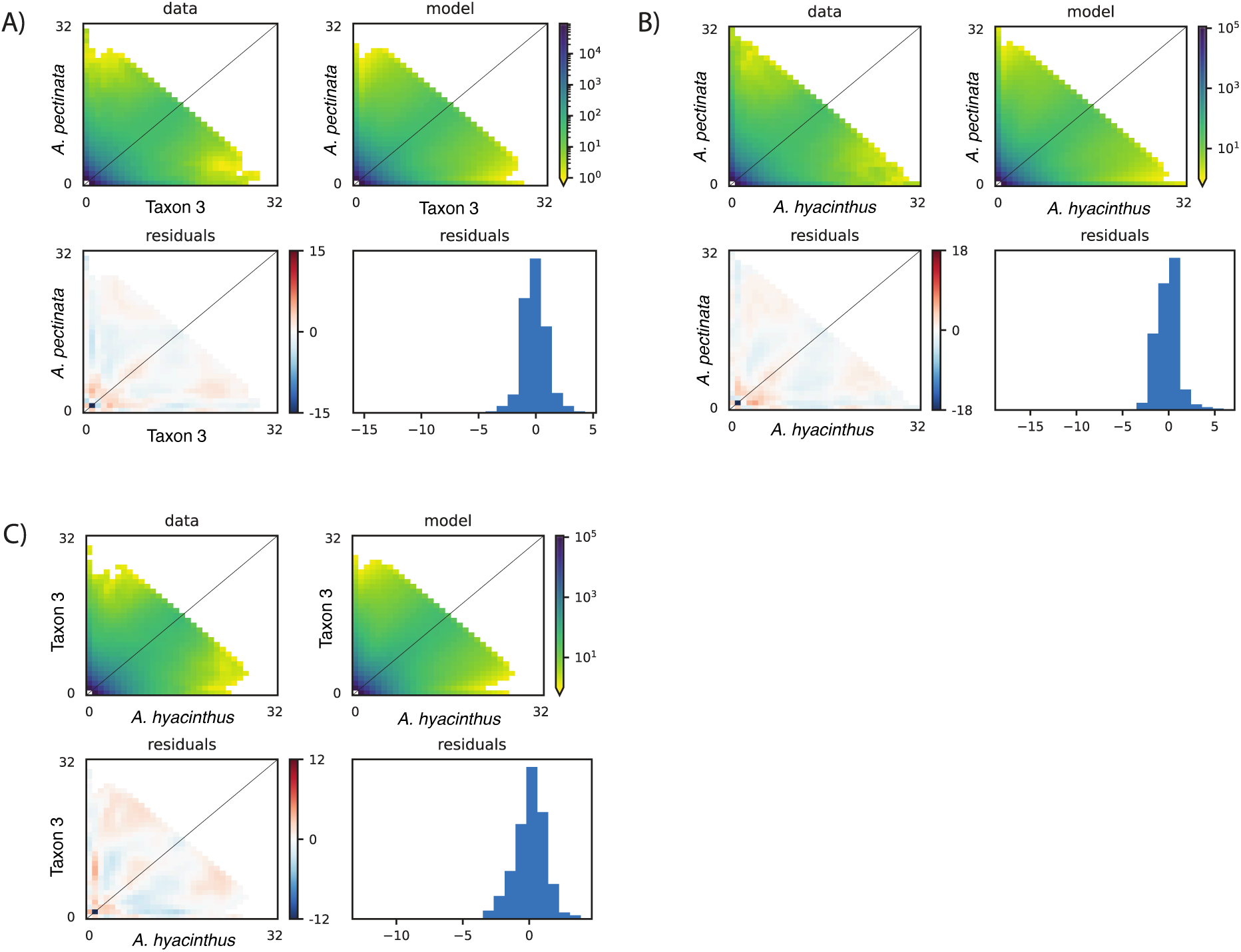
Residuals (model - data) for favoured heterogeneous gene flow *dadi* models among *A. pectinata*, Taxon 3 and *A. hyacinthus* comparisons. Most bias occurs at low frequency bins however differences amongst the model and data are negligible (12 - 18 SNPs) and residuals are normally distributed.

**Figure S26.**
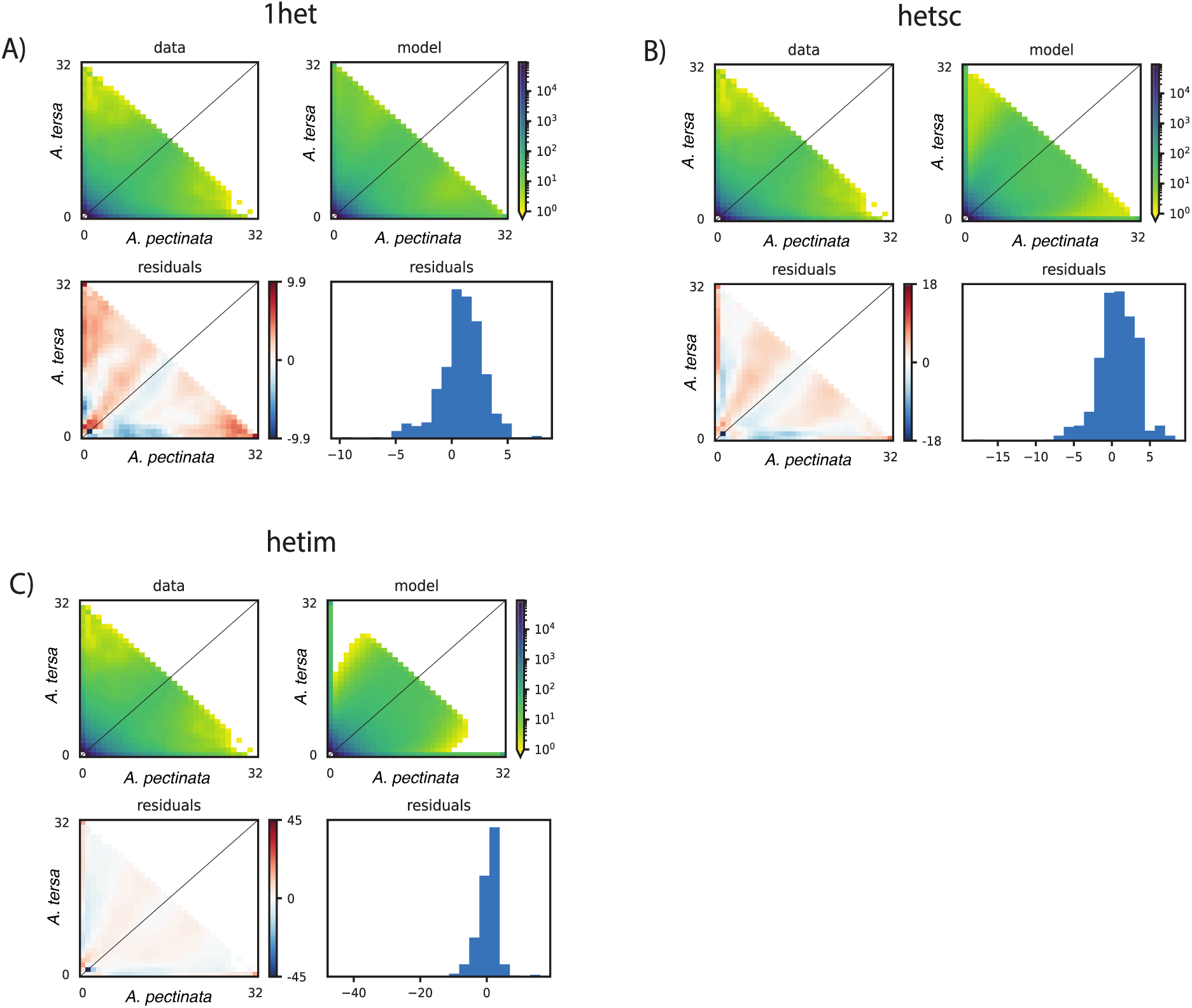
Residuals (model - data) for three different heterogeneous gene flow *dadi* models (1het, hetsc and hetim) for *A. tersa* and *A. pectinata* comparisons. 1het is a continuous gene flow model with two different gene flow periods with different gene flow rates; hetsc is a secondary contact model and hetim is a continuous gene flow model with only one gene flow rate. The secondary contact model (hetsc) is the favoured model following log-likelihood ratio tests with the more complex model. Bias occurs at shared frequency bins and private allele frequencies; however, differences among models and data are negligible (10-45 SNPs) and residuals are slightly positively skewed.

**Figure S27.**
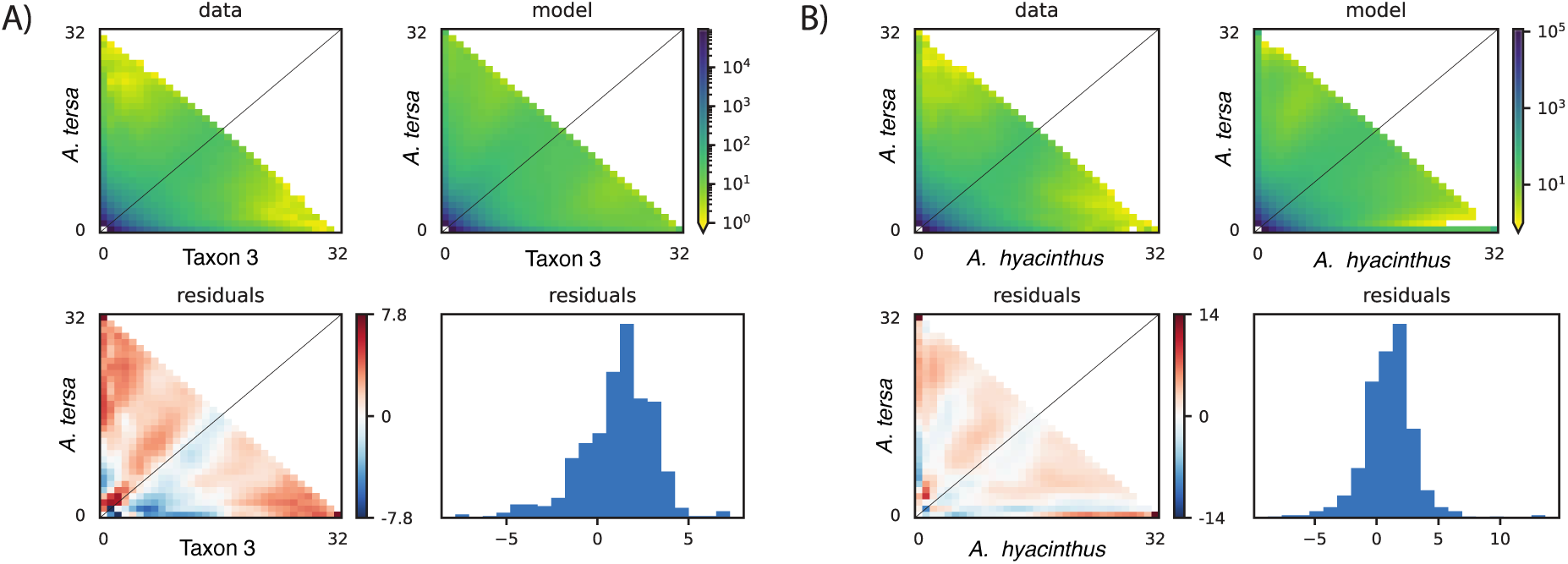
Residuals (model - data) for favoured heterogeneous gene flow *dadi* models for *A. tersa* comparisons with Taxon 3 and *A. hyacinthus.* Some bias occurs; however, differences among models and data are negligible (8-14 SNPs) and residuals are slightly positively skewed.

**Figure S28.**
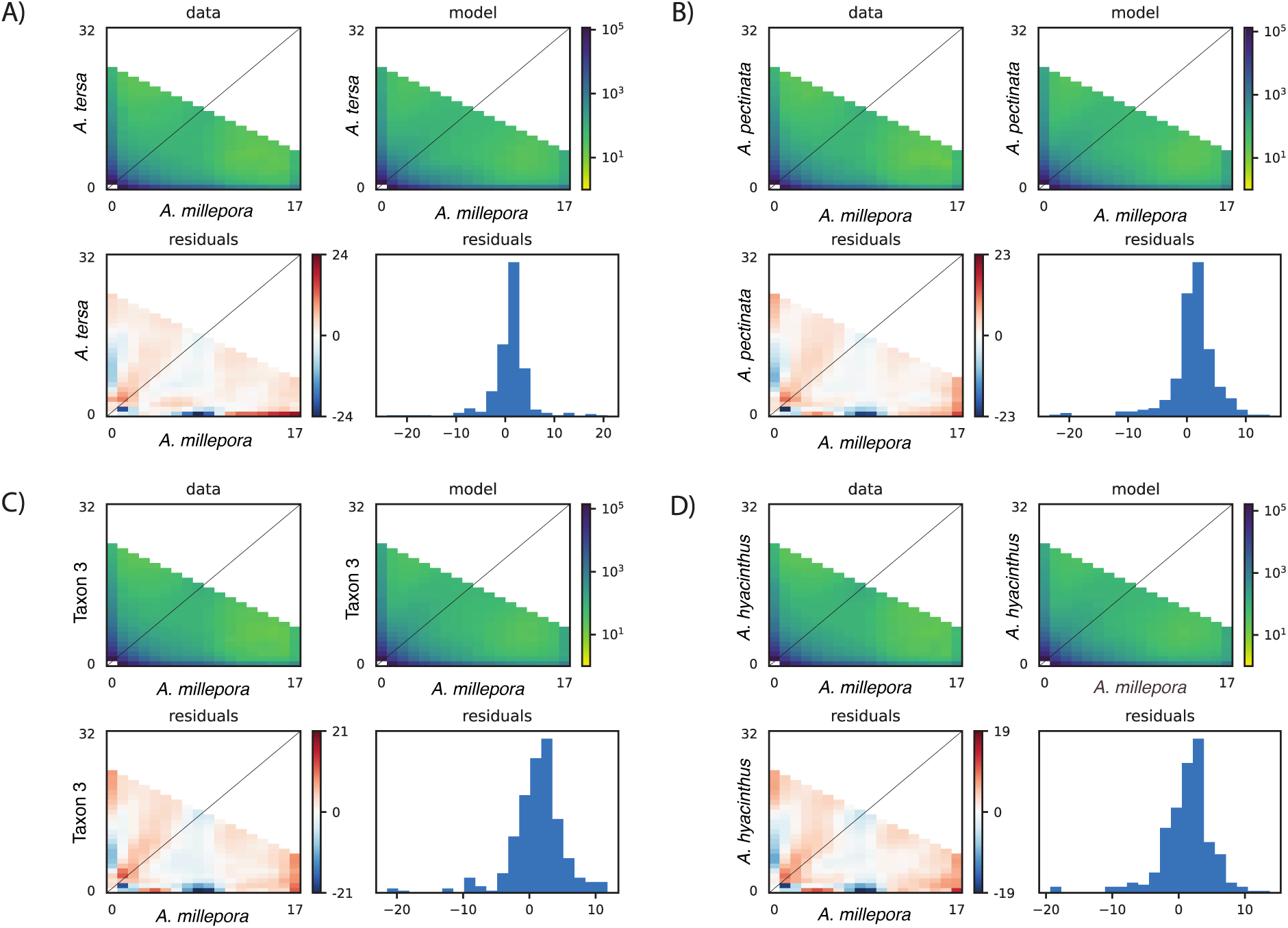
Residuals (model - data) for favoured heterogeneous gene flow *dadi* models for *A. millepora* and four AH species. Although some bias occurs, differences among the models and data are negligible (19-34 SNPs) and residuals are slightly positively skewed.

**Table S1.** D-statistics for genome-wide introgression among AH species in northern, central and southern GBR regions. D-statistics were implemented in ANGSD using the *-doAbbabab2* option with *-blocksize* of 5 Mbp for a four-population test and using *A. kenti* as an outgroup. Abbreviations are as follows: nABBA = number of ABBA patterns, nBABA = number of BABA patterns, nBlocks = number of blocks with data. Significant comparisons are shown in bold and grey shaded rows. Numbers correspond to species as follows: 1 = *A. tersa*, 2 = *A. pectinata*, 3 = Taxon 3 and 4 = *A. hyacinthus*.

| Region | D-statistic | Taxon P1 | Taxon P2 | Taxon P3 | Taxon P4 | nBlocks | nABBA | nBABA | P-value | Z Score | Gene flow Test |
| --- | --- | --- | --- | --- | --- | --- | --- | --- | --- | --- | --- |
| North | -0.0067 | 4 | 2 | 1 | *A. kenti* | 576 | 1796 | 1820 | 0.056 | -1.91 | NA |
| North | 0.0027 | 3 | 2 | 1 | *A. kenti* | 577 | 1715 | 1706 | 0.302 | 1.03 | NA |
| North | -0.0099 | 4 | 3 | 1 | *A. kenti* | 577 | 1671 | 1705 | 0.00074 | -3.37 | NA |
| **North** | **0.104** | **4** | **3** | **2** | ***A. kenti*** | **569** | **1966** | **1597** | **0** | **33.77** | **2 : 3** |
| Central | -0.009 | 4 | 2 | 1 | *A. kenti* | 562 | 1792 | 1825 | 0.0048 | -2.82 | NA |
| Central | -0.0009 | 3 | 2 | 1 | *A. kenti* | 562 | 1710 | 1713 | 0.74 | -0.34 | NA |
| Central | -0.0087 | 4 | 3 | 1 | *A. kenti* | 555 | 1679 | 1708 | 0.0017 | -3.14 | NA |
| Central | 0.105 | 4 | 3 | 2 | *A. kenti* | 552 | 1962 | 1590 | 0 | 34.82 | 2 : 3 |
| South | -0.0055 | 4-2 | 2-2 | 1 | *A. kenti* | 568 | 1804 | 1824 | 0.10 | -1.62 | NA |
| South | -0.0103 | 4-1 | 2-2 | 1 | *A. kenti* | 572 | 1801 | 1839 | 0.0017 | -3.13 | NA |
| South | 0.0068 | 3-2 | 2-2 | 1 | *A. kenti* | 570 | 1723 | 1700 | 0.0070 | 2.70 | NA |
| **South** | **-0.013** | **4-2** | **3-2** | **1** | ***A. kenti*** | **568** | **1691** | **1734** | **0.000003** | **-4.66** | **4-2 : 1** |
| **South** | **-0.018** | **4-1** | **3-2** | **1** | ***A. kenti*** | **574** | **1691** | **1752** | **0** | **-6.36** | **4-1 : 1** |
| South | 0.0054 | 4-2 | 4-1 | 1 | *A. kenti* | 573 | 1634 | 1617 | 0.0054 | 2.78 | NA |
| **South** | **0.105** | **4-2** | **3-2** | **2-2** | ***A. kenti*** | **562** | **2046** | **1659** | **0** | **38.02** | **3-2 : 2-2** |
| **South** | **0.12** | **4-1** | **3-2** | **2-2** | ***A. kenti*** | **568** | **2068** | **1619** | **0** | **41.65** | **3-2 : 2-2** |
| **South** | **-0.019** | **4-2** | **4-1** | **2-2** | ***A. kenti*** | **565** | **1606** | **1668** | **0** | **-8.92** | **4-2 : 2-2** |
| **South** | **-0.021** | **4-2** | **4-1** | **3-2** | ***A. kenti*** | **568** | **1750** | **1825** | **0** | **-10.40** | **4-2 : 3-2** |

**Table S2.** D-statistics for genome-wide introgression among *A. millepora* and AH species in northern, central and southern GBR regions. D-statistics were implemented in ANGSD using the *-doAbbabab2* option with *-blocksize* of 5 Mbp for a four-population test and using *A. kenti* as an outgroup. Abbreviations are as follows: nABBA = number of ABBA patterns, nBABA = number of BABA patterns, nBlocks = number of blocks with data. Significant comparisons are shown in bolded and grey shaded rows. Numbers correspond to species as follows: 1=*A. tersa*, 2=*A. pectinata*, 3=Taxon 3 and 4=*A. hyacinthus*.

| Region | D-statistic | Taxon P1 | Taxon P2 | Taxon P3 | Taxon P4 | nBlocks | nABBA | nBABA | P-value | Z Score | Gene Flow Test |
| --- | --- | --- | --- | --- | --- | --- | --- | --- | --- | --- | --- |
| North | **-0.03** | **4** | **1** | *A. millepora* | *A. kenti* | **563** | **1212** | **1287** | **0** | **-7.74** | *A. millepora* : 4 |
| North | **-0.044** | **3** | **1** | *A. millepora* | *A. kenti* | **562** | **1193** | **1303** | **0** | **-11.69** | *A. millepora* : 3 |
| North | **-0.037** | **2** | **1** | *A. millepora* | *A. kenti* | **562** | **1218** | **1312** | **0** | **-9.62** | *A. millepora* : 2 |
| North | 0.0070 | 4 | 2 | *A. millepora* | *A. kenti* | 560 | 1305 | 1286 | 0.038 | 2.072 | NA |
| North | -0.0063 | 3 | 2 | *A. millepora* | *A. kenti* | 547 | 1227 | 1242 | 0.037 | -2.087 | NA |
| North | **0.014** | **4** | **3** | *A. millepora* | *A. kenti* | **558** | **1246** | **1213** | **0.000014** | **4.34** | *A. millepora* : 3 |
| Central | **-0.02886** | **4** | **1** | *A. millepora* | *A. kenti* | **562** | **1211** | **1283** | **0** | **-7.26** | *A. millepora* : 4 |
| Central | **-0.04576** | **3** | **1** | *A. millepora* | *A. kenti* | **561** | **1189** | **1303** | **0** | **-12.06** | *A. millepora* : 3 |
| Central | **-0.03669** | **2** | **1** | *A. millepora* | *A. kenti* | **564** | **1215** | **1308** | **0** | **-9.87** | *A. millepora* : 2 |
| Central | 0.007971 | 4 | 2 | *A. millepora* | *A. kenti* | 554 | 1304 | 1284 | 0.025 | 2.25 | NA |
| Central | -0.00858 | 3 | 2 | *A. millepora* | *A. kenti* | 555 | 1223 | 1244 | 0.012 | -2.50 | NA |
| Central | **0.017095** | **4** | **3** | *A. millepora* | *A. kenti* | **540** | **1248** | **1206** | **0.000003** | **4.64** | *A. millepora* : 3 |
| South | -0.01317 | 4-2 | 1 | *A. millepora* | *A. kenti* | 564 | 1233 | 1266 | 0.00011 | -3.87 | NA |
| South | **-0.02471** | **4-1** | **1** | *A. millepora* | *A. kenti* | **564** | **1217** | **1279** | **0** | **-7.023** | *A. millepora* : 4-1 |
| South | **-0.03106** | **3-2** | **1** | *A. millepora* | *A. kenti* | **567** | **1221** | **1300** | **0** | **-8.19** | *A. millepora* : 3-2 |
| South | **-0.01855** | **2-2** | **1** | *A. millepora* | *A. kenti* | **564** | **1238** | **1285** | **0** | **-5.18** | *A. millepora* : 2-2 |
| South | 0.00529 | 4-2 | 2-2 | *A. millepora* | *A. kenti* | 555 | 1296 | 1283 | 0.11 | 1.60 | NA |
| South | -0.00601 | 4-1 | 2-2 | *A. millepora* | *A. kenti* | 561 | 1290 | 1305 | 0.053 | -1.94 | NA |
| South | **-0.01275** | **3-2** | **2-2** | *A. millepora* | *A. kenti* | **559** | **1220** | **1252** | **0.000007** | **-4.50** | *A. millepora* : 3-2 |
| South | **0.018135** | **4-2** | **3-2** | *A. millepora* | *A. kenti* | **557** | **1263** | **1218** | **0** | **6.31** | *A. millepora* : 3-2 |
| South | 0.006449 | 4-1 | 3-2 | *A. millepora* | *A. kenti* | 564 | 1256 | 1240 | 0.029 | 2.18 | NA |
| South | **0.012323** | **4-2** | **4-1** | *A. millepora* | *A. kenti* | **560** | **1190** | **1161** | **0** | **6.18** | *A. millepora* : 4-1 |

**Table S3-A.** Optimised model parameters for divergence times and effective population sizes from demographic modelling in *dadi*. Physical units with confidence intervals are in brackets and numbers correspond to species as follows: 1 = *A. tersa*, 2 = *A. pectinata*, 3 = Taxon 3, 4 = *A. hyacinthus* and *Amil* = *A. millepora.* Divergence time periods (*T_E1_* and *T_E2_*) and population sizes *(N1, N2, N11, N12, N21, N22*) are in units of 1000 generations and 1000 individuals respectively, where the divergence time is equivalent to the sum of *T_E1_* and *T_E2_.* Lower parameter bounds for **bolded** models hit 0 values. In these cases, nested models (excluding these parameters; shown in grey) were optimised and log-likelihood ratio tests were conducted among them to assess whether to include the parameter in the final model (refer to Table S6).

| Taxon | Model | LogL | $T_{E1}$ | $N1$ | $N2$ | $N11$ | $N12$ | $T_{E2}$ | $N21$ | $N22$ | $\theta$ |
| --- | --- | --- | --- | --- | --- | --- | --- | --- | --- | --- | --- |
| 1-2 | *lhet* | -2754 | 87.57<br>(24.17,<br>151) | **27.29 (0,**<br>**76.2)** | 259 (243.7,<br>274.3) | 541.3<br>(530.1,<br>552.5) | 530.4 (511,<br>549.7) | **0.41 (0,**<br>**8.728)** | 553.2<br>(113.7,<br>992.8) | 475.4<br>(415.2,<br>535.6) | 3994 |
| 1-2 | *hetsc* | -3445 | 60.47<br>(59.09,<br>61.86) | 32.72<br>(21.71,<br>43.73) | 7391<br>(6846,<br>7936) | 554.6<br>(528.4,<br>580.9) | 360.4<br>(358.7,<br>362.1) | 20.35<br>(5.011,<br>35.69) | 551.2<br>(549.3,<br>553.1) | 592.3 (545,<br>639.7) | 6089 |
| 1-2 | *hetim* | -4910 | 83.37<br>(79.39,<br>87.36) | 24.06<br>(8.094,<br>40.02) | 316 (301.3,<br>330.8) | 546.8<br>(535.9,<br>557.6) | 521.8<br>(509.2,<br>534.3) | - | - | - | 5160 |
| 1-3 | *lhet* | -2793 | 325.9<br>(316.2,<br>335.6) | 136.3<br>(129.5,<br>143.1) | 2190<br>(1747,<br>2634) | 2477<br>(2398,<br>2555) | 2010<br>(1811,<br>2210) | 16.66<br>(5.544,<br>27.77) | 1775<br>(1734,<br>1816) | 3149<br>(2002,<br>4296) | 5448 |
| 1-4 | *lhet* | -2864 | 368.1<br>(358.9,<br>377.3) | 103.5<br>(78.47,<br>128.5) | 25500<br>(25480,<br>25520) | 1845<br>(1819,<br>1872) | 2550<br>(2540,<br>2561) | **7.53 (0,**<br>**34.6)** | 3178<br>(3094,<br>3263) | 4378<br>(3981,<br>4776) | 8177 |
| 2-3 | *lhet* | -1759 | 291.1<br>(270.7,<br>311.4) | 736.4<br>(709.2,<br>763.6) | 718.9<br>(704.2,<br>733.6) | 2279<br>(2265,<br>2294) | 2710<br>(2701,<br>2720) | **2.03 (0,**<br>**5.379)** | 460.2<br>(369.4,<br>550.9) | 255.1<br>(199.7,<br>310.4) | 5047 |
| 2-4 | *lhet* | -1940 | 322.4<br>(315.9,<br>328.8) | 485.2<br>(484.1,<br>486.3) | 906.9<br>(900.7,<br>913.1) | 2387<br>(2381,<br>2392) | 3729<br>(3723,<br>3736) | **3.47 (0,**<br>**7.776)** | 387.2<br>(375.8,<br>398.6) | 459.3<br>(449.8,<br>468.7) | 4770 |
| 3-4 | *lhet* | -1757 | 282.1<br>(268.8,<br>295.4) | 510.6<br>(503.6,<br>517.7) | 1303<br>(1262,<br>1344) | 1683<br>(1663,<br>1703) | 3158<br>(3112,<br>3204) | **3.91 (0,**<br>**8.105)** | 358.9<br>(340.6,<br>377.1) | 629.5<br>(534.6,<br>724.5) | 6535 |
| 1-*Amil* | *Ihet* | -3754 | 317.3<br>(279.1,<br>355.6) | 79.61<br>(57.6,<br>101.6) | 49190<br>(48980,<br>49400) | 1497<br>(1468,<br>1526) | 1033<br>(1025,<br>1041) | 45.95<br>(31.09,<br>60.8) | 3011<br>(2947,<br>3075) | 1039<br>(1024,<br>1054) | 24260 |
| 2-*Amil* | *Ihet* | -3665 | 378.9<br>(334.8,<br>423) | 118.6<br>(86.84,<br>150.4) | 28970<br>(28890,<br>29050) | 1928<br>(1923,<br>1932) | 799.8<br>(796.3,<br>803.3) | **45.02 (0,**<br>**153.3)** | 2418<br>(2404,<br>2432) | 1400<br>(1256,<br>1544) | 14560 |
| 3-*Amil* | *Ihet* | -3797 | 396.7<br>(387.7,<br>405.7) | 132.1 (0,<br>312.8) | 26410<br>(26250,<br>26570) | 1958<br>(1803,<br>2112) | 790.9<br>(779.3,<br>802.5) | 24.77<br>(23.14,<br>26.41) | 2308<br>(2044,<br>2573) | 1629<br>(1314,<br>1945) | 14160 |
| 4-*Amil* | *Ihet* | -3547 | 417.7 (242,<br>593.4) | 122.1<br>(106.1,<br>138.2) | 27290<br>(27140,<br>27450) | 2251<br>(2135,<br>2366) | 743.3<br>(657.3,<br>829.4) | 26.64<br>(15.12,<br>38.16) | 2407<br>(2260,<br>2553) | 1059<br>(1020,<br>1098) | 15700 |

**Table S3-B.** Optimised model parameters for migration from demographic modelling in *dadi*. Physical units with confidence intervals are in brackets and numbers correspond to species as follows: 1=*A. tersa*, 2=*A. pectinata*, 3=Taxon 3, 4=*A. hyacinthus* and *Amil* = *A. millepora.* Gene flow parameters *Nm1_12, Nm1_21, Nm2_12,* and *Nm2_21* represent the number of migrants entering the population per generation at *1* – *P(E1,E2)* proportion of the genome, where the parameters *P_E1_* and *P_E2_* represent the proportion of the genome experiencing no gene flow in the first and second time periods respectively (i.e. Epoch 1 and Epoch 2). Migration rate parameters *m1_12, m1_21, m2_12,* and *m2_21* represent the proportion of migrants entering the population per generation. Lower parameter bounds for **bolded** models hit 0 values. In these cases, nested models (excluding these parameters; shown in grey) were optimised and log-likelihood ratio tests were conducted among them to assess whether to include the parameter in the final model (refer to Table S6).

| Taxon | Model | $Nm1\_12$ | $m1\_12$ | $Nm1\_21$ | $m1\_21$ | $Nm2\_12$ | $P_{E1}$ | $m2\_12$ | $Nm2\_21$ | $m2\_21$ | $P_{E2}$ |
| --- | --- | --- | --- | --- | --- | --- | --- | --- | --- | --- | --- |
| 1-2 | *lhet* | 0.22 (0, 15.3) | 4.06429E-07 | **7.313 (0, 36.97)** | 1.37877E-06 | 256.7 (242.8, 270.7) | **0.36 (0, 0.74)** | 0.000464027 | 209 (189.9, 228.2) | 0.00043963 | 0.44 (0.12, 0.75) |
| 1-2 | *hetsc* | - | - | - | - | 6.377 (4.929, 7.825) | - | 1.15693E-05 | 19.43 (16.75, 22.12) | 3.28043E-05 | 0.29 (0.12, 0.46) |
| 1-2 | *hetim* | **1.818 (0, 14.96)** | 3.3248E-06 | 6.671 (6.559, 6.783) | 1.27846E-05 | - | **0.16 (0, 0.32)** | - | - | - | - |
| 1-3 | *lhet* | 0.78 (0, 1.941) | 3.14897E-07 | **0.95 (0, 2.442)** | 4.72637E-07 | 23.11 (14.99, 31.23) | **0.18 (0, 0.41)** | 1.30197E-05 | 173.8 (162.2, 185.4) | 5.51921E-05 | 0.46 (0.44, 0.49) |
| 1-4 | *lhet* | 0.15 (0, 1.038) | 8.13008E-08 | 8.47 (4.93, 12) | 3.32157E-06 | 124.6 (113.2, 136.1) | 0.29 (0.054, 0.52) | 3.9207E-05 | 50.26 (40.94, 59.57) | 1.14801E-05 | 0.42 (0.36, 0.48) |
| 2-3 | *lhet* | 124.9 (117.1, 132.8) | 5.48047E-05 | 91.2 (57.03, 125.4) | 1.2543E-06 | 40.44 (39.08, 41.81) | 0.65 (0.64, 0.66) | 8.78748E-05 | 25.68 (24.87, 26.49) | 0.000100666 | 0.29 (0.24, 0.34) |
| 2-4 | *lhet* | 115.6 (70.07, 161.2) | 4.8429E-05 | 130.3 (76.97, 183.7) | 3.45358E-06 | 22.52 (21.79, 23.24) | 0.66 (0.63, 0.68) | 5.81612E-05 | 19.02 (18.57, 19.48) | 4.14108E-05 | 0.40 (0.28, 0.52) |
| 3-4 | *Ihet* | 114.4<br>(95.09,<br>133.7) | 6.79739E-05 | 104.4<br>(36.07,<br>172.7) | 3.30589E-05 | 24.57<br>(23.97,<br>25.17) | 0.54 (0.52,<br>0.56) | 6.36693E-05 | 15.86<br>(15.04,<br>16.68) | 2.51946E-05 | 0.16 (0,<br>0.46) |
| 1-*Amil* | *Ihet* | 0.053<br>(0.0055,<br>0.1) | 3.54041E-08 | **0.25 (0,**<br>**0.67)** | 2.42014E-07 | 6.768<br>(2.748,<br>10.79) | **0.15 (0,**<br>**0.31)** | 2.24776E-06 | 21.12<br>(19.65,<br>22.58) | 2.03272E-05 | 0.58 (0.57,<br>0.58) |
| 2-*Amil* | *Ihet* | **0.14 (0,**<br>**2.679)** | 7.26141E-08 | **0.13 (0,**<br>**2.944)** | 1.62541E-07 | 7.651<br>(3.898,<br>11.4) | **0.16 (0,**<br>**0.33)** | 3.16419E-06 | 30.01<br>(24.41,<br>35.62) | 2.14357E-05 | 0.57 (0.49,<br>0.66) |
| 3-*Amil* | *Ihet* | **0.17 (0,**<br>**2.81)** | 8.68233E-08 | **0.19 (0,**<br>**0.82)** | 2.40233E-07 | 13.95<br>(9.673,<br>18.22) | 0.24 (0.14,<br>0.33) | 6.04419E-06 | 61.69<br>(41.12,<br>82.26) | 3.78699E-05 | 0.6 (0.59,<br>0.6) |
| 4-*Amil* | *Ihet* | 0.22<br>(0.012,<br>0.42) | 9.77343E-08 | 0.26 (0.21,<br>0.3) | 3.49791E-07 | **12.68 (0,**<br>**55.9)** | 0.26 (0.14,<br>0.38) | 5.26797E-06 | 38.8<br>(27.21,<br>50.4) | 3.66383E-05 | 0.58 (0.57,<br>0.59) |

**Table S4.** Two-sample Kolmogorov-Smirnov tests linkage (r^2^) distributions for three datasets in each taxon. *Minor-parent peaks:* comparisons among sites with private or shared minor-parent ancestry peaks (exceeding 50% ancestry proportion); *Candidate regions:* comparisons among target sites falling within 29 candidate genomic regions for parallel admixture (normalised to 200 kb base pairs wide); and *Random:* comparisons among a randomly selected subset of ancestry informative sites equivalent to the number of sites in *Minor peaks*.

| Genomic Background | Dataset A | Dataset B | D | P-value | P-value adjusted (BH) |
| --- | --- | --- | --- | --- | --- |
| *Acropora pectinata* | *Random* | *Minor-parent peaks* | 0.04625 | $2.53 \times 10^{-93}$ | $7.58 \times 10^{-93}$ |
| *Acropora pectinata* | *Random* | *Candidate regions* | 0.04615 | $6.37 \times 10^{-93}$ | $9.55 \times 10^{-93}$ |
| *Acropora pectinata* | *Minor peaks* | *Candidate regions* | 0.02593 | $1.26 \times 10^{-29}$ | $1.26 \times 10^{-29}$ |
| Taxon 3 | *Random* | *Minor-parent peaks* | 0.01985 | $1.54 \times 10^{-17}$ | $1.54 \times 10^{-17}$ |
| Taxon 3 | *Random* | *Candidate regions* | 0.05517 | $1.30 \times 10^{-132}$ | $3.90 \times 10^{-132}$ |
| Taxon 3 | *Minor peaks* | *Candidate regions* | 0.03687 | $1.83 \times 10^{-59}$ | $2.75 \times 10^{-59}$ |
| *Acropora hyacinthus* | *Random* | *Minor-parent peaks* | 0.03709 | $3.60 \times 10^{-60}$ | $5.40 \times 10^{-60}$ |
| *Acropora hyacinthus* | *Random* | *Candidate regions* | 0.03815 | $1.24 \times 10^{-63}$ | $3.71 \times 10^{-63}$ |
| *Acropora hyacinthus* | *Minor peaks* | *Candidate regions* | 0.01293 | $1.10 \times 10^{-07}$ | $1.10 \times 10^{-07}$ |

**Table S5.**
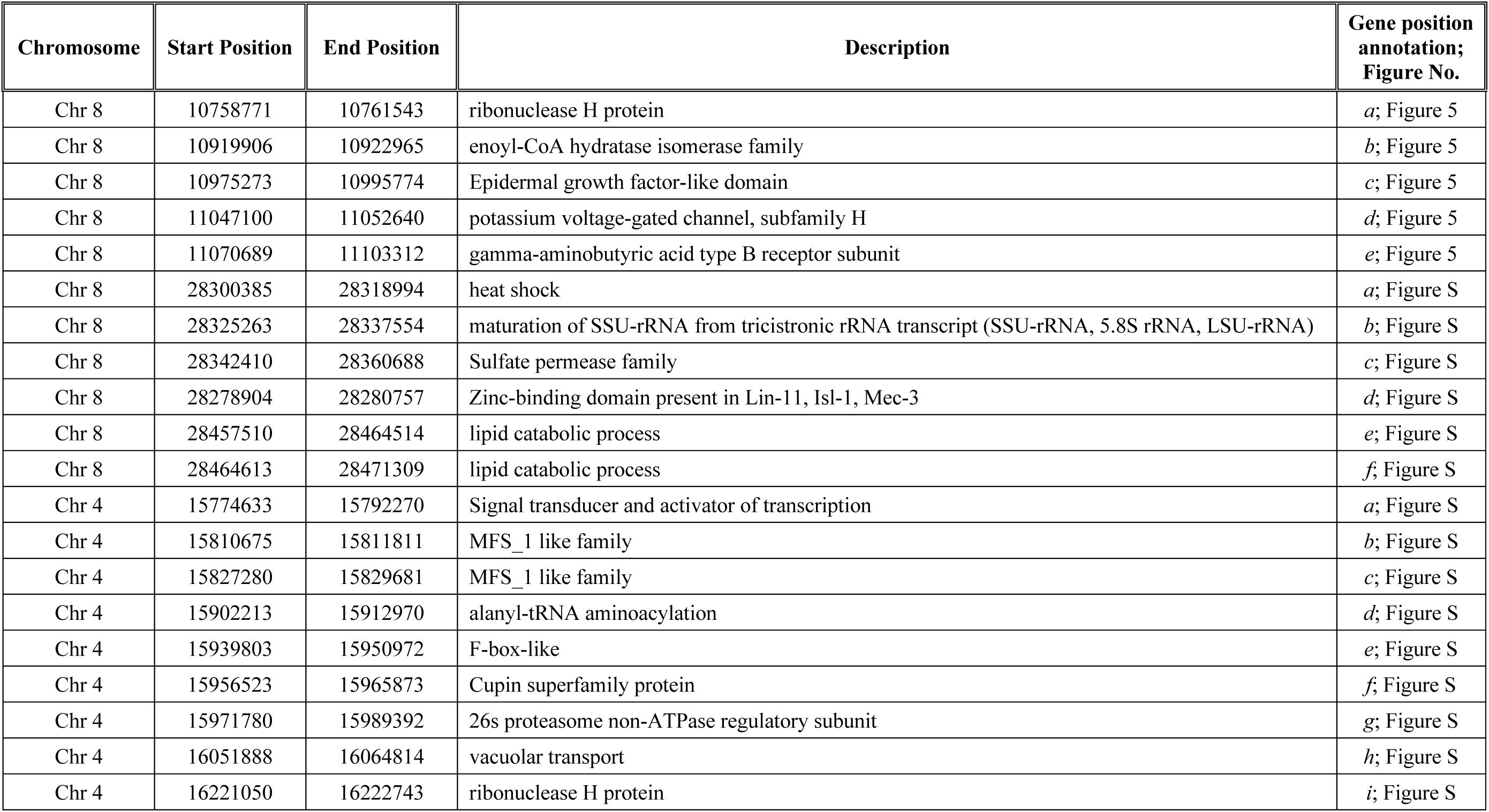
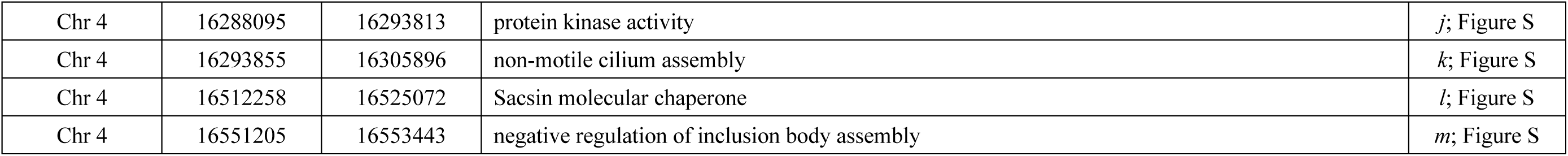
Functional annotations for genes surrounding example candidate regions for parallel adaptive admixture. Focal candidate regions correspond to Chromosome 8 (10.85-10.92 Mb; Figure 5), Chromosome 8 (28.32-28.32 Mb; Figure S21) and Chromosome 4 (16.17-16.2 Mb; Figure S22). Gene positions are annotated by *italic* letters in corresponding figures.

| Chromosome | Start Position | End Position | Description | Gene position annotation; Figure No. |
| --- | --- | --- | --- | --- |
| Chr 8 | 10758771 | 10761543 | ribonuclease H protein | *a*; Figure 5 |
| Chr 8 | 10919906 | 10922965 | enoyl-CoA hydratase isomerase family | *b*; Figure 5 |
| Chr 8 | 10975273 | 10995774 | Epidermal growth factor-like domain | *c*; Figure 5 |
| Chr 8 | 11047100 | 11052640 | potassium voltage-gated channel, subfamily H | *d*; Figure 5 |
| Chr 8 | 11070689 | 11103312 | gamma-aminobutyric acid type B receptor subunit | *e*; Figure 5 |
| Chr 8 | 28300385 | 28318994 | heat shock | *a*; Figure S |
| Chr 8 | 28325263 | 28337554 | maturation of SSU-rRNA from tricistronic rRNA transcript (SSU-rRNA, 5.8S rRNA, LSU-rRNA) | *b*; Figure S |
| Chr 8 | 28342410 | 28360688 | Sulfate permease family | *c*; Figure S |
| Chr 8 | 28278904 | 28280757 | Zinc-binding domain present in Lin-11, Isl-1, Mec-3 | *d*; Figure S |
| Chr 8 | 28457510 | 28464514 | lipid catabolic process | *e*; Figure S |
| Chr 8 | 28464613 | 28471309 | lipid catabolic process | *f*; Figure S |
| Chr 4 | 15774633 | 15792270 | Signal transducer and activator of transcription | *a*; Figure S |
| Chr 4 | 15810675 | 15811811 | MFS_1 like family | *b*; Figure S |
| Chr 4 | 15827280 | 15829681 | MFS_1 like family | *c*; Figure S |
| Chr 4 | 15902213 | 15912970 | alanyl-tRNA aminoacylation | *d*; Figure S |
| Chr 4 | 15939803 | 15950972 | F-box-like | *e*; Figure S |
| Chr 4 | 15956523 | 15965873 | Cupin superfamily protein | *f*; Figure S |
| Chr 4 | 15971780 | 15989392 | 26s proteasome non-ATPase regulatory subunit | *g*; Figure S |
| Chr 4 | 16051888 | 16064814 | vacuolar transport | *h*; Figure S |
| Chr 4 | 16221050 | 16222743 | ribonuclease H protein | *i*; Figure S |
| Chr 4 | 16288095 | 16293813 | protein kinase activity | *j*; Figure S |
| Chr 4 | 16293855 | 16305896 | non-motile cilium assembly | *k*; Figure S |
| Chr 4 | 16512258 | 16525072 | Sacsin molecular chaperone | *l*; Figure S |
| Chr 4 | 16551205 | 16553443 | negative regulation of inclusion body assembly | *m*; Figure S |

**Table S6.** Optimal single-population models including population size changes. Best log-likelihood and AIC models for AH species and *A. millepora* using the one-dimensional site frequency spectrum demographic modelling in *dadi.* Models tested: snm (standard neutral model), size_change, bottleneck, bottle.

| Population | Best Model | LL | AIC | Parameters | Parameter Values | Direction of Size Change |
| --- | --- | --- | --- | --- | --- | --- |
| *Amil* | bottle | -754.12 | 1514.24 | $nuB, nuF, T$ | 7.54,3.58,0.42 | Increase - Decrease |
| 1 | size_change | -613.24 | 1230 | $nu, T$ | 22.75,2.08 | Increase |
| 2 | bottle_neck | -126.62 | 261.14 | $nuB, nuF, TB, TF$ | 0.01, 5.82, 0.01, 0.67 | Decrease - Increase |
| 3 | size_change | -174.85 | 353.7 | $nu, T$ | 32.59, 3.53 | Increase |
| 4 | size_change | -421.1 | 846.2 | $nu, T$ | 30.97,3.51 | Increase |

**Snm:** Standard neutral model. No changes occur.
**Size\_change:** Instantaneous size change. Parameters: $nu$ and $T$ . $nu$ : ratio of contemporary to ancient population size. $T$ : time at which instantaneous size change occurred.
**Bottleneck:** Two instantaneous change periods. Parameters: $nuB, nuF, TB$ and $TF$ . $nuB$ : ratio of contemporary to ancient population size at first instantaneous size change, $nuF$ : ratio of contemporary to ancient population size at second instantaneous size change. $TB$ : time first change occurred. $TF$ : time a second change occurred.
**Bottle:** Instantaneous size change followed by exponential growth/shrink. Parameters: $nuB, nuF$ and $T$ . $nuB$ : ratio of population size after instantaneous change to ancient population size. $NuF$ : ratio of contemporary to ancient population size. $T$ : time in the past at which the instantaneous change happened and the exponential growth began.

**Table S7.** Log-likelihood ratio tests among full and nested models using the bootstrap Godambe adjustment. Numbers correspond to species as follows: 1 = *A. tersa*, 2 = *A. pectinata*, 3 = Taxon 3, 4 = *A. hyacinthus,* and *Amil* = *A. millepora*

| Taxon Comparison | Complex model | Simple model | Log-likelihood complex model | Log-likelihood simple model | Godambe adjustment | Adjusted LRT statistic | LRT P-value | Accepted model |
| --- | --- | --- | --- | --- | --- | --- | --- | --- |
| 1-2 | *lhet*<sup>a</sup> | *lhetim*<sup>b</sup> | -2754 | -4910 | 0.0007 | 2.9 | 0.08 | *lhetim* |
| 1-2 | *lhet* | *lhetsc*<sup>c</sup> | -2754 | -3445 | 0.0006 | 0.9 | 0.35 | *lhetsc* |
| 1-3 | *lhet* | *lhetsc* | -2793 | -3404 | 0.87 | 1066 | 0.0* | *lhet* |
| 1-4 | *lhet* | *lhetsc* | -2864 | -4064 | 0.12 | 315 | 0.0* | *lhet* |
| 1-4 | *lhet* | *lhetim* | -2864 | -5487 | 0.26 | 1368 | 0.0* | *lhet* |
| 2-3 | *lhet* | *lhetim* | -1759 | -4514 | 0.25 | 1385 | 0.0* | *lhet* |
| 2-4 | *lhet* | *lhetim* | -1940 | -4430 | 0.15 | 767 | 0.0* | *lhet* |
| 3-4 | *lhet* | *lhetim* | -1757 | -4340 | 0.06 | 319 | 0.0* | *lhet* |
| 1-*Amil* | *lhet* | *lhetsc* | -3754 | -4480 | 0.47 | 680 | 0.0* | *lhet* |
| 2-*Amil* | *lhet* | *lhetsc* | -3665 | -4439 | 0.14 | 221 | 0.0* | *lhet* |
| 2-*Amil* | *lhet* | *lhetim* | -3556 | -7790 | 0.40 | 3355 | 0.0* | *lhet* |
| 3-*Amil* | *lhet* | *lhetsc* | -3797 | -4707 | 0.04 | 70 | 0.0* | *lhet* |
| 4-*Amil* | *lhet* | *lhetsc* | -3547 | -4763 | 0.19 | 463 | 0.0* | *lhet* |

<sup>a</sup> *lhet*: one gene flow rate heterogeneous model with two time periods. Model parameters: *nref*, *t1*, *nu\_1*, *nu\_2*, *me1\_12*, *me1\_21*, *P<sub>E1</sub>*, *nu11*, *nu12*, *t2*, *me2\_12*, *me2\_21*, *P<sub>E2</sub>*, *nu21*, *nu22*
<sup>b</sup> *lhetim*: one gene flow rate heterogeneous model with one time period (isolation with migration). Model parameters: *nref*, *t1*, *nu\_1*, *nu\_2*, *me1\_12*, *me1\_21*, *P<sub>E1</sub>*, *nu11*, *nu12*
<sup>c</sup> *lhetsc*: one gene flow rate heterogeneous model with secondary contact. Model parameters: *nref*, *t1*, *nu\_1*, *nu\_2*, *nu11*, *nu12*, *t2*, *me2\_12*, *me2\_21*, *P<sub>E2</sub>*, *nu21*, *nu22*

